# Construction of two-input logic gates using Transcriptional Interference

**DOI:** 10.1101/724278

**Authors:** Antoni E. Bordoy, Nolan J. O’Connor, Anushree Chatterjee

## Abstract

Transcriptional Interference (TI) has been shown to regulate gene expression at the DNA level via different molecular mechanisms. The obstacles present on the DNA that a transcribing RNA Polymerase might encounter, e.g. a DNA-bound protein or another RNA Polymerase, can result in TI causing termination of transcription, thus reducing gene expression. However, the potential of TI as a new strategy to engineer complex gene expression modules has not been fully explored yet. Here we created a series of two-input devices using the presence of a roadblocking protein using both experimental and mathematical modeling approaches. We explore how multiple characteristics affect the response of genetic devices engineered to act like either AND, OR, or Single Input logic gates. We show that the dissociation constant of the roadblocking protein, inducer activation of promoter and operator sites, and distance between tandem promoters tune gate behavior. This work highlights the potential of rationally creating different types of genetic responses using the same transcription factors in subtly different genetic architectures.

## INTRODUCTION

Engineering bacteria to perform industrially and clinically useful tasks requires the implementation of sophisticated artificial gene regulation programs ^1^. The size and complexity of these programs has been shown to induce several design challenges, including varying construct performance in different hosts ^2–4^, the propagation of noise through cascading repressors ^5^, and cross-talk between genetic parts ^6,7^. Thus, in order to be able to engineer gene expression in an efficient and sophisticated manner, new genetic devices with minimal size, i.e a low DNA footprint, are required. One strategy towards this goal is reducing the DNA length needed to encode a certain response, i.e. minimizing the DNA footprint of a genetic device. Here we study how similar transcription factor recognition sequences with a similar DNA footprint can lead to diverse logic gate behaviors.

Transcription factors bind to specific DNA recognition sequences to regulate RNA polymerase (RNAP) activity by either recruiting it to promoter sites (activators) or blocking its binding to the DNA (repressors). Additionally, traffic of RNAPs can be controlled during transcription by the presence of “obstacles”, i.e. DNA-binding proteins and other RNAPs, usually causing the transcriptional process to prematurely end, decreasing gene expression. This second layer of regulation is a mode of Transcriptional Interference (TI) ^8^ and is present at different extents in a variety of organisms comprising the three domains of life ^9–13^. The presence of TI in a multitude of organisms and its potential to create higher-order gene regulation ^14,15^ has brought interest in its modeling ^15,16^ and engineered use as a tool to control gene expression ^17–20^.

Here we propose that TI can be used to obtain more complex gene regulation functions compared to gene expression driven by a single promoter. If the DNA to be transcribed is free of obstacles, transcription can proceed freely. However, if an obstacle to transcription is deliberately placed downstream of a promoter region, the RNAP traffic can be regulated by the controlled presence and absence of such an obstacle. In this transcriptional context, we refer as obstacle to: (i) a DNA binding protein in the same (sense) DNA strand from which transcription is taking place, and (ii) an RNAP initiating at or originated from a downstream sense promoter (Fig. 1a). For constitutive promoters, only the latter case occurs; however, inducible promoters can be understood as a conditionally activated combination of both obstacles. Since these obstacles can lead to TI, hereafter we refer to them as Transcriptional Interference Modules (TIMs).

**Figure 1.**
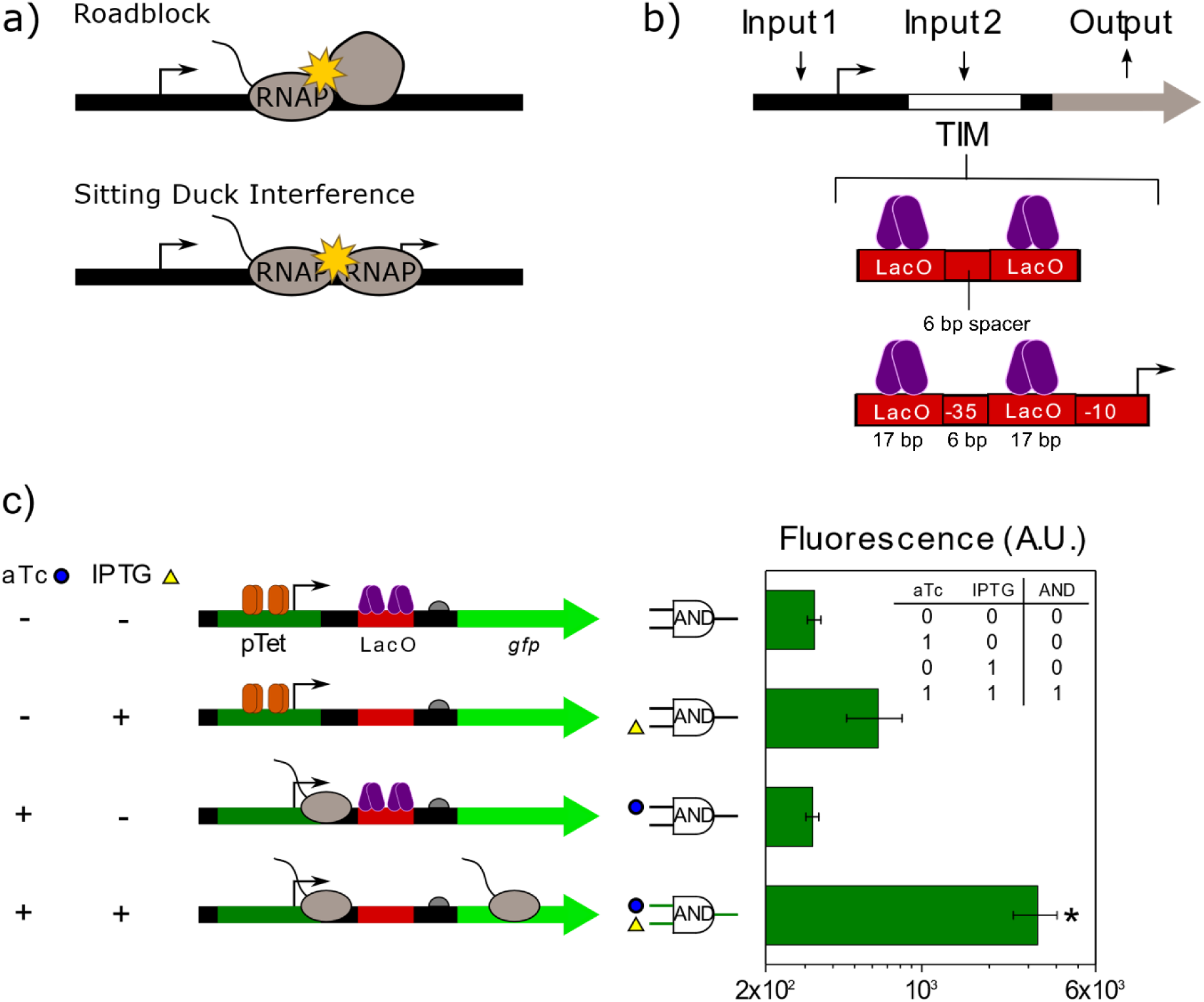
AND behavior can be created by placing a TIM composed of a strong roadblock downstream of an inducible promoter. **a)** Transcribing ECs, shown with an associated nascent RNA, can be intercepted by the presence of an obstacle bound to the downstream DNA. A transcription factor bound to the DNA causes roadblock, causing an elongating RNAP to stall and eventually terminate transcription. During sitting duck interference, an EC is prevented from transcribing further due to the presence of an initiating RNAP (sitting duck) at a downstream promoter. **b)** A transcriptional interference module (TIM) downstream from an inducible promoter can be used to engineering TI-based genetic devices. Shown are the two types of TIMs used in this study. **c)** Schematic showing how aTc and IPTG act as inputs of a genetic device designed to act as an AND logic gate the output of which is GFP. Gene expression is only highly activated when both inducers are present and is reduced to intermediate levels when only aTc is present due to the roadblock caused by bound LacI. The truth table for an AND gate is inserted in the plot. * indicates significant difference with the rest of conditions (Mann-Whitney *U* test, *p*-value<0.05).

Depending on which TIM the transcribing RNAP elongating complexes (ECs) encounter, different TI mechanisms may occur: (i) roadblock, in which the presence of a DNA bound protein either in the sense or antisense strand can impede the progression of ECs (Fig. 1a) ^21,22^; (ii) sitting duck interference, which is the unbinding of a promoter-bound RNAP, can also be caused by the movement of a tandem (Fig. 1a) or convergent EC ^23^; (iii) occlusion, which can be caused by an upstream tandem promoter or a downstream convergent promoter, is the process by which an RNAP is prevented to bind to a promoter due to the presence of an EC in that promoter region ^24–27^; and (iv) collision ^14,15,18,19,28,29^, occurring between two ECs moving in opposite directions, in which case either one or both ECs are susceptible to fall off the DNA ^30^. This study will focus on engineering the TI mechanisms of roadblock and sitting duck interference.

The combination of an inducible promoter with different downstream TIMs can lead to diverse gene expression patterns. Therefore, genetic devices with multiple inputs can be created to control the production of a protein, which is considered the output of the device (Fig. 1b). Here we focus on two-input logic gates. We show that a genetic device that has an architecture of an inducible promoter followed by a downstream roadblock site can perform AND logic, while a device with a similar architecture in which transcriptional activity is provided to the downstream TIM (thus transforming it into an inducible promoter) can exhibit OR logic. In this work, we present a series of two-input genetic devices designed with the aim of understanding and exploiting more complex ways of controlling RNAP traffic. We explore how the positioning and recognition sequence of the TIMs affect both gene expression and logic gate performance and develop mathematical models representing these constructs to validate and predict construct behavior. We demonstrate how the behavior of the genetic devices can be modified in a predictable manner by tuning biological parameters such as the dissociation constant of the roadblocking protein, activation of promoter and operator parts by their chemical inducers, and the distance between the two modules. Taken together, our results demonstrate the diverse gene expression profiles that are possible through rationally engineering simple genetic architectures.

## RESULTS

### Creation of AND logic using a downstream LacI roadblock site

We first anticipated that a TIM composed of a roadblocking protein downstream of an inducible promoter would behave as a two-input AND gate. Thus, we designed construct pAE_LG01 (Supplementary Table S1) consisting of a pTet promoter followed by the native LacI binding site (LacO) located 47 bp downstream. LacO is composed of two O1 sites separated by a 6 bp spacer sequence (Fig. 1b, Supplementary Table S1) ^31^. Transcriptional activity of pTet is controlled by the presence of aTc, which prevents the binding of repressor TetR to the DNA. The extent of successful transcription can then be further controlled by the magnitude of roadblock at the downstream TIM caused by the presence of the LacI repressor, which has been observed to greatly reduce transcription both *in vitro* and *in vivo* ^21,32^. Therefore, the transcriptional activity can also be controlled by the level of IPTG in the system, which binds to LacI and impedes its binding to the DNA. This construct is expected to minimally activate gene expression when only aTc is added while not responding to IPTG addition unless it is in combination with aTc (Supplementary Figures S11-S18), in which case the construct is expected to behave as an AND gate and produce high levels of GFP expression as its output only when both aTc and IPTG are present.

To express GFP, RNAPs need to be able to bind to pTet and freely transcribe through the roadblocking LacO site, i.e. without being roadblocked by LacI (Fig. 1c, bottom). Whereas if only aTc is available, transcription will be reduced by the presence of LacI (Fig. 1c, middle top). As expected, pAE_LG01 had a 10-fold increase in GFP expression only when both aTc and IPTG were added to the cells (Fig. 1c, bottom). However, in presence of aTc only, expression increased just 1.9-fold (Fig. 1c, middle top). Therefore, bound LacI caused a 5.2-fold decrease in GFP expression due to roadblock. This demonstrates that AND logic can be created by placing a roadblock site downstream of an inducible promoter.

### Point mutations in the LacO site tune the extent of roadblock repression caused by LacI, changing logic behavior

We hypothesized that in order to achieve good AND behavior, the roadblock interference needs to be strong, i.e. most RNAPs must not be able to read through it (Fig. 2a, top). Conversely, if the roadblock strength is low, RNAPs will read through it (Fig. 2a, bottom). To test this hypothesis, we created a library of constructs with one mutation in each of the O1 sites of LacO that modifies the dissociation constant, KD, of LacI (Fig. 2a, Supplementary Table S1) ^33^. LacI KD values ranged from 0.0092 to 2.34 pM. We measured the GFP expression of these constructs at the following four possible inducer combinations—no inducers, aTc only (50 ng/mL), IPTG only (1 mM), and aTc+IPTG. Expression was always the highest when both aTc and IPTG were present and lowest at the basal and IPTG only conditions (Fig. 2b). We observed increases in GFP expression ranging from 3.2- to 45-fold at the aTc+IPTG condition compared to the basal condition across the constructs. As anticipated, when cells were induced only with aTc, intermediate levels of gene expression ranging from 1.9- to 19-fold, with respect to basal, were observed (Fig. 2b).

**Figure 2.**
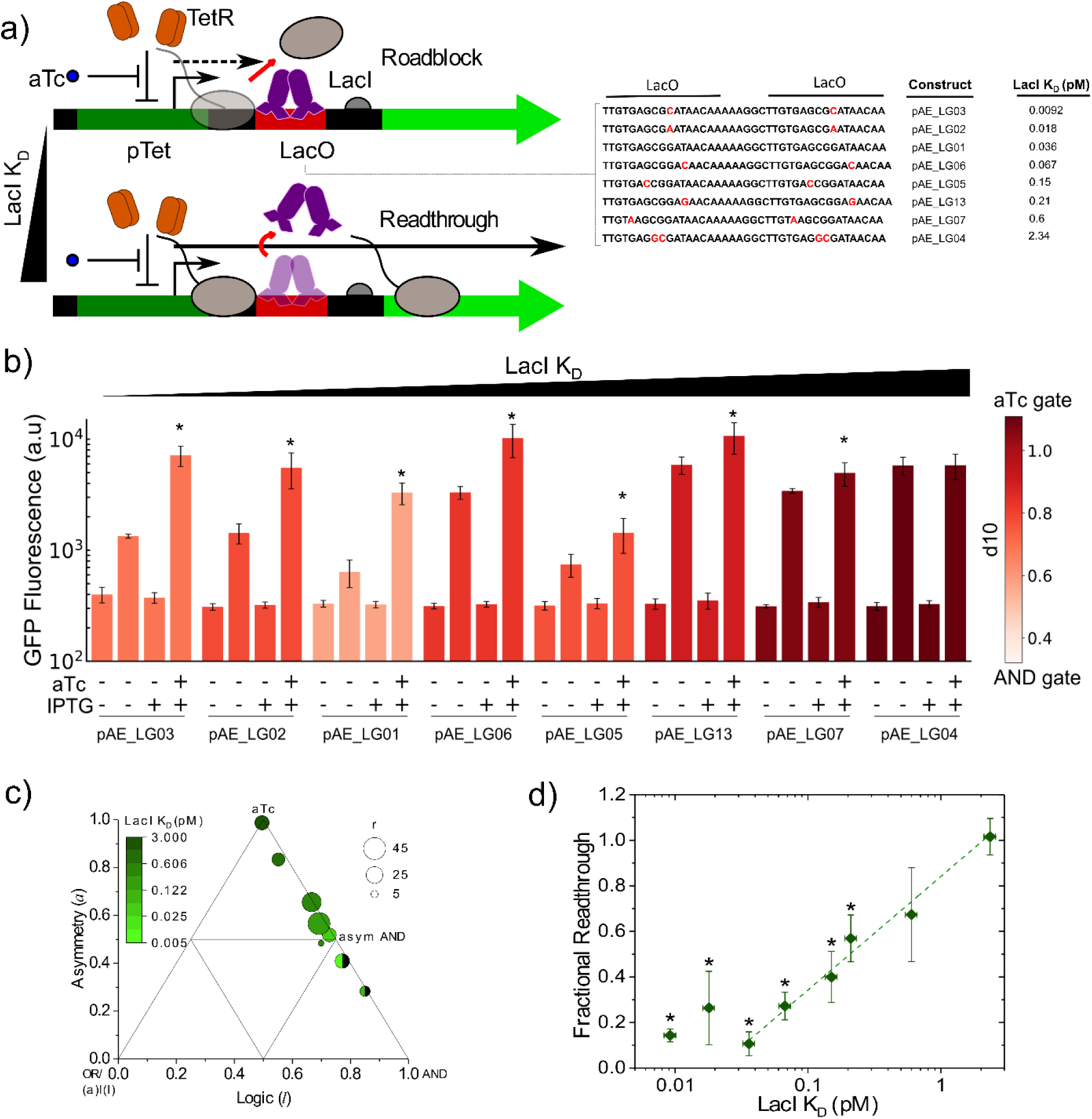
The switching from AND gate behavior to Single Input aTc Gate behavior as the dissociation constant of LacI increases. **a)** Mutations to the LacO site changes LacI dissociation constant, K_D_, and tunes the extent of RNAP readthrough. Small LacI dissociation constant values produce strong roadblock, which may lead to ECs falling off the DNA, stopping transcription (dashed arrow). With increasing LacI K_D_ values, LacI is dislodged in a greater extent from the DNA and transcription continues (plain arrow). **b)** GFP expression profiles for constructs with different LacI K_D_ at each of the four possible inducer combinations. Darker filling indicates a better AND behavior, measured by the value of *d*_*10*_. * indicates significant differences for the aTc+IPTG construct with respect to the other conditions (Mann-Whitney *U* test, *p*-value<0.05). **c)** Triangle plot showing the gate behaviors of AND constructs with different LacI K_D_ values. The plot shows dynamic range, *r;* the asymmetry, *a*, i.e. the responsiveness of the device to each input; and the logic type, *l*, i.e. whether one, two or three input combinations result in the output being ON. For a pure AND gate, *l*=1 and *a*=0. The designed constructs lie in the AND-aTc gate diagonal. Constructs were closer to the aTc gate vertex when LacI K_D_ was high, and closer to the AND vertex when LacI K_D_ was low. Full circles belong to the OR-AND-aTc space (GFP_a+I_> GFP_aTc_> GFP_IPTG_> GFP_Basal_). Half circles belong to the (a)I(I)-AND-aTc space (GFP_a+I_> GFP_aTc_> GFP_Basal_> GFP_IPTG_). **d)** Fractional readthrough is a function of LacI K_D_ for values >∼0.03 pM.

We then characterized the logic behavior of these constructs using a model previously developed by Cox *et al.* ^34^. The model quantifies the dynamic range in expression, *r;* the asymmetry, *a*, i.e. the relative responsiveness of the device to each input; and the logic type, *l*, i.e. whether one, two or three input combinations result in the output being ON. For the constructs presented here, we observed behaviors ranging from asymmetric AND gate to Single Input aTc Gate (Fig. 2b, c). We have also represented these results in Fig. 2b by color-coding the GFP expression profile of each construct with the calculated Euclidian distance from the logic (*l*) and asymmetry (*a*) values obtained for each construct to the perfect AND gate (*l*=1, *a*=0). This distance, *d*_*10*_, is dependent on *l* and *a*, and is a measure of the deviation from pure AND behavior (Equation 1, Materials and Methods), with higher values indicating a greater deviation from AND gate behavior. Graphically, this parameter *d*_*10*_ represents the distance of a particular gate from the bottom right vertex of a triangle plot, which corresponds to pure AND behavior (*l*=1, *a*=0) (Fig. 2c). In general, better AND behavior is obtained when LacI K_D_ is small, whereas the behavior tends to resemble an aTc gate (*l*=0.5, *a*=1) as LacI K_D_ becomes larger (Fig. 2b, c). This can also be observed by looking at the difference between the GFP expression levels at the aTc only condition and at the aTc+IPTG condition (Fig. 2b). This difference becomes virtually negligible for our control construct, pAE_LG04 (Supplementary Table S1), in which two mutations in each O_1_ sequence completely removed LacI binding. Therefore, pAE_LG04 behaves as a pure aTc gate.

The logic behavior of each construct is mainly determined by the intermediate levels of GFP expression when only aTc is present relative to the high GFP expression obtained when both aTc and IPTG are present. In other words, as we had hypothesized, the magnitude of the roadblock is the main factor dictating how well each AND construct behaves. If the roadblock caused by LacI is weak, then the ECs originating from pTet are able to dislodge LacI and continue transcription downstream, ultimately producing higher levels of GFP. To quantify the extent of successful transcription through the downstream roadblocking LacI site, we used Equation 3:

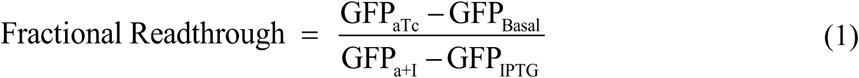

When plotted against LacI K_D_, we observed that the fractional readthrough follows an excellent logarithmic trend (Adj. R^2^=0.99) for K_D_ values greater than ∼0.03 pM (Fig. 2d). This data agrees with our suggested model for roadblock (Fig. 2a). For K_D_ values smaller than ∼0.03 pM, a plateau exists in which further decrease in K_D_ does not diminish readthrough any further. In this scenario, the unbinding frequency of LacI from the DNA is so small that the possibility for either an EC to escape roadblock due to momentarily LacI unbinding or for an EC to dislodge LacI is at its minimum. Therefore, most encounters between an EC and LacI result in the unbinding of the EC from the DNA (Fig. 2a, top, 2d). However, when LacI K_D_ values are high, the more frequent unbinding of LacI from the DNA increases the chances of a certain EC to escape the roadblock event before LacI rebinds to the DNA, while dislodging of LacI upon a clash with an incoming EC is also possible (Fig. 2a, bottom). Accordingly, six constructs that had lower K_D_ (pAE_LG01, 02, 03, 05, 06, 13) values had a significantly decreased GFP expression in presence of aTc and absence of IPTG compared to 1 mM IPTG (Mann-Whitney *U* test, *p*-value<0.05). The decrease in GFP expression caused by the presence of LacI ranged from 1.8- to 5.4-fold between constructs (Fig. 2b), with the closest AND gate behavior exhibited by the constructions with the lowest LacI K_D_. This is consistent with the fractional read through observed as K_D_ is increased (Fig. 2d). The change in K_D_ also impacted the regulatory range (*r*) between constructs, which tended to be higher at intermediate K_D_ values. These results show how roadblock can be engineered to downregulate gene expression in a predictable manner.

### AND behavior is improved through tuning inducer concentrations

Though dynamic range is typically maximized at saturating inducer concentrations, optimal AND behavior, quantified with *d*_*10*_, does not always occur at these conditions. We calculated *d*_*10*_ and the parameters that comprise it—asymmetry, *a*, and logic, *l*—at each set of inducer concentrations for our library of AND constructs (Materials and Methods, Supplementary Figures S3-S10). Low K_D_ (pAE_LG02) and high K_D_ (pAE_LG04) constructs experienced different inducer-dependent expression trends. We observed that, in the case of pAE_LG02, *d*_*10*_ generally decreased with high IPTG and low aTc concentrations (Fig. 3a-b). Higher pTet activation apparently permits readthrough of the LacI roadblock in the absence of IPTG, and therefore high aTc concentrations reduce AND-like behavior (increase *d*_*10*_). In the case of pAE_LG04, the LacI K_D_ is so high that there is no clear trend in *d*_*10*_ or other logic parameters with changing inducer concentration (Fig. 3a-b).

**Figure 3:**
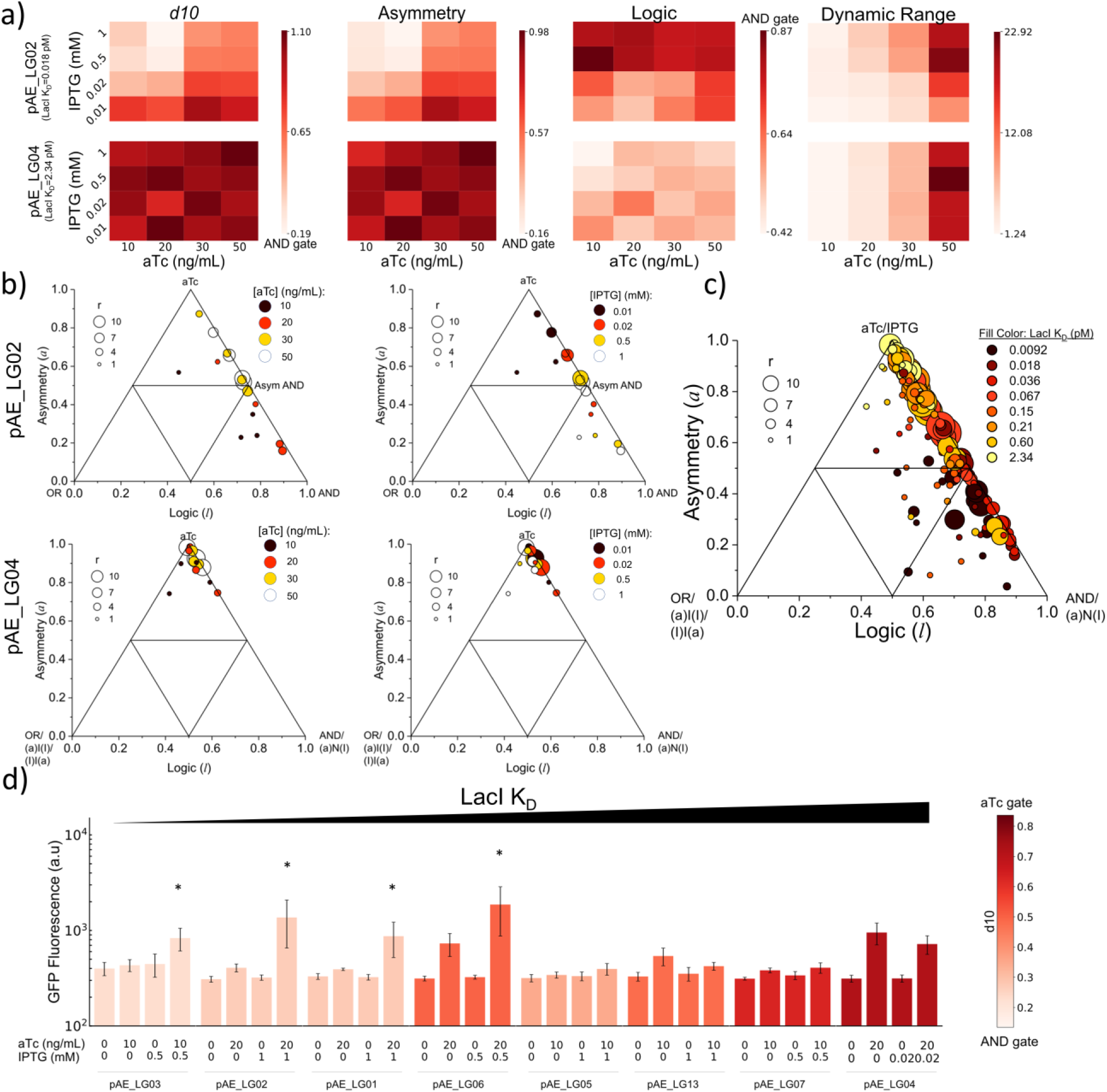
Optimization of AND gate behavior. **a)** Heat maps of logic gate analysis parameters for varying aTc and IPTG inducer concentrations for two AND constructs—pAE_LG02, which has a low K_D_ value and pAE_LG04, which has a high K_D_. **b)** Triangle plots for pAE_LG02 (top) and pAE_LG04 (bottom) showing trends in gate behavior with changing IPTG (left) and aTc (right). All data points resulting from combinations of four inducer levels belonged to the OR-AND-aTc space for pAE_LG02 while for pAE_LG04 they belonged to one of the following spaces: (a)I(I)-AND-aTc, OR-AND-aTc, OR-(a)N(I)-aTc or (I)I(a)-(a)N(I)-aTc ((a)N(I) means (aTc)NIMPLY(IPTG)), which share the aTc gate behavior at *l*=0.5 and *a*=1. **c)** Triangle plot for all 8 AND constructs at different aTc and IPTG combinations. The observed logic parameter spaces were a(I)I-AND-aTc, OR-AND-aTc, OR-AND-IPTG, OR-(a)N(I)-aTc or (I)I(a)-(a)N(I)-aTc. **d)** AND behaviors for each construct at conditions that minimize *d*_*10*_. * indicates significant differences in the aTc+IPTG condition with respect to the other inducer combinations shown. (Mann-Whitney *U* test, *p*-value<0.05)

The LacI K_D_ value of a construct, along with the aTc and IPTG concentrations, influence AND behavior (Supplementary Figures S3-S10). When all combinations of inducers for each construct are plotted, it is evident that low K_D_ constructs exhibit more AND-like behaviors at all concentrations of aTc and IPTG (Fig. 3c, Supplementary Figures S3-S10). When GFP expression of all the constructs at the concentrations of aTc and IPTG that minimize *d*_*10*_ are plotted, more AND-like behavior is apparent among the constructs with LacI K_D_ values below 0.21 pM (Fig. 3d). The GFP expression patterns at these *d*_*10*_-minimizing conditions offer low asymmetry—the construct responds to both inducers equally—and high logic—there is a clear ON state and three OFF states, corresponding to AND behavior (Fig. 3d). In the case of pAE_LG03, for example, this adjustment of inducer concentrations resulted in a >3-fold decrease in *d*_*10*_, trending toward ideal AND behavior.

Though each of these *d10*-minimizing conditions reduces the construct’s dynamic range compared with its expression profile at maximum aTc and IPTG concentrations (Fig. 2b, 3d), the improvement in AND behavior may in some conditions be more useful. For example, in applications where expression of the protein of interest needs to be tightly restricted in OFF conditions, true AND behavior may improve circuit performance. More broadly, this analysis shows that the activation of a tandem promoter and operator site are highly dependent on their relative strengths, and that small changes in the strength of each part significantly changes their performance in tandem.

### Mathematical modeling of RNAP roadblock and development of transfer functions

To gain mechanistic insights into the logic gate performance of our constructs, we developed mathematical models that predict GFP expression as a function of inducer concentrations (Fig 4a). We first used the Hill equation ^35^ to derive an inducer-dependent expression for the fraction of free transcription factor capable of binding to the promoter or operator sites. For example, the fraction of TetR with aTc bound is a function of the equilibrium dissociation constant for aTc to TetR, *K*_d, aTc:TetR_, the concentration of aTc, [*aTc*], and the Hill coefficient, *m*, which was either fitted or set to a value of 2, corresponding to the 2 molecules of aTc shown to substantially repress TetR:DNA binding ^36^.

**Figure 4.**
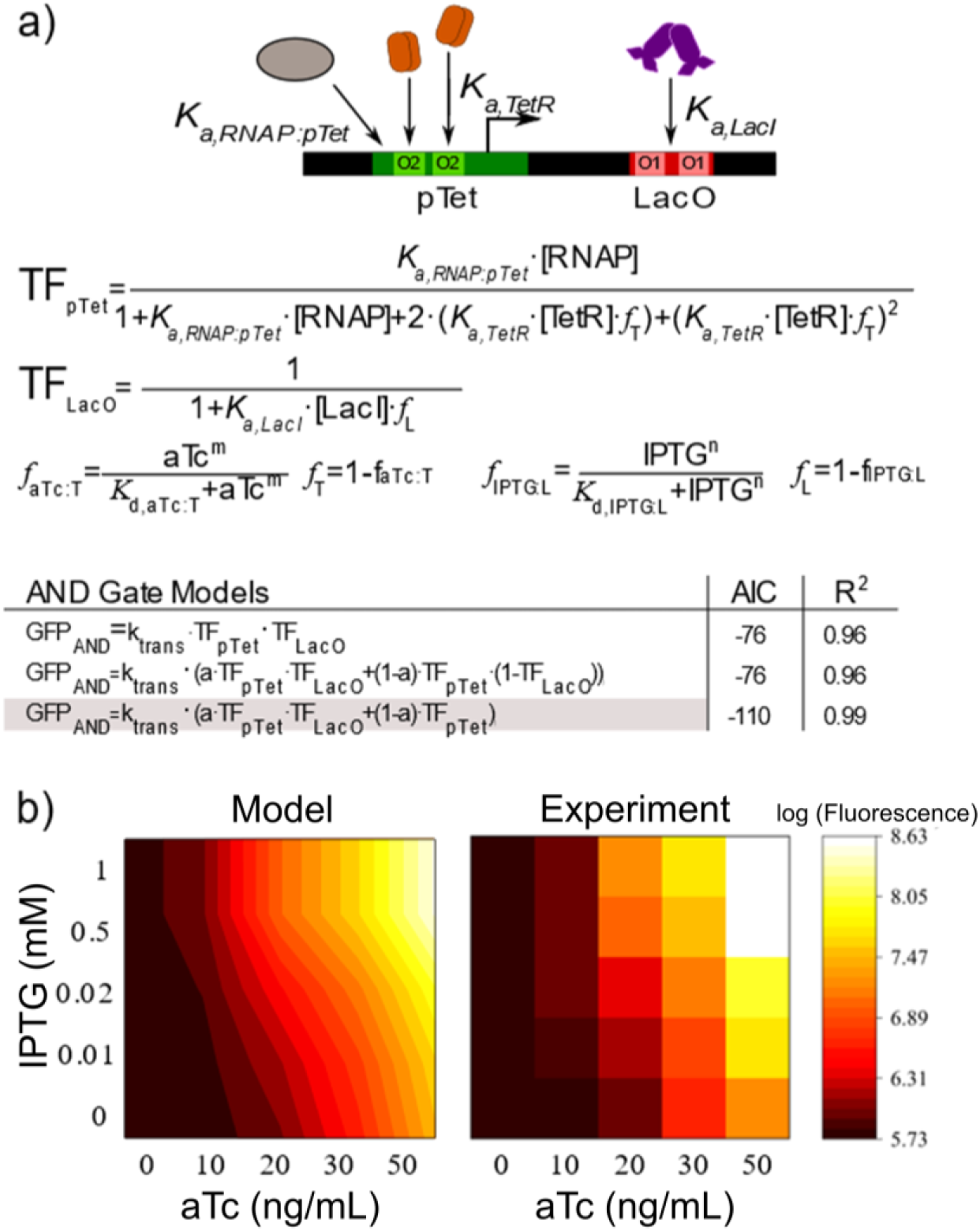
Mathematical modeling of promoter and operator occupancies captures AND behavior. Schematic showing the competing binding interactions at the pTet promoter and LacO operator site, with specific association constants for each protein. Transfer functions describing the occupancy of the promoter or operator are derived using the Shea-Ackers formalism—partitioning binding events that allow transcription in the numerator and all possible states in the denominator. These parameters are assembled into model equations describing the observed gate behaviors and quantified using the Akaike Information Criterion (AIC), a model selection criterion that penalizes spurious parameters in the model equations, with low AIC values indicating a better fit. We find find that, for pAE_LG02, AND + Single Input aTc gate behavior yields the best fit, with an R^2^ of 0.99. **b)** Log-transformed GFP expression data from 5 different concentrations of aTc and IPTG was used to fit the model equations.

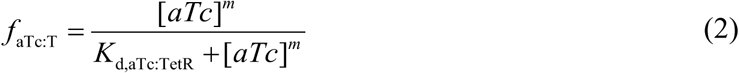

Thus, the fraction of free TetR capable of binding to TetO was estimated as:

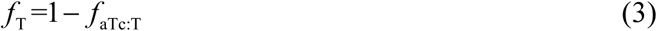

We then used the Shea-Ackers formalism ^37^ to derive transfer functions describing the occupancy of promoter and operator sites (*TF*_pTet_, *TF*_LacO_), with binding events that permit transcription in the numerator and all possible states in the denominator (Fig. 4a). For example, *TF*_pTet_ includes RNAP binding to the promoter (*K*_a,RNAP_ × [*RNAP*]) to initiate transcription in the numerator, and all other possible states—including TetR binding to one or both TetO sites on the pTet promoter—in the denominator:

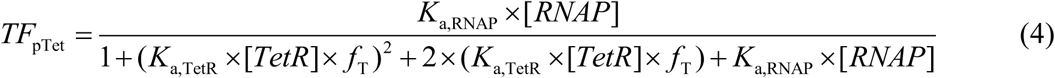

The resulting transfer functions were combined to develop model equations (see Supplementary section AND Gate Model Equation Derivations) that were fit to GFP expression data for varying aTc and IPTG concentrations for each of the constructs described (Fig. 4a). The constants and fitted biophysical parameters are reported in Supplementary Table 22 and Supplementary Figures S11-S18, respectively, and were compared against literature values whenever possible (see Supplementary section Mathematical Model Derivations).

After comparing several model equations, we found that model fits were improved through the addition of terms to our model equations that describe the effects of TI—namely the relationship between the LacO association strength and transcriptional roadblock. We quantitatively compared models using the Akaike Information Criterion (AIC) ^38^, a model selection criterion that compares the goodness of each model’s fit with respect to the number of terms in the model equation, with lower AIC values indicating a better model (Fig. 4a, Supplementary Table S24). Our three best performing models are shown in Fig. 4a. We found that our AND gate behavior was best captured using a model equation that consists of both an AND term (TF_pTet_·TF_pLac_) and a single-input gate term (TF_pTet_), as this function better describes the observed transcriptional readthrough than a pure AND function (Fig. 4a). The model was able to predict the fold change in GFP for the entire range of aTc and IPTG concentration variations as shown by the heat maps for construct pAE_LG02 with R^2^ value of 0.99 (Fig. 4b). The model also fit our other AND constructs well, with R^2^ values ranging from 0.97-1 (Figures S11-S18). The predictive ability of this simple mathematical model demonstrates how small sequence modifications can reliably and significantly change AND gate behavior.

### Addition of transcriptional activity associated with a weak roadblock to the downstream TIM creates OR logic

Transcriptional factor regulation can also result in other Boolean behaviors. For example, adding transcriptional activity to the LacO site, converting it to pLac promoter, will result in tandem transcription from both the upstream pTet and the downstream pLac. Tandem transcription has been previously used to create OR logic ^39,40^; however, the design specifications enable and optimize such behavior have not yet been investigated in depth. The output of an OR gate is ON when either one or both its inputs are ON, thus being only OFF when both inputs are OFF. Here we show that OR logic is only achieved when the roadblock created by the downstream inducible promoter is reduced. In other words, we show that any hypothetical pair of tandem promoters can potentially result in OR logic by engineering the extent of roadblock by tuning the downstream TF dissociation constant.

First, we created construct pAE_LG15, which is characterized by a pTet-pLac separation of 47 bp and a LacI K_D_=0.036 pM (Fig. 5a). This construct’s behavior demonstrated that providing the downstream TIM with transcriptional activity is not sufficient to create OR behavior (Fig. 5b, pAE_LG15). We use the Euclidean distance parameter *d*_*00*_ as a metric for OR behavior, since pure OR behavior is defined by *l*=0, *a*=0. Our results show that tandem transcription in this construct did not result in OR logic. Rather, the observed behavior for pAE_LG15, with an associated *d*_*00*_=0.76, was a single input gate (SIG) responsive to IPTG, the inducer of the downstream promoter, pLac (Fig. 5a and b, left). For this construct, the dissociation constant of LacI is small; therefore, LacI has a dual role of repressing transcriptional activity of pLac by blocking binding of RNAP to it while also roadblocking the upstream ECs originating from pTet; the addition of IPTG causes the alleviation of both forms of repression (Fig. 5a, left). We then hypothesized that in order to achieve a more OR-like behavior, the readthrough at the downstream TIM had to be increased (Fig. 5a, right), i.e. the roadblock magnitude needed to be decreased. For the particular system presented here, this means a higher GFP expression upon the addition of aTc only.

**Figure 5.**
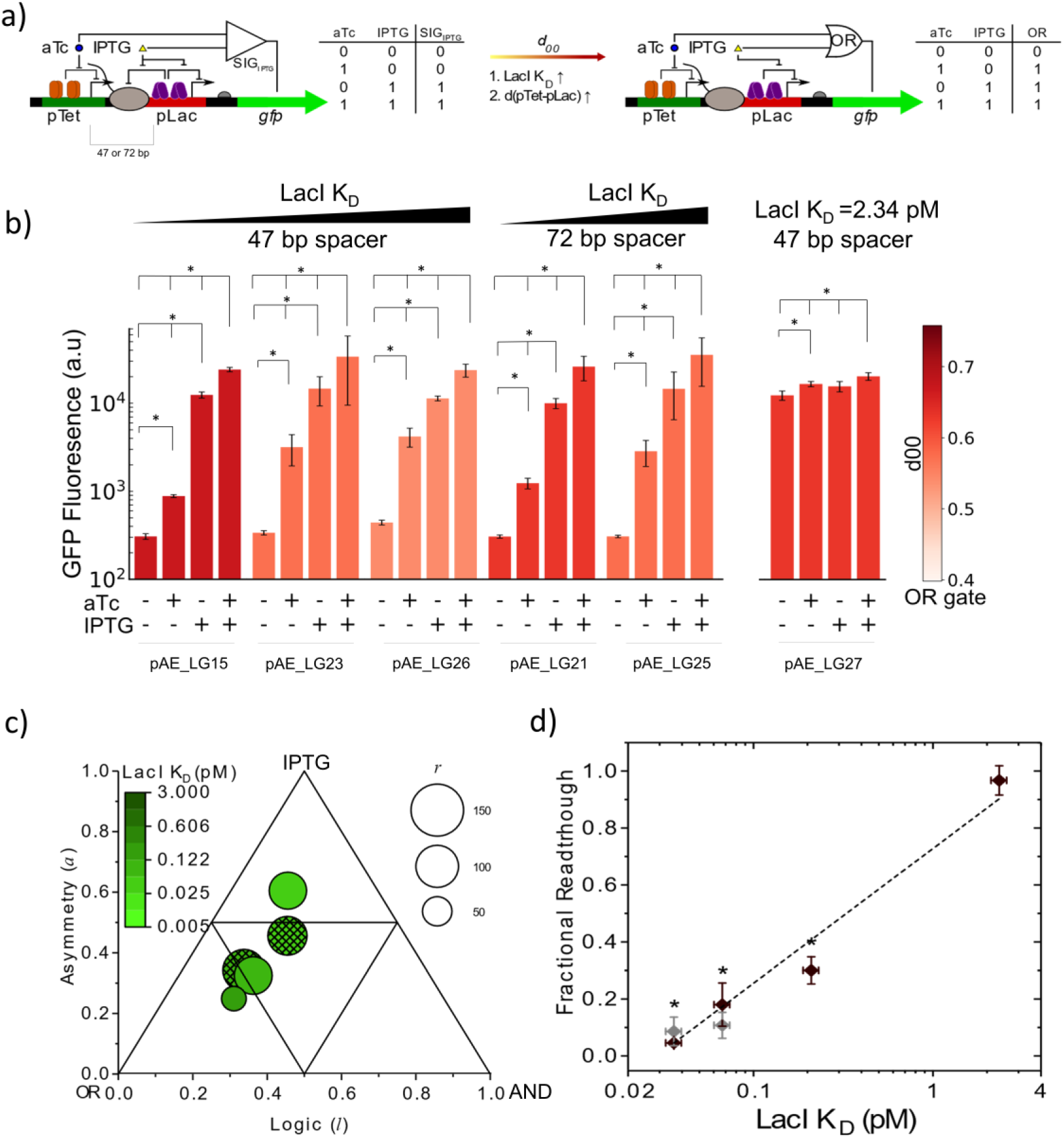
Tandem inducible promoters generate tunable OR logic behavior. **a)** If the roadblock caused by a downstream TIM formed by an inducible promoter is strong, then SIG behavior is obtained. Thus, the device only responds to the inducer of the downstream transcription factor. OR behavior can be improved by either increasing the dissociation constant of the roadblocking protein or by increasing the inter-promoter distance from 47 to 72 bp. **b)** Constructs with different LacI K_D_ values and inter-promoter distances (either 47 or 72 bp) exhibit varying logic behaviors. Modifications of the original TIM lead to higher GFP expression in the aTc only condition, improving OR behavior. * indicates significant differences in the aTc+IPTG condition with respect to the other inducer combinations shown. (Mann-Whitney *U* test, *p*-value<0.05). **c)** The triangle plot shows three metrics of gate behavior: dynamic range, *r;* the asymmetry, *a*, i.e. the responsiveness of the device to each input; and the logic type, *l.* OR gate is defined by, *l*=0 since three inputs should be able to turn gene expression ON, while all should turn it on to the same levels, i.e., *a*=0. Constructs lied close to a line parallel to the OR-IPTG gate axis, which indicates some AND gate component. OR behavior is improved with the increase of LacI K_D_, since allowing the upstream, aTc-induced pTet to readthrough the roadblock increases the contributions of the aTc input. Empty circles: 47 bp separation; patterned circles: 72 bp separation. Note: pAE_LG27 is not represented in this plot because it belongs to a separate logic gate parameter space (OR-AND-aTc). **d)** Fractional readthrough increases with LacI K_D_ independently of the pTet-pLac distance (brown: 47 bp, grey: 72 bp).

We took two different approaches to optimize OR behavior: (i) we increased the separation between pTet and pLac from 47 bp to 72 bp (Fig. 5a, middle), and (ii) we increased LacI K_D_ by introducing mutations in LacO. Specifically, we increased the separation between pTet and pLac from 47 bp to 72 bp in order to allow two stalled ECs, assuming an EC footprint of 35 bp, to sit in front of the roadblocking protein at the longest distance ^41^. This is based on previous reports that suggest that longer separations between the transcribing promoter and the roadblocking site can reduce the extent of roadblock due to RNAP cooperativity ^22^. The latter approach was taken to explore whether the results previously obtained for AND constructs would also hold true with this new architecture.

While keeping LacI K_D_ constant at 0.036 pM, we increased the separation between pTet and pLac, from 47 bp (pAE_LG15) to 72 bp (pAE_LG21). Only a slight improvement in OR behavior was observed (*d*_*00*_ decreased from 0.76 to 0.64, Supplementary Table S7) due to a significant 1.4-fold increase in GFP expression when only aTc was added to the cells (Fig. 5b, pAE_LG15 and pAE_LG21). The increased spacing likely allowed for an extra stalled EC in front of LacI—that could potentially induce RNAP cooperativity ^42^—only to increase readthrough 1.4-fold (Supplementary Figure S2). However, no significant differences were observed for the basal and aTc+IPTG conditions, suggesting the cause of the change in gene expression is a reduced LacI roadblock interference.

We next increased LacI K_D_ either ∼2- or ∼6-fold by mutating the LacO region within pLac for all the constructs (Fig. 2a). In the case of the 2-fold K_D_ increase, the GFP expression at the aTc only condition increased 9.6±3.5-fold and 10.0±3.3-fold with respect to the basal condition for 47 bp (Fig. 5b, pAE_LG23) and 72 bp (Fig. 5b, pAE_LG25) separation, respectively, and remained constant for the basal, IPTG, aTc+IPTG conditions. This improvement in OR behavior is reflected in lower *d*_*00*_ values for pAE_LG23 and pAE_LG25 (Supplementary Table S7) and suggests that increasing the K_D_ of the downstream roadblock to allow readthrough from the upstream promoter is necessary for more OR-like behavior. Interestingly, this increase in LacI K_D_ removed any effect of increasing interpromoter spacing—the *d*_*00*_ values of pAE_LG23 and pAE_LG25 are nearly identical (Fig. 5b; Supplementary Table S7), reflecting the similar GFP expression profiles across both interpromoter distances. This is an indication that the effect of RNAP cooperativity to facilitate dislodging the roadblock might only be effective when the dissociation constant of the roadblocking protein at the downstream TIM is small—that RNAP cooperation effects are only notable when the downstream roadblock is strong.

Further increasing LacI K_D_ to 0.21 pM for a pTet-pLac separation of 47 bp lead to a higher GFP expression in the basal and aTc only conditions, compared to pAE_LG23, while not significantly affecting the GFP levels at IPTG only or aTc+IPTG (Fig 5b, pAE_LG26). This increased basal expression is likely due to leaky expression at pLac. Since this construct’s gene expression was low only when both inducers were absent and high in the other three conditions, it closely resembles an OR gate. Accordingly, for this improved construct, *d*_*00*_ showed a reduction of ∼2-fold compared to our initial attempt to create OR behavior (Fig 5b and Supplementary Table S7, pAE_LG15 and pAE_LG26). The triangle plot containing the OR constructs similarly shows the trend towards pure OR gate behavior with increasing LacI K_D_ (Fig. 5c). In addition, the difference between the lowest ON state (aTc only) and the highest ON state (aTc+IPTG) was only 5.6±1.7-fold, which is smaller than the difference between the OFF state (basal) and the lowest ON state (aTc only).

When LacI K_D_ was increased to 2.34 pM, effectively abolishing the LacI roadblock, we observe a loss of OR behavior (Fig. 5b, pAE_LG27). This dramatic change in behavior can be attributed to the increase in leaky transcription in the absence of aTc and IPTG. Taken together, these results suggest that optimal OR behavior is achieved at moderate LacI K_D_ values that permit some readthrough from the upstream promoter but effectively block leaky transcription at the downstream TIM (Fig. 5b, 5c).

Though it is clear that increasing LacI K_D_ permits readthrough from the upstream pTet, it was not obvious that the trend in fractional readthrough would follow the one observed in our AND constructs (Fig. 2d). To address this, we also quantified the extent of readthrough for our OR gates using Equation 3. Intriguingly, a logarithmic correlation was also observed (Fig. 5d). The trend in fractional readthrough and LacI K_D_ was comparable to the one observed for the AND category (Supplementary Figure S2), suggesting that despite LacI’s dual purpose in roadblock and in RNAP occlusion, LacI K_D_ influences upstream RNAP readthrough in a manner similar to that of our AND constructs.

### Tuning inducer concentrations improves OR gate behavior

Just as the AND construct performance was sensitive to the relative strength of the promoter and operator parts and performed best (lowest *d*_*10*_) at sub-saturating aTc and IPTG conditions, the relative strength of the tandem promoters in our OR constructs can be tuned to improve OR gate performance. For constructs with moderately high LacI K_D_ values—pAE_LG26 (K_D_=0.21 pM), for instance—we find that OR gate performance is best (*d*_*00*_ is lowest) at high aTc and low IPTG concentrations (Fig. 6a). At these conditions, gate asymmetry is minimized since the upstream pTet requires high activation to read through the LacI roadblock. Thus, high aTc and low IPTG equalizes the relative GFP contributions from both promoters, creating more OR-like behavior. There is a strong trend in *d*_*00*_ with aTc concentration, since low pTet activity with a LacI roadblock produces a consistently low signal (Fig 6a). This trend is seen for all OR constructs (Supplementary Figures S19-S24) except pAE_LG27, which has a very high LacI K_D_ value (K_D_=2.34 pM) and does not respond to IPTG (Fig. 5d, 6a).

**Figure 6:**
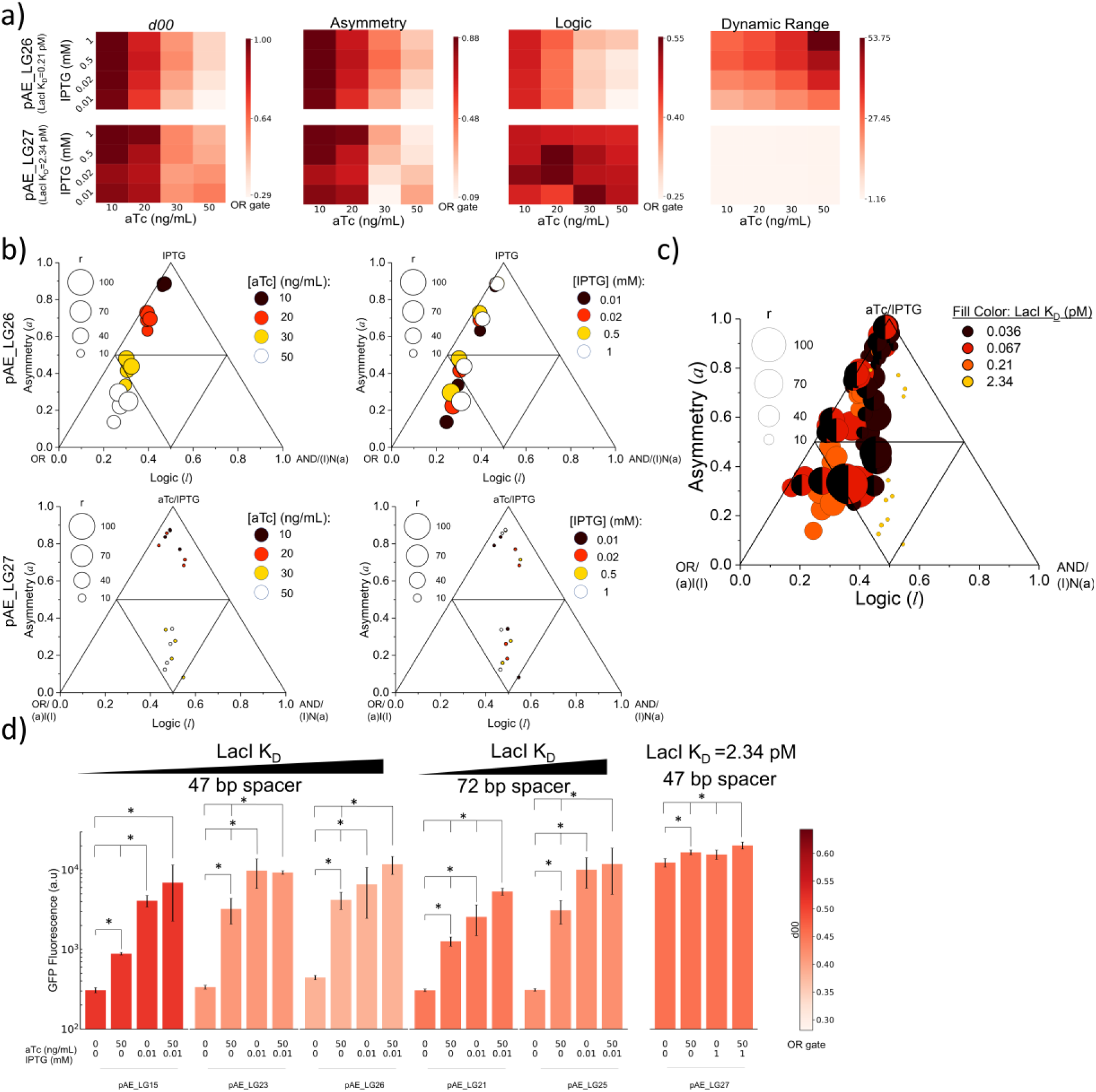
Optimization of OR gate behavior. **a)** Heat maps of logic gate analysis parameters for varying aTc and IPTG inducer concentrations for two OR constructs—pAE_LG26, which exhibited best OR behavior at saturating inducer concentrations and pAE_LG27, which has a high LacI K_D_. **b)** Triangle plots for pAE_LG26 (top) and pAE_LG27 (bottom) showing trends in gate behavior with changing IPTG (left) and aTc (right). For pAE_LG26, data points resulting from combinations of four inducer levels belonged to the OR-AND-IPTG space (GFP_a+I_> GFP_IPTG_> GFP_aTc_> GFP_Basal_) or the OR-(I)N(a)-IPTG space (GFP_IPTG_> GFP_a+I_> GFP_aTc_> GFP_Basal_). For pAE_LG27 data belonged to one of the following spaces: (a)I(I)-(I)N(a)-IPTG, OR-(I)N(a)-IPTG, OR-AND-IPTG, OR-AND-aTc or a(I)I-(I)N(a)-IPTG. Due to the high expression conditions of control construct pAE_LG27 at any inducer combination, its regulatory range, *r*, is very small and strictly its GFP expression profile is not always GFP_a+I_> GFP_aTc_> GFP_IPTG_> GFP_Basal_, thus resulting in this myriad of logic spaces. **c)** Triangle plot for all 6 OR constructs at different aTc and IPTG combinations. Half circles denote a 72 bp spacer. The observed logic parameter spaces excepting control construct pAE_LG27 were OR-AND-IPTG and OR-(I)N(a)-IPTG. **d)** OR behaviors for each construct at conditions that minimize *d*_*00*_. * indicates significant differences in the aTc+IPTG condition with respect to the other inducer combinations shown. (Mann-Whitney *U* test, *p*-value<0.05)

On a triangle plot of pAE_LG26 at varying aTc and IPTG conditions, OR constructs are clustered primarily by aTc concentration, again demonstrating the importance for high pTet strength in its upstream position (Fig. 6b). Within each cluster, low IPTG conditions trend toward more OR-like behavior, largely due to the equality of pTet and pLac strength at these conditions. Visualizing all conditions from all OR constructs on a single triangle plot shows that moderate LacI K_D_ values show more OR-like behavior, where low LacI K_D_ constructs trend toward IPTG single-input gate behavior (Fig. 6c).

Plotting OR behaviors for each construct at conditions that minimize *d*_*00*_, it is qualitatively apparent that OR behavior is improved at sub-saturating IPTG concentrations (Fig 6d). The best OR gate at saturating conditions, pAE_LG26, is improved with a reduction in *d*_*00*_ of 0.4 to 0.28 (Supplementary Table S7). Though these OR-optimal conditions reduce the dynamic range compared with saturating inducer conditions (Fig 6a), the emergence of more OR-like behavior may in some cases be more important than a large regulatory range. Thus, this optimization of gate behaviors should be considered alongside regulatory range when designing dual-input logic gates.

### Mathematical modeling of OR gate behavior

Modeling OR gate behavior suggests potential roadblock effects and RNAP interactions between tandem promoters. First, transfer functions describing promoter occupancy were derived similarly to the transfer functions describing promoter and operator occupancy for AND gates, though here *TF*_LacO_ is replaced with *TF*_pLac_ to represent the change in gate architecture (Fig. 7a, Supplementary Information section OR Gate Model Equation Derivations). Model equations used to fit OR gate behavior were derived considering the relative contributions of tandem promoters (Fig. 7a). Tamsir *et al.* had previously used a model equation accounting for the interference of upstream and downstream promoters (a_pTet_ and a_pLac_) with the maximum GFP expression from that promoter (X_pTet_ and X_pLac_) and the transfer functions describing promoter occupancy ^39^. We found that this model equation adequately described our OR gates— it provided the fit with the lowest AIC value—and fit inducer-dependent GFP expression with an R^2^ of 0.95 (Fig. 7b). R^2^ values for fits to other constructs range from 0.61-0.98 (Supplementary Figures S25-S30).

**Figure 7:**
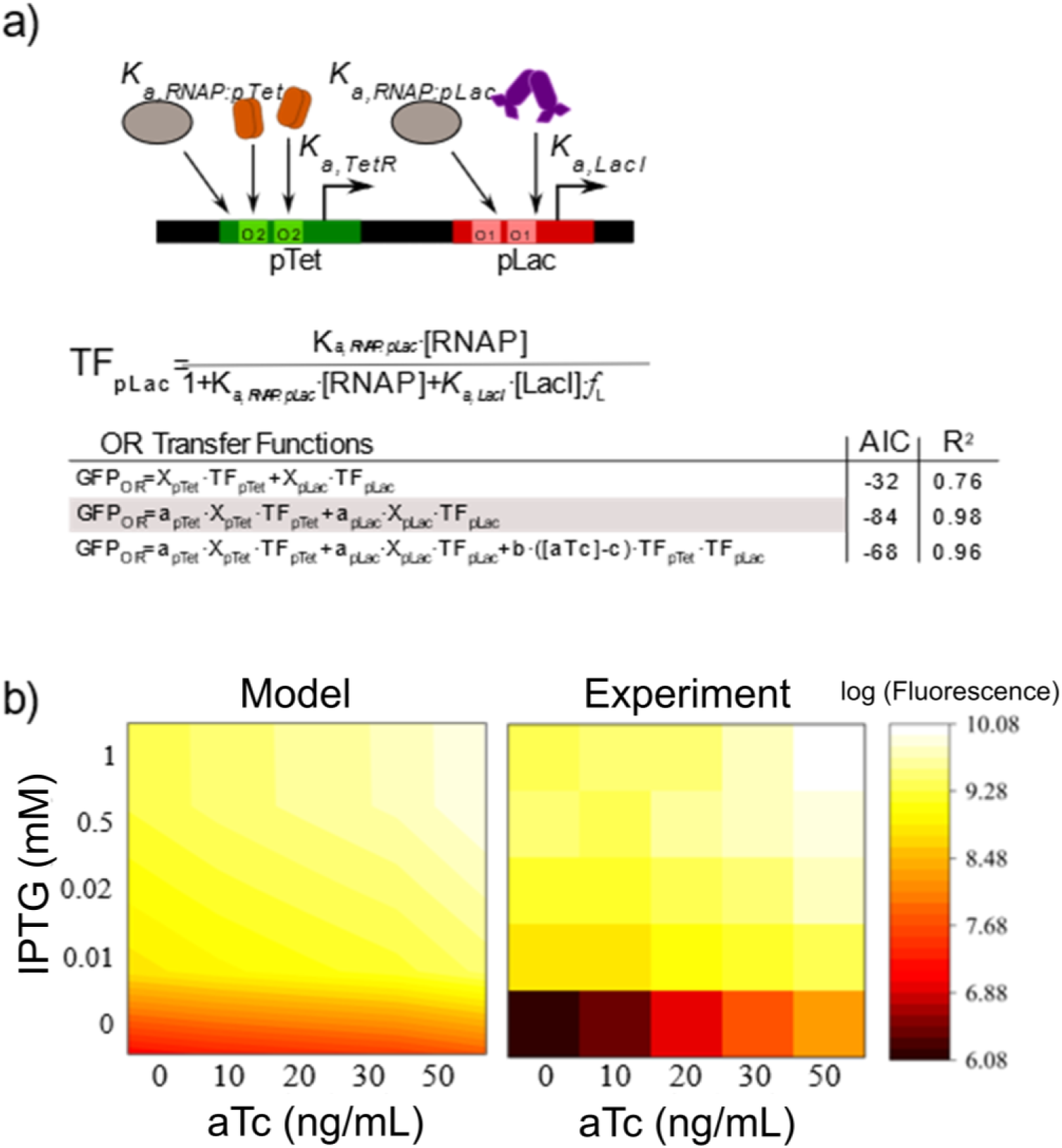
Mathematical models capture OR behavior. **a)** Schematic showing the competing binding interactions at the pTet and pLac promoters, with specific association constants for each binding protein. Transfer functions describing the occupancy of the promoters are derived using the Shea-Ackers formalism—partitioning binding events that allow transcription in the numerator and all possible states in the denominator. Note that all of the transfer functions and derivations are identical to that presented in the AND modeling in Figure 3, only the LacO transfer function, TF_LacO_ has been replaced with the pLac transfer function, TF_pLac_. These parameters are assembled into model equations describing the observed gate behaviors and quantified using the Akaike Information Criterion (AIC), a model selection criterion that penalizes spurious parameters in the model equations, with low AIC values indicating a better fit. For pAE_LG26, a model equation describing the additive contributions from tandem promoters with terms a_pTet_ and a_pLac_ to describe the promoters’ relative contributions to GFP expression provided the best fit, with an R^2^ of 0.95. **b)** Log-transformed GFP expression data for pAE_LG26 from 5 different concentrations of aTc and IPTG was used to fit the model equation with the lowest AIC value.

This model equation also revealed insights into potential interference and interactions between gates (Fig. 7a-b, see also Supplementary section OR Gate Modeling Derivation, Supplementary Table S23, and Supplementary Figures S25-S30). For example, the weight term describing the relative pLac contributions, a_pLac_, was in general significantly higher than a_pTet_, suggesting that the downstream pLac promoter interferes with the upstream pTet, either through RNAP interactions or through the LacI roadblock. The latter mode of interference may explain the trend in increasing a_pTet_ with increasing LacI K_D_ values (Supplementary Tables S2 and S23); the former may explain how the construct pAE_LG27, with a LacO K_D_ of 2.34 pM and high GFP expression under basal conditions (Fig. 5b), still has an a_pLac_ value over 3-fold higher than a_pTet_. Additionally, that these promoter weight values are below 1 suggests some level of interference between the tandem promoters, since the combined tandem promoter activities are not simply additive.

Here we have shown that OR behavior can be obtained by fine tuning the components of a pair of tandem promoters. Importantly, our results suggest that mutating the DNA recognition sequence of the transcriptional factor controlling the activity of the downstream promoter in a set of two tandem promoters is a more effective way to modulate TI and achieve OR behavior than increasing the inter-promoter distance.

## DISCUSSION

The presence of TI in naturally occurring systems has brought interest in the modeling and engineering of this regulatory phenomenon. Here, making use of two different TIMs downstream of an inducible pTet promoter we have been able to create AND and OR behaviors in a rational manner. Recently, Hao *et al*. showed how increasing LacI K_D_ strongly increased readthrough, doing so in a more effective manner than decreasing LacI concentration ^22^—an observation that is in agreement with our experiments— demonstrating that tuning LacI K_D_ is the most efficient manner to tune roadblock. In addition, our results regarding the different pTet-pLac separations also agree with the observations of Epshtein *et al.* that demonstrated how during a roadblock event the trailing EC helps the blocked complex to read through the roadblock site by keeping it in the active state. This is because once the blocked EC assumes its active configuration, it has a chance to move through the roadblock as soon as the latter dissociates ^21^. This mechanism could also explain the observation that as LacI K_D_ was increased in the AND and OR constructs, higher GFP expression was obtained when only aTc was added to the system because of the higher chances of escape of the ECs through the roadblock due to the more frequent unbinding events of the roadblocking protein. However, an alternative mechanism could be that stalled ECs actively dislodge the roadblocking protein and this action is increasingly favored as LacI K_D_ becomes larger. Thus, it remains unknown whether LacI dissociation and consequent readthrough of an EC occurs via a passive mechanism (ECs are just able to readthrough by waiting for spontaneous roadblock unbinding) or an active mechanism (ECs promote dislodgement of LacI), or a combination of both.

Using LacI and cAMP receptor protein (CRP) to control gene expression, Mayo *et al.* ^43^ showed how point mutations in the operator sites of each transcription factor changed the production of GFP. Their studies focused on experimentally demonstrating the plasticity of the input function of gene expression. A similar approach was used by Cox *et al.* ^34^ to construct a library of activation-repression and repression-repression promoters that ranged in their observed behavior from SIG to AND gates. Here, for the first time, we have been able to demonstrate this plasticity of the input function using rationally *de novo* engineered constructs by converting an initially AND gate to an aTc gate, and an IPTG gate into an OR gate (Fig. 2b; Fig. 5a, b). We show that rationally changing LacI K_D_ and inducer concentrations modulates TI and tunes AND and OR logic behaviors.

We have shown that both the position of operators of a certain transcription factor and the existence of point mutations in such operator sequences can affect the gene expression pattern of multi-input genetic devices. Our experimental observations indicate that diversification of transcription factor regulation is indeed readily achievable by DNA mutations or the insertion/deletion of small DNA fragments in the regulatory region. The binding of the same transcription factor to two slightly different sequences upstream of two different genes could result in disparate gene expression. This has important consequences on how we understand the design of synthetic genetic circuits in cells. Orthogonality between the new or existing parts of devices in a cell or its own cellular machinery is often considered essential for the good functioning of the synthetic device. However, our results indicate that a defined set of genetic elements can actually lead to various gene expression patterns, emphasizing that cells could use a certain transcription factor to obtain different responses depending on how it is arranged to other genetic elements and their relative strengths.

Moreover, the different degrees of readthrough observed at various LacI K_D_ values hint at how a downstream roadblock could be a mechanism utilized by microorganisms to fine-tune the expression of a gene under the transcription of either a constitutive promoter or an inducible promoter at a certain induction level. For example, increasing the number of transcription factors in the *cis*-regulatory region of a gene can increase its complexity in a small genetic space. In addition, the recent finding that promoters can rapidly evolve throughout the genome raises the prospect of interactions between neighboring promoters through pervasive transcription ^44^. Our results suggest that interplay between tandem RNAPs and RNAP interactions with protein roadblocks could allow nature to sample diverse gene expression profiles and tune as needed. Our work also highlights the ability of TI to control RNAP traffic to create and tune logic behaviors for synthetic biology while also exploring fundamental regulatory dynamics of RNAP-transcription factor and RNAP-RNAP interactions.

## ACKNOWLEDGEMENTS

The authors wish to acknowledge Basells Fellowship given to A.E.B., GAANN fellowship given to N.J.O. through the Department of Education, and the S10ODO21601 grant given to the Flow Cytometry Facility of the University of Colorado Boulder, and the National Science Foundation grant number MCB1714564 to A.C.

## AUTHOR CONTRIBUTIONS

A.E.B and A.C conceived of the study and designed the experiments. A.E.B designed the constructs and performed the experiments. N.J.O performed mathematical modeling. A.E.B, N.J.O, and A.C wrote the manuscript.

## MATERIALS AND METHODS

### Strains, Plasmids and cell culture

Constructs designed for AND behavior were cloned into pZE21MCS (Expressys). SalI and BamHI were used for the insertion of GFP, while the LacO operator site was inserted between KpnI and SalI, making the LacO sequence exchangeable for modified sequences with different LacI dissociation constants ^33^. Polymerase Chain Reaction (PCR) primers for inserting different LacO sites and pLac were purchased from Integrated DNA Technologies (IDT) and Life Technologies (Thermo Fisher). GFP was obtained from pAKgfp1 (Addgene #14076). For a list of inserted LacO sequences see Supplementary Table S1. The LacO fragment was then replaced with pLac containing different LacO sequences in order to create OR behavior (Supplementary Table S2). The separation between pTet and pLac was increased by the inserting DNA fragments of random sequence between EcoRI and KpnI (Supplementary Table S3).

Cloning and experiments to show logic behavior using TI with GFP were done in *E. coli* strain DH5*α*Z1 (Expressys). Transformation colonies were grown in Luria-Bertani (LB) and agar plates supplemented with kanamycin (50 μg/mL).

### GFP induction assays

Individual colonies were picked from LB and agar plates supplemented with 50 μg/mL kanamycin and incubated for 16 h at 37 °C under orbital shaking at 200 rpm. Then, the cells were diluted 1:10 into fresh LB media supplemented with 50 μg/mL kanamycin. Induction was performed at various inducer concentrations using anhydrous tetracycline (aTc), (0, 10, 20, 30 or 50 ng/mL) and isopropyl β-D-1-thiogalactopyranoside (IPTG), (0, 0.01, 0.02, 0.5 or 1 mM), creating a matrix of 25 different inducer combinations. Cells were grown for 6 h at 37 °C under shaking in a flat bottom 96-well plate in a microplate reader (Tecan Genios). Optical density at 590 nm was measured during induction. Following the growth period, the cells were transferred to a V-bottom 96-well plate and pelleted by centrifugation of the plate at 4000 rpm for 10 min at 4 °C. The supernatant was removed by vigorously inverting the plate and then the pellets were re-suspended in 100 μL PBS each. The centrifugation and supernatant removal processes were repeated and then each pellet was re-suspended in 100 μL PBS+4% formaldehyde and the plate was stored at 4 °C.

### Flow cytometry

Before fluorescence measurements conducted with a FACSCelesta instrument, samples were diluted 1:50 in PBS. The 588B 530/30V (800 V) channel was used to measure GFP levels. FSC-V=420 V, SSC-V=260 V, FSC-Threshold= 8000, SSC-Threshold= 200. For each sample, 50,000 cells were measured. At least four biological replicates were collected for each construct. Data was analyzed using MATLAB. Statistical differences were examined using the Mann-Whitney *U* test.

### Mathematical characterization of logic behavior

To measure the logic gate behavior of the engineered TI constructs, it was useful to characterize their GFP reporter expression using a previously developed mathematical model that classified the behavior of each construct into a certain type of ‘pure’ or ‘hybrid’/asymmetric logic gate ^34^. Such model, developed by Cox *et al.*, utilizes three parameters: (i) regulatory range, *r*, which measures the increase in gene expression using the ratio from the highest expressing condition compared to the lowest; (ii) logic, *l*, which quantifies whether the two intermediate expression levels are closer to the ON (*l*=1) or OFF state (*l*=0); and asymmetry, *a*, which quantifies the activation of gene expression caused by each inducer. Greater *a* values indicate that the device is more responsive to only one of the two inducers. Asymmetry varies between 0 and 1, with 0 indicating the gate responds equally to both, and 1 indicating the gate responds only to one. The parameters *a, l*, and *r* are defined mathematically in Equations S11-S13. We expanded the previously existing model by calculating the Euclidian distance, *d*_*la*_, between the *a* and *l* values observed for a certain construct (*a*_obs,_ *l*_obs_) and the *a* and *l* values corresponding to the desired behavior. The parameter *d*_*la*_ can range from 0 to 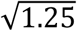. Pure AND behavior is characterized by *l*=1 and *a*=0. The deviation from AND behavior is thus represented by *d*_*10*_. In the case of an OR gate (*l*=0, *a*=0) the deviation is represented by *d*_*00*_.

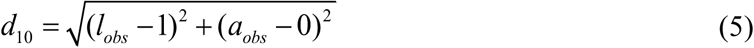

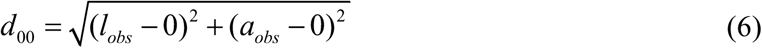

Experimental logic behavior is delimited to a certain parameter space defined by three pure logic gate behaviors. Constructs were assigned to their corresponding 3-gate parameter space, which is defined by their highest to lowest response at the four extreme conditions of basal, aTc only, IPTG only and aTc+IPTG. Each 3-gate parameter space can be represented in a triangular plot in which the base of the triangle corresponds to logic, *l*, and the height of the triangle corresponds to asymmetry, *a*. Since different types of logic gates can have the same *l* and *a* values, e.g. OR gate and AND gate each have *l*=0, *a*=0 because both have 3 states ON and one state OFF, then multiple parameter spaces exist, adding up to a total of 24 unique parameter spaces, e.g. 24 combinations of three pure logic gates, one with *l*=0, *a*=0, one with *l*=1, *a*=0 and one with *l*=0.5, *a*=0.5. For each parameter space, the behavior of a construct was classified into seven possible logic gate categories corresponding to the 3 pure gates at the corners of the triangle (*l*=0, *a*=0; *l*=0.5, *a*=1; *l*=1, *a*=0), 3 asymmetric gates (*l*=0.25, *a*=0.5; *l*=0.5, *a*=0.5; *l*=0.75, *a*=0.5) and the pure SLOPE gate (*l*=0.5, *a*=0) depending on which of these seven was closest to their observed behavior. The truth tables for the logic states defining these parameter spaces are reported in Supplementary Table S4; the possible logic parameter spaces resulting from each observed GFP expression profile is reported in Supplementary Table S5.

### Triangular AND gate plots

For constructs designed to behave as AND gates, i.e. pTet-LacO architecture, we observed that, when the induction levels were 50 ng/mL aTc and 1 mM IPTG, they belonged to either the OR-AND-aTc space (all but LG01 and LG03) or the (aTc)IMPLY(IPTG)-AND-aTc space (henceforth (a)I(I)-AND-aTc) (see Supplementary Table S4 for truth table for all gates described). The former is defined by the GFP expression levels being aTc+IPTG>aTc>IPTG>basal, while the latter is defined by the GFP expression levels being aTc+IPTG>aTc>basal>IPTG. This transition into different logic parameter spaces does not necessarily alter the logic behavior of a construct; different parameter spaces represent alternate possible deviations from the pure AND or OR logic behavior and highlight the versatile tunability achievable from two tandem regulatory parts (promoters or operators). (A more detailed explanation of this change is given in Supplemental section Assignment of Logic Parameter Space). These logic spaces are contiguous to one another and they share the AND-aTc diagonal ((*l*=0, *a*=1)-(*l*=0.5, *a*=1) diagonal) while their *l*=0, *a*=0 condition differs (either OR behavior or (aTc)IMPLY(IPTG)). The behaviors of the constructs fell either in the category of asymmetric AND gate (*l*=0.75, *a*=0.5) or aTc gate (*l*=0.5, *a*=1). With the best AND gate behavior observed for pAE_LG01 (*l*=0.85, *a*=0.28, *d*_*10*_=0.32), (Fig. 2b, c). The corresponding *d*_*10*_ values for all tested constructs with pTet-LacO architecture can be found in Supplementary Table S6.

### Triangular OR gate plots

For the OR gates (*l*=0, *a*=0), i.e. pTet-pLac architecture, when the induction levels were 50 ng/mL aTc and 1 mM IPTG, the constructs belonged to the OR-AND-IPTG space (GFP_aTc+IPTG_>GFP_IPTG_>GFP_aTc_>GFP_basal_) with the exception of control construct pAE_LG27 which belonged to the OR-AND-aTc space. The constructs of the OR-AND-IPTG space were categorized as either asymmetric SLOPE gate (*l*=0.5, *a*=0.5) or asymmetric OR gate (*l*=0.25, *a*=0.5). In this case, the construct with the best-performing OR logic was pAE_LG26 (l=0.31, *a*=0.25, *d*_*00*_=0.40), (Fig. 4b, c). The corresponding *d*_*00*_ values for all tested constructs with pTet-pLac architecture can be found in Supplementary Table S7.

This type of mathematical analysis is useful to determine the quality of the desired logic behavior, and it also helps demonstrate the plasticity of constructs that share a certain type of promoter architecture in the logic gates parameter space ^43^. To further demonstrate this plasticity, we applied the previously described analysis to all possible combinations of inducer conditions tested. Keeping the basal condition at 0 ng/mL aTc and 0 mM IPTG, we considered the “high” aTc condition to be either 10, 20, 30 or 50 ng/mL and the “high” IPTG condition to be either 0.01, 0.02, 0.5 or 1 mM. This results in 16 possible combinations of four “extreme” points. Sometimes this resulted in the tested conditions for a certain construct belonging to different logic parameter spaces (see Supplementary section Assignation of Logic Parameter Space, Supplementary Tables S8-S21).

### Transfer function modeling

Transfer functions were derived as described in Supplemental Information section Mathematical Model Derivation and Supplementary Figure S1 and assembled in model equations to fit AND and OR gate data. These transfer function derivations and model equations are defined in Equations S1-S10. Model equations were fit to experimental data using lsqcurvefit in MATLAB using a custom script. Several parameters—K_a,TetR:TetO_, K_a,IPTG:LacI_, [LacI], [TetR], [RNAP], and in some cases, the Hill coefficients m, n—were held constant to literature values (see Supplementary Table S22) in order to compare fitted parameters with experimentally observed values. Goodness of fit statistics from all best fits are available in Supplementary Figures S11-18, S25-S30. To compare model equations and prevent overfitting, we compared Akaike Information Criterion (AIC) values corresponding to each fit, which were calculated in MATLAB. AIC values for all fits are available in Supplementary Tables S24-S25.

## REFERENCES

(1) Brophy, J. A. N.; Voigt, C. A. Principles of Genetic Circuit Design. Nat. Methods 2014, 11 (5), 508–520. https://doi.org/10.1038/nmeth.2926.

(2) Cardinale, S.; Joachimiak, M. P.; Arkin, A. P. Report Effects of Genetic Variation on the E. Coli Host-Circuit Interface. CellReports 2013, 4 (2), 231–237. https://doi.org/10.1016/j.celrep.2013.06.023.

(3) Klumpp, S.; Zhang, Z.; Hwa, T. Theory Growth Rate-Dependent Global Effects on Gene Expression in Bacteria. Cell 2009, 139 (7), 1366–1375. https://doi.org/10.1016/j.cell.2009.12.001.

(4) Moser, F.; Broers, N. J.; Hartmans, S.; Tamsir, A.; Kerkman, R.; Roubos, J. A.; Bovenberg, R.; Voigt, C. A. Genetic Circuit Performance under Conditions Relevant for Industrial Bioreactors. 2012. https://doi.org/10.1021/sb3000832.

(5) Hooshangi, S.; Thiberge, S.; Weiss, R. Ultrasensitivity and Noise Propagation in a Synthetic Transcriptional Cascade. Proc. Natl. Acad. Sci. U. S. A. 2005, 102 (10), 3581–3586. https://doi.org/10.1073/pnas.0408507102.

(6) Moon, T. S.; Lou, C.; Tamsir, A.; Stanton, B. C.; Voigt, C. A. Genetic Programs Constructed from Layered Logic Gates in Single Cells. Nature 2012, 491 (7423), 249–253. https://doi.org/10.1038/nature11516.

(7) Stanton, B. C.; Nielsen, A. A. K.; Tamsir, A.; Clancy, K.; Peterson, T.; Voigt, C. A. Genomic Mining of Prokaryotic Repressors for Orthogonal Logic Gates. Nat. Chem. Biol. 2014, 10 (2), 99–105. https://doi.org/10.1038/nchembio.1411.

(8) Shearwin, K. E.; Callen, B. P.; Egan, J. B. Transcriptional Interference--a Crash Course. Trends Genet. 2005, 21 (6), 339–345. https://doi.org/10.1016/j.tig.2005.04.009.

(9) Wurtzel, O.; Sapra, R.; Chen, F.; Zhu, Y.; Simmons, B. A.; Sorek, R. A Single-Base Resolution Map of an Archaeal Transcriptome. Genome Res. 2010, 20 (1), 133–141. https://doi.org/10.1101/gr.100396.109.

(10) Dornenburg, J. E.; DeVita, A. M.; Palumbo, M. J.; Wade, J. T. Widespread Antisense Transcription in Escherichia Coli. MBio 2010, 1 (1), e00024-10–e00024-10. https://doi.org/10.1128/mBio.00024-10.

(11) Hongay, C. F.; Grisafi, P. L.; Galitski, T.; Fink, G. R. Antisense Transcription Controls Cell Fate in Saccharomyces Cerevisiae. Cell 2006, 127 (4), 735–745. https://doi.org/10.1016/j.cell.2006.09.038.

(12) Yelin, R.; Dahary, D.; Sorek, R.; Levanon, E. Y.; Goldstein, O.; Shoshan, A.; Diber, A.; Biton, S.; Tamir, Y.; Khosravi, R.; et al. Widespread Occurrence of Antisense Transcription in the Human Genome. Nat. Biotechnol. 2003, 21 (4), 379–386. https://doi.org/10.1038/nbt808.

(13) Katayama, S.; Tomaru, Y.; Kasukawa, T.; Waki, K.; Nakanishi, M.; Nakamura, M.; Nishida, H.; Yap, C. C.; Suzuki, M.; Kawai, J.; et al. Antisense Transcription in the Mammalian Transcriptome. Science (80-.). 2005, 309 (5740), 1564–1566. https://doi.org/10.1126/science.1112009.

(14) Chatterjee, A.; Johnson, C. M.; Shu, C.-C.; Kaznessis, Y. N.; Ramkrishna, D.; Dunny, G. M.; Hu, W.-S. Convergent Transcription Confers a Bistable Switch in Enterococcus Faecalis Conjugation. Proc. Natl. Acad. Sci. U. S. A. 2011, 108 (23), 9721–9726. https://doi.org/10.1073/pnas.1101569108.

(15) Bordoy, A. E.; Chatterjee, A. Cis-Antisense Transcription Gives Rise to Tunable Genetic Switch Behavior: A Mathematical Modeling Approach. PLoS One 2015, 10 (7), e0133873. https://doi.org/10.1371/journal.pone.0133873.

(16) Sneppen, K.; Dodd, I. B.; Shearwin, K. E.; Palmer, A. C.; Schubert, R. a; Callen, B. P.; Egan, J. B. A Mathematical Model for Transcriptional Interference by RNA Polymerase Traffic in Escherichia Coli. J. Mol. Biol. 2005, 346 (2), 399–409. https://doi.org/10.1016/j.jmb.2004.11.075.

(17) Brophy, J. A. N.; Voigt, C. A. Antisense Transcription as a Tool to Tune Gene Expression. 2016, 1–14.

(18) Bordoy, A. E.; Varanasi, U. S.; Courtney, C. M.; Chatterjee, A. Transcriptional Interference in Convergent Promoters as a Means for Tunable Gene Expression. ACS Synth. Biol. 2016, acssynbio.5b00223. https://doi.org/10.1021/acssynbio.5b00223.

(19) Hoffmann, S. A.; Kruse, S. M.; Arndt, K. M. Long-Range Transcriptional Interference in E. Coli Used to Construct a Dual Positive Selection System for Genetic Switches. Nucleic Acids Res. 2016, 44 (10), 1–12. https://doi.org/10.1093/nar/gkw125.

(20) Hoffmann, S. A.; Hao, N.; Shearwin, K. E.; Arndt, K. M. Characterizing Transcriptional Interference between Converging Genes in Bacteria. 2019. https://doi.org/10.1021/acssynbio.8b00477.

(21) Epshtein, V.; Toulmé, F.; Rahmouni, A. R.; Borukhov, S.; Nudler, E. Transcription through the Roadblocks: The Role of RNA Polymerase Cooperation. EMBO J. 2003, 22 (18), 4719–4727. https://doi.org/10.1093/emboj/cdg452.

(22) Hao, N.; Krishna, S.; Ahlgren-Berg, A.; Cutts, E. E.; Shearwin, K. E.; Dodd, I. B. Road Rules for Traffic on DNA—Systematic Analysis of Transcriptional Roadblocking in Vivo. Nucleic Acids Res. 2014, 42 (14), 8861–8872. https://doi.org/10.1093/nar/gku627.

(23) Callen, B. P.; Shearwin, K. E.; Egan, J. B. Transcriptional Interference between Convergent Promoters Caused by Elongation over the Promoter. Mol. Cell 2004, 14 (5), 647–656. https://doi.org/10.1016/j.molcel.2004.05.010.

(24) Adhya, S.; Gottesman, M. Promoter Occlusion: Transcription through a Promoter May Inhibit Its Activity. Cell 1982, 29 (July), 939–944. https://doi.org/10.1016/0092-8674(82)90456-1.

(25) Palmer, A. C.; Ahlgren-Berg, A.; Egan, J. B.; Dodd, I. B.; Shearwin, K. E. Potent Transcriptional Interference by Pausing of RNA Polymerases over a Downstream Promoter. Mol. Cell 2009, 34 (5), 545–555. https://doi.org/10.1016/j.molcel.2009.04.018.

(26) Greger, I. H.; Aranda, a; Proudfoot, N. Balancing Transcriptional Interference and Initiation on the GAL7 Promoter of Saccharomyces Cerevisiae. Proc. Natl. Acad. Sci. U. S. A. 2000, 97 (15), 8415–8420. https://doi.org/10.1073/pnas.140217697.

(27) Greger, I. H.; Demarchi, F.; Giacca, M.; Proudfoot, N. J. Transcriptional Interference Perturbs the Binding of Sp1 to the HIV-1 Promoter. Nucleic Acids Res. 1998, 26 (5), 1294–1300. https://doi.org/10.1093/nar/26.5.1294.

(28) Prescott, E. M.; Proudfoot, N. J. Transcriptional Collision between Convergent Genes in Budding Yeast. Proc. Natl. Acad. Sci. U. S. A. 2002, 99 (13), 8796–8801. https://doi.org/10.1073/pnas.132270899.

(29) Brophy, J. A. N.; Voigt, C. A. Antisense Transcription as a Tool to Tune Gene Expression Appendix Figures :

(30) Crampton, N.; Bonass, W. A.; Kirkham, J.; Rivetti, C.; Thomson, N. H. Collision Events between RNA Polymerases in Convergent Transcription Studied by Atomic Force Microscopy. Nucleic Acids Res. 2006, 34 (19), 5416–5425. https://doi.org/10.1093/nar/gkl668.

(31) Lutz, R.; Bujard, H. Independent and Tight Regulation of Transcriptional Units in Escherichia Coli via the LacR / O, the TetR / O and AraC / I 1 -I 2 Regulatory Elements. Nucleic Acids Res. 1997, 25 (6), 1203–1210.

(32) Deuschle, U.; Kammerer, W.; Gentz, R.; Bujard, H. Promoters of Escherichia Coli: A Hierarchy of in Vivo Strength Indicates Alternate Structures. EMBO J. 1986, 5 (11), 2987–2994.

(33) Betz, J. L.; Sasmor, H. M.; Buck, F.; Insley, M. Y.; Caruthers, M. H. Base Substitution Mutants of the Lac Operator: In Vivo and in Vitro Affinities for Lac Repressor. Gene 1986, 50 (1–3), 123–132. https://doi.org/10.1016/0378-1119(86)90317-3.

(34) Cox, R. S.; Surette, M. G.; Elowitz, M. B. Programming Gene Expression with Combinatorial Promoters. Mol. Syst. Biol. 2007, 3 (145), 145. https://doi.org/10.1038/msb4100187.

(35) Hill, A. V. The Possible Effects of the Aggregation of the Molecule of Hemoglobin on Its B Dissociation Curves. J. Physiol. 1910, 40, iv–vii. https://doi.org/10.1017/CBO9781107415324.004.

(36) Lederer, T.; Takahashi, M.; Hillen, W. Thermodynamic Analysis of Tetracycline-Mediated Induction of Tet Repressor by a Quantitative Methylation Protection Assay. Anal. Biochem. 1995, 232 (2), 190–196. https://doi.org/10.1006/abio.1995.0006.

(37) Shea, M. A.; Ackers, G. K. The OR Control System of Bacteriophage Lambda. A Physical-Chemical Model for Gene Regulation. J. Mol. Biol. 1985, 181 (2), 211–230. https://doi.org/10.1016/0022-2836(85)90086-5.

(38) Bozdogan, H. Model Selection and Akaike’s Information Criterion (AIC): The General Theory and Its Analytical Extensions. Psychometrika 1987, 52 (3), 345–370. https://doi.org/10.1007/BF02294361.

(39) Tamsir, A.; Tabor, J. J.; Voigt, C. A. Robust Multicellular Computing Using Genetically Encoded NOR Gates and Chemical ‘Wires.’ Nature 2011, 469 (7329), 212–215. https://doi.org/10.1038/nature09565.

(40) Stanton, B. C.; Nielsen, A. a K.; Tamsir, A.; Clancy, K.; Peterson, T.; Voigt, C. a. Genomic Mining of Prokaryotic Repressors for Orthogonal Logic Gates. Nat. Chem. Biol. 2014, 10 (2), 99–105. https://doi.org/10.1038/nchembio.1411.

(41) Krummel, B.; Chamberlin, M. J. Structural Analysis of Ternary Complexes of Escherichia Coli RNA Polymerase. Deoxyribonuclease I Footprinting of Defined Complexes. J. Mol. Biol. 1992, 225 (2), 239–250.

(42) Epshtein, V. Cooperation Between RNA Polymerase Molecules in Transcription Elongation. Science (80-.). 2003, 300 (5620), 801–805. https://doi.org/10.1126/science.1083219.

(43) Mayo, A. E.; Setty, Y.; Shavit, S.; Zaslaver, A.; Alon, U. Plasticity of the Cis-Regulatory Input Function of a Gene. PLoS Biol. 2006, 4 (4), e45. https://doi.org/10.1371/journal.pbio.0040045.

(44) Yona, A. H.; Alm, E. J.; Gore, J. Random Sequences Rapidly Evolve into de Novo Promoters. Nat. Commun. 2018, 9 (1), 1–10. https://doi.org/10.1038/s41467-018-04026-w.

