## Supplementary Information for "Construction of two-input logic gates using Transcriptional Interference"

### Mathematical Model Derivation

#### Transfer Functions for Promoter and Operator Occupancy

The equilibrium binding of a ligand to a transcription factor can be modeled through the Hill equation (Equation S1, same as Equation 4 in main text):

$$f_{aTc:T} = \frac{aTc^m}{K_d + aTc^m} \qquad \text{(S1)}$$

Where $f$ represents the fraction of transcription factor that is bound to the ligand, $m$ is the Hill coefficient, $aTc$ represents the ligand concentration—in this case, the concentration of aTc—and $K_d$ is the dissociation constant of the ligand to the transcription factor (Fig. 4 and 7, Fig. S1). The fraction of free transcription factor is then determined using Equation S2 (same as Equation 5 in main text):

$$f_T = 1 - f_{aTc:T} \qquad \text{(S2)}$$

The binding events, depicted in Fig. 4 for AND gate constructs and Fig. 7 for OR gate constructs, are represented in terms of the relevant protein concentrations and DNA dissociation constants. The model fitted values for the DNA dissociation constants are provided in Supplementary Table S22-S23 and in Supplementary Figures S11-S18, S25-S30 and literature-derived values are reported in Supplementary Table S23 in terms of the fraction of free protein that is not bound to a ligand. These states are described in Supplementary Figure S1, adapted from Tamsir *et al.*'s derivation (Tamsir *et al*, 2011) of transfer functions describing promoter states. GFP expression is a function of the inducer concentrations, which permits the observed logic gate behavior as a function of inducer concentrations.

We developed transfer function for pTet, pLac, and LacO utilizing the Shea-Ackers formalism (Shea & Ackers, 1985), a statistical mechanical approach used to describe DNA occupancy. The states describing the possible binding events for each part, depicted both in Figures 4a, 7a, and Supplementary Figure S1 for pTet, were arranged into transfer functions with states that permit transcription in the numerator and all possible states in the denominator.

In the case of LacO, shown in Equation S3a, the state that permits transcription is the lack of protein binding, denoted here with a 1, the binding repression term is shown in the denominator. The binding term is dependent on inducer through Equations S1 and S2, rendering this transfer function dependent on IPTG concentration. $TF_{pLac}$ results from the inclusion of RNAP binding, where instead transcription occurs with the RNAP:pLac binding term, $K_{a,RNAP:pLac} * [RNAP]$, in the numerator (Equation S3b). $TF_{pTet}$ contains a similar form to $TF_{pLac}$, but has additional terms capturing the binding dynamics of TetR to the two available TetO sites (Supplementary Figure S1).

$$\boldsymbol{TF_{LacO}} = \frac{1}{1 + (K_{a,LacI} * [LacI] * f_L)} \qquad \text{(S3a)}$$

$$\boldsymbol{TF_{pLac}} = \frac{K_{a,RNAP:pLac} * [RNAP]}{1 + (K_{a,LacI} * [LacI] * f_L) + K_{a,RNAP:pLac} * [RNAP]} \tag{S3b}$$

$$\boldsymbol{TF_{pTet}} = \frac{K_{a,RNAP} * [RNAP]}{1 + (K_{a,TetR} * [TetR] * f_T)^2 + 2 * (K_{a,TetR} * [TetR] * f_T) + K_{a,RNAP} * [RNAP]} \tag{S3c}$$

Transfer functions describing promoter and operator sites are then combined into model equations that attempt to describe the behavior of a logic gate system. These model equations were fit to grids of log-transformed flow cytometry data of GFP expression measured at 5 different aTc and 5 different IPTG concentrations and the goodness of fit statistics were recorded. Model equations were quantitatively compared using the Akaike Selection Criterion (Bozdogan, 1987), as reported in Supplementary Tables S24-25.

### AND Gate Model Equation Derivation

AND gates were first fitted using a model equation multiplying the two transfer functions, $TF_{pTet}$, $TF_{LacO}$, normalized through a parameter $k$, which accounts for translation. This equation is given as Equation S4:

$$GFP = k(TF_{pTet} * TF_{LacO}) \quad \text{(S4)}$$

However, this equation does not capture observed readthrough of RNAP through the LacI roadblock, which was observed in all AND constructs. Three models described by equations S5-S7, shown below, were developed in order to account for this readthrough.

$$GFP = k(a * TF_{pTet} * TF_{LacO} + (1 - a) * TF_{pTet} * (1 - TF_{LacO})) \quad (S5)$$

$$GFP = k(TF_{pTet} * TF_{LacO} + TF_{pTet}) \quad (S6)$$

$$GFP = k(a * TF_{pTet} * TF_{LacO} + (1 - a) * TF_{pTet}) \quad (S7)$$

Equation S5 effectively splits the GFP expression into true AND behavior for some fraction $a$, and RNAP from pTet reading through the LacI roadblock for some fraction (1-$a$), where $a$ is a fitted parameter.

Equation S6 models AND gate behavior as a combination of a true AND gate with a single aTc gate, and Equation 4 splits these contributions into fractions represented by a fitted parameter, $a$.

For the majority of constructs, Equation S7 provided the best fit for the number of parameters, as indicated by AIC and reported in Supplementary Table S24. Hill coefficients were either fit or fixed at a value of 2—the name 'Hill' next to the equation number in Supplementary Table S24 indicates that the coefficients were fixed. Parameter values and 95% confidence intervals are reported in Supplementary Figures S11-S18.

### OR Gate Model Equation Derivation

OR gates were fit using parameters from the AND gate with the corresponding operator site. For instance, in the case of pAE_LG25, the OR construct represented in the main text, all association constants except for $K_{LacI:pLac}$ Hill coefficients, and other coefficients unique to OR gate model equations, such as $a_{pTet}$, $a_{pLac}$, b, and c were obtained from the best fit of the AND gate containing the same operator site, pAE_LG06.

The tandem promoter OR constructs were initially modeled as a simple sum of the two promoter activities, shown in Equation S8:

$$GFP_{OR} = X_{pTet} * TF_{pTet} + X_{pLac} * TF_{pLac} \quad (S8)$$

Where $X_{pTet}$ and $X_{pLac}$ are the maximum promoter activities attributed to pTet and pLac, respectively. The downstream pLac activity, $X_{pLac}$, was obtained under the maximum IPTG and zero aTc condition; the upstream pTet activity, $X_{pTet}$, was obtained through mutations to the downstream pLac in both the -10, -35, and LacO regions that removed promoter activity and LacI binding activity.

Previously, Tamsir et al(Tamsir *et al*, 2011) proposed a model equation for OR gates that contains two terms describing the individual contributions of each tandem promoter with weighting terms $a_{pTet}$ and $a_{pLac}$ used to describe the interference of one promoter on another and the relative contributions of each promoter to GFP expression. This equation is represented in Equation S9.

$$GFP_{OR} = \boldsymbol{a}_{pTet} * X_{pTet} * TF_{pTet} + \boldsymbol{a}_{pLac} * X_{pLac} * TF_{pLac} \quad (S9)$$

These promoter weights suggest the relative importance of each promoter to GFP expression in the different OR constructs and are reported in Supplementary Table S23. As expected, the downstream pLac promoter is generally weighted more highly in all constructs, indicating that it contributes more to GFP expression in the presence of IPTG, which is in good agreement with experimental data shown in Figure 5b.

Through model selection using AIC, reported in Supplementary Table S25, we found that, for some constructs, such as pAE_LG25, this model equation performed best. But in some cases, the fits benefited from the addition of a third term, shown in the equation below:

$$GFP_{OR} = \boldsymbol{a}_{pTet} * X_{pTet} * TF_{pTet} + \boldsymbol{a}_{pLac} * X_{pLac} * TF_{pLac} + b * ([aTc] - c) * TF_{pTet} * TF_{pLac} \quad \text{(S10)}$$

Where $c$ is some critical value at which the promoters cooperate and acquire an AND-like behavior. This OR behavior is represented by the multiplication of the two promoters' transfer functions, and the term itself is weighted by coefficient $b$. In all fits in which this model equation was determined the best, the value of $b$ was relatively small, indicating that any possible cooperative effects between the two promoters are in some cases non-negligible.

### Assignment of Logic Parameter Space

In order to assign the parameter space to each data point, we followed the Cox *et. al* model(Cox *et al*, 2007). First the GFP expression was ordered from highest to lowest by inducer condition (none, aTc only, IPTG only, aTc+IPTG) and labelled as follows: IV>III>II>I. Even though the GFP expression from two different conditions might not be significantly different, we have strictly followed this rule in order to thoroughly and completely assign proper logic parameter space for each construct under each inducer conditions. Otherwise, calculations of asymmetry, *a*, and logic, *l*, might yield values outside of their theoretical range.

Each logic parameter space was then defined as a triangular space with vertices corresponding to the logic parameters, *a*, the asymmetry of the gate inputs aTc and IPTG, and *l*, the logic behavior that each gate exhibits—i.e which inputs are on vs. off under different inducer conditions. These parameters are defined in Equations S11 and S12 and are further described in the Methods section. Both Equations S11 and S12 contain the dynamic range term, *r*, which is defined in Equation S13.

$$a = \frac{log_{10}(III) - log_{10}(II)}{log_{10}(r)} \quad \text{(S11)}$$

$$l = \frac{2*log_{10}(IV) - log_{10}(II*III)}{2*log_{10}(r)} \quad \text{(S12)}$$

$$r = \frac{IV}{I} \quad \text{(S13)}$$

The vertices of the triangle plots are located at *a*=0, *l*=0*;* *a*=0, *l*=1; and *a*=1, *l*=0.5, which correspond to extreme or ideal logic behaviors. If three conditions are ON and one is OFF (IV=III=II>I) then *a*=0, *l*=0; if one condition is ON and three are OFF IV>(III=II=I) then *a*=0, *l*=1; finally, if two conditions are ON and two are OFF (IV=III)>(II=I) then *a*=1, *l*=0.5.

The logic space assigned to each construct/inducer combination reflects its GFP expression profile—the ranked order of all inducer conditions. For example, if GFP expression profile is $GFP_{a+I}$>$GFP_{aTc}$>$GFP_{IPTG}$>$GFP_{Basal}$, then the logic parameter space will be OR-AND-aTc, representing the *a*=0, *l*=0*;* *a*=0, *l*=1; and *a*=1, *l*=0.5 vertices, respectively. For this GFP expression profile, the *a*=0, *l*=0 vertex (IV=III=II>I) corresponds to OR logic, where only the uninduced, basal condition outputs a 0 while the presence of any input outputs a 1. The *a*=0, *l*=1 vertex represents AND behavior because, given the $GFP_{a+I}$>$GFP_{aTc}$>$GFP_{IPTG}$>$GFP_{Basal}$ expression profile, the IV>(III=II=I) state that defines the *a*=0, *l*=1 vertex represents ideal AND gate behavior characterized by only both inducers eliciting a 1 output. The state of the *a*=1, *l*=0.5, in this study, represents either a pure aTc or pure IPTG gate, since the (IV=III)>(II=I) condition indicates that the gate responds only to one inducer, aTc or IPTG.

Different GFP expression profiles will change the logic parameter space. For example, a GFP expression profile of $GFP_{a+I}$>$GFP_{aTc}$>$GFP_{Basal}$>$GFP_{IPTG}$ is assigned to an aTc(IMPLY)IPTG-AND-aTc logic space, because the (IV=III=II>I) state here defines an aTc-IMPLY-IPTG gate. The truth tables for all observed logic spaces are reported in Supplementary Table S4. All observed logic parameter spaces based on the GFP expression profile can be found in Supplementary Table S5.GFP expression data with the corresponding *a*, *l*, *r* (regulatory range), logic parameter space and logic behavior for each construct and combination of inducers can be found in Supplementary Tables S6-S21.

**Supplementary Table S1**: LacO sequences and their corresponding LacI $K_D$ values. WT: Wild Type. Each $O_1$ sequence is underlined. Mutations are highlighted in red and bold. $K_D$ values are taken from Betz *et al.*(Betz *et al*, 1986)

| Construct | LacO Sequence | $K_D$ (pM) |
|---|---|---|
| pAE_LG01 (WT) | TTGTGAGCGGATAACAAAAGGCTTGTGAGCGGATAACAA | 0.036 |
| pAE_LG02 | TTGTGAGCG**A**ATAACAAAAGGCTTGTGAGCG**A**ATAACAA | 0.018 |
| pAE_LG03 | TTGTGAGCG**C**ATAACAAAAGGCTTGTGAGCG**C**ATAACAA | 0.0092 |
| pAE_LG04 | TTGTGAG**GC**GATAACAAAAGGCTTGTGAG**GC**GATAACAA | 2.34 |
| pAE_LG05 | TTGTGA**C**CGGATAACAAAAGGCTTGTGA**C**CGGATAACAA | 0.15 |
| pAE_LG06 | TTGTGAGCGGA**C**AACAAAAGGCTTGTGAGCGGA**C**AACAA | 0.067 |
| pAE_LG07 | TT**A**TGAGCGGATAACAAAAGGCTT**A**TGAGCGGATAACAA | 0.6 |
| pAE_LG13 | TTGTGAGCGGA**G**AACAAAAGGCTTGTGAGCGGA**G**AACAA | 0.21 |

**Supplementary Table S2:** pLac sequences and their corresponding LacI $K_D$ values. WT: Wild Type. Each $O_1$ sequence is underlined. Mutations are highlighted in red and bold. The transcriptional start site is double underlined. $K_D$ values are taken from Betz *et al.*(Betz *et al*, 1986)

| Constructs | pLac Sequence | $K_D$ (pM) |
|---|---|---|
| pAE_LG15<br>pAE_LG21 | TTGTGAGCGGATAACAATTGACATTGTGAGCGGATAACAAGATA<br>CTGAGCACATCAGCAGGACGCACTGACT | 0.036 |
| pAE_LG23<br>pAE_LG25 | TTGTGAGCGGA**C**AACAATTGACATTGTGAGCGGA**C**AACAAGATA<br>CTGAGCACATCAGCAGGACGCACTGACT | 0.067 |
| pAE_LG26 | TTGTGAGCGGA**G**AACAATTGACATTGTGAGCGGA**G**AACAAGAT<br>ACTGAGCACATCAGCAGGACGCACTGACT | 0.21 |
| pAE_LG27 | TTGTGAG**GC**GATAACAATTGACATTGTGAG**GC**GATAACAAGATA<br>CTGAGCACATCAGCAGGACGCACTGACT | 2.34 |

**Supplementary Table S3:** Sequence content of the fragments of different lengths separating pTet and pLac.

| Constructs | pTet-pLac spacer | Length (bp) |
|---|---|---|
| pAE_LG15, pAE_LG23,<br>pAE_LG26, pAE_LG27 | ATCAGCAGGACGCACTGACCGAATTCATTAAAGAG<br>GAGAAAGGTACC | 47 |
| pAE_LG21, pAE_LG25 | ATCAGCAGGACGCACTGACCGAATTCGAATTGTGCG<br>ATCCCTGCACCTCAGCTAAGGTAGCTACCAGGTACC | 72 |

### Characterization of Logic Parameter Space

**Supplementary Table S4:** Truth tables of logic gates defining the logic parameter spaces spanning between AND and OR logic.

| | | 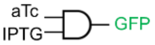 | 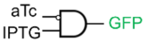 | 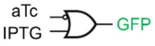 | 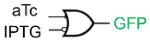 | 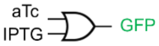 |
|---|---|---|---|---|---|---|
| aTc | IPTG | AND | (aTc) NIMPLY (IPTG) | (aTc) IMPLY (IPTG) | (IPTG) IMPLY (aTc) | OR |
| 0 | 0 | 0 | 0 | 1 | 1 | 0 |
| 1 | 0 | 0 | 0 | 0 | 1 | 1 |
| 0 | 1 | 0 | 1 | 1 | 0 | 1 |
| 1 | 1 | 1 | 0 | 1 | 1 | 1 |

**Supplementary Table S5:** Definitions of logic parameter spaces based on GFP expression profile.

| **GFP expression profile** | | | | **Extreme behaviors** | | | |
|---|---|---|---|---|---|---|---|
| **IV** | **III** | **II** | **I** | ***a*=0, *l*=0** | ***a*=0, *l*=1** | ***a*=1, *l*=0.5** | **Logic Space** |
| a+I | aTc | IPTG | Basal | OR | AND | aTc gate | OR-AND-aTc |
| a+I | aTc | Basal | IPTG | (aTc)IMPLY(IPTG) | AND | aTc gate | (a)I(I)-AND-aTc |
| aTc | a+I | IPTG | Basal | OR | (aTc)NIMPLY(IPTG) | aTc gate | OR-(a)N(I)-aTc |
| aTc | a+I | Basal | IPTG | (aTc)IMPLY(IPTG) | (aTc)NIMPLY(IPTG) | aTc gate | (a)I(I)-(a)N(I)-aTc |
| a+I | IPTG | aTc | Basal | OR | AND | IPTG gate | OR-AND-IPTG |
| IPTG | a+I | aTc | Basal | OR | (IPTG)NIMPLY(aTc) | IPTG gate | OR-(I)N(a)-IPTG |
| a+I | IPTG | Basal | aTc | (aTc)IMPLY(IPTG) | (IPTG)NIMPLY(aTc) | IPTG gate | (a)I(I)-(I)N(a)-IPTG |

**Supplementary Table S6:** Mathematical characterization of logic gate behavior using the expanded Cox *et al.* model for all tested constructs designed to behave as AND gates (pTet-LacO architecture).

| **Construct** | ***l*** | ***a*** | ***$d_{10}$*** | **Logic gate behavior** |
|---|---|---|---|---|
| pAE_LG01* | 0.85 | 0.28 | 0.32 | Asymmetric AND gate |
| pAE_LG02 | 0.73 | 0.52 | 0.59 | Asymmetric AND gate |
| pAE_LG03* | 0.77 | 0.41 | 0.47 | Asymmetric AND gate |
| pAE_LG04 | 0.50 | 0.99 | 1.11 | aTc gate |
| pAE_LG05 | 0.70 | 0.53 | 0.61 | Asymmetric AND gate |
| pAE_LG06 | 0.66 | 0.66 | 0.75 | Asymmetric AND gate |
| pAE_LG07 | 0.60 | 0.76 | 0.86 | aTc gate |
| pAE_LG13 | 0.58 | 0.81 | 0.91 | Asymmetric AND gate |

* belong to the (a)I(I)-AND-aTc space.

**Supplementary Table S7:** Mathematical characterization of logic gate behavior using the expanded Cox *et al.* model for all tested constructs designed to behave as OR gates (pTet-pLac architecture).

| **Construct** | ***l*** | ***a*** | $d_{00}$ | **Logic gate behavior** |
|---|---|---|---|---|
| pAE_LG15 | 0.46 | 0.61 | 0.76 | Asymmetric SLOPE gate |
| pAE_LG21 | 0.45 | 0.46 | 0.64 | Asymmetric SLOPE gate |
| pAE_LG23 | 0.34 | 0.34 | 0.48 | Asymmetric OR gate |
| pAE_LG25 | 0.36 | 0.32 | 0.49 | Asymmetric OR gate |
| pAE_LG26 | 0.31 | 0.25 | 0.40 | Asymmetric OR gate |
| pAE_LG27* | 0.46 | 0.12 | 0.48 | SLOPE gate |

* belongs to the OR-AND-aTc space.

**Supplementary Table S8:** GFP expression data with the corresponding *a*, *l*, *r*, logic parameter space and logic behavior for construct pAE_LG01. Basal always corresponds to aTc=0 ng/mL and IPTG=0 mM.

| **Inducer concentrations** | | **Condition** | | | | **Logic parameters** | | | | |
|---|---|---|---|---|---|---|---|---|---|---|
| **aTc (ng/mL)** | **IPTG (mM)** | **Basal** | **aTc** | **IPTG** | **a+I** | ***r*** | ***a*** | ***l*** | **Logic Space** | **Behavior** |
| 10 | 0.01 | 330 | 355 | 324 | 363 | 1.1 | 0.6 | 0.5 | (a)I(I)-AND-aTc | Asym SLOPE gate |
| 20 | 0.01 | 330 | 392 | 324 | 440 | 1.4 | 0.6 | 0.7 | (a)I(I)-AND-aTc | Asym AND gate |
| 30 | 0.01 | 330 | 484 | 324 | 631 | 1.9 | 0.6 | 0.7 | (a)I(I)-AND-aTc | Asym AND gate |
| 50 | 0.01 | 330 | 637 | 324 | 1216 | 3.8 | 0.5 | 0.7 | (a)I(I)-AND-aTc | Asym AND gate |
| 10 | 0.02 | 330 | 355 | 328 | 380 | 1.2 | 0.5 | 0.7 | (a)I(I)-AND-aTc | Asym AND gate |
| 20 | 0.02 | 330 | 392 | 328 | 484 | 1.5 | 0.4 | 0.8 | (a)I(I)-AND-aTc | Asym AND gate |
| 30 | 0.02 | 330 | 484 | 328 | 915 | 2.8 | 0.4 | 0.8 | (a)I(I)-AND-aTc | Asym AND gate |
| 50 | 0.02 | 330 | 637 | 328 | 2236 | 6.8 | 0.3 | 0.8 | (a)I(I)-AND-aTc | Asym AND gate |
| 10 | 0.5 | 330 | 355 | 332 | 438 | 1.3 | 0.2 | 0.9 | OR-AND-aTc | AND gate |
| 20 | 0.5 | 330 | 392 | 332 | 731 | 2.2 | 0.2 | 0.9 | OR-AND-aTc | AND gate |
| 30 | 0.5 | 330 | 484 | 332 | 1538 | 4.7 | 0.2 | 0.9 | OR-AND-aTc | AND gate |
| 50 | 0.5 | 330 | 637 | 332 | 3304 | 10.0 | 0.3 | 0.9 | OR-AND-aTc | Asym AND gate |
| 10 | 1 | 330 | 355 | 324 | 426 | 1.3 | 0.3 | 0.8 | (a)I(I)-AND-aTc | Asym AND gate |
| 20 | 1 | 330 | 392 | 324 | 872 | 2.7 | 0.2 | 0.9 | (a)I(I)-AND-aTc | AND gate |
| 30 | 1 | 330 | 484 | 324 | 1857 | 5.7 | 0.2 | 0.9 | (a)I(I)-AND-aTc | AND gate |
| 50 | 1 | 330 | 637 | 324 | 3299 | 10.2 | 0.3 | 0.9 | (a)I(I)-AND-aTc | Asym AND gate |

**Supplementary Table S9:** GFP expression data with the corresponding *a*, *l*, *r*, logic parameter space and logic behavior for construct pAE_LG02. Basal always corresponds to aTc=0 ng/mL and IPTG=0 mM.

| **Inducer concentrations** | | **Condition** | | | | **Logic parameters** | | | | |
|---|---|---|---|---|---|---|---|---|---|---|
| **aTc (ng/mL)** | **IPTG (mM)** | **Basal** | **aTc** | **IPTG** | **a+I** | ***r*** | ***a*** | ***l*** | **Logic Space** | **Behavior** |
| 10 | 0.01 | 309 | 338 | 318 | 344 | 1.1 | 0.6 | 0.4 | OR-AND-aTc | Asym SLOPE gate |
| 20 | 0.01 | 309 | 407 | 318 | 459 | 1.5 | 0.6 | 0.6 | OR-AND-aTc | Asym SLOPE gate |
| 30 | 0.01 | 309 | 804 | 318 | 894 | 2.9 | 0.9 | 0.5 | OR-AND-aTc | aTc gate |
| 50 | 0.01 | 309 | 1434 | 318 | 2152 | 7.0 | 0.8 | 0.6 | OR-AND-aTc | aTc gate |
| 10 | 0.02 | 309 | 338 | 313 | 385 | 1.2 | 0.3 | 0.8 | OR-AND-aTc | Asym AND gate |
| 20 | 0.02 | 309 | 407 | 313 | 593 | 1.9 | 0.4 | 0.8 | OR-AND-aTc | Asym AND gate |
| 30 | 0.02 | 309 | 804 | 313 | 1269 | 4.1 | 0.7 | 0.7 | OR-AND-aTc | Asym AND gate |
| 50 | 0.02 | 309 | 1434 | 313 | 3129 | 10.1 | 0.7 | 0.7 | OR-AND-aTc | Asym AND gate |
| 10 | 0.5 | 309 | 338 | 317 | 404 | 1.3 | 0.2 | 0.8 | OR-AND-aTc | Asym AND gate |
| 20 | 0.5 | 309 | 407 | 317 | 1112 | 3.6 | 0.2 | 0.9 | OR-AND-aTc | AND gate |
| 30 | 0.5 | 309 | 804 | 317 | 1788 | 5.8 | 0.5 | 0.7 | OR-AND-aTc | Asym AND gate |
| 50 | 0.5 | 309 | 1434 | 317 | 5109 | 16.5 | 0.5 | 0.7 | OR-AND-aTc | Asym AND gate |
| 10 | 1 | 309 | 338 | 321 | 387 | 1.3 | 0.2 | 0.7 | OR-AND-aTc | Asym AND gate |
| 20 | 1 | 309 | 407 | 321 | 1369 | 4.4 | 0.2 | 0.9 | OR-AND-aTc | AND gate |
| 30 | 1 | 309 | 804 | 321 | 2171 | 7.0 | 0.5 | 0.7 | OR-AND-aTc | Asym AND gate |
| 50 | 1 | 309 | 1434 | 321 | 5543 | 17.9 | 0.5 | 0.7 | OR-AND-aTc | Asym AND gate |

**Supplementary Table S10:** GFP expression data with the corresponding *a*, *l*, *r*, logic parameter space and logic behavior for construct pAE_LG03. Basal always corresponds to aTc=0 ng/mL and IPTG=0 mM.

| **Inducer concentrations** | | **Condition** | | | | **Logic parameters** | | | | |
|---|---|---|---|---|---|---|---|---|---|---|
| **aTc (ng/mL)** | **IPTG (mM)** | **Basal** | **aTc** | **IPTG** | **a+I** | ***r*** | ***a*** | ***l*** | **Logic Space** | **Behavior** |
| 10 | 0.01 | 399 | 433 | 471 | 535 | 1.3 | 0.3 | 0.6 | OR-AND-IPTG | Asym SLOPE gate |
| 20 | 0.01 | 399 | 656 | 471 | 792 | 2.0 | 0.5 | 0.5 | OR-AND-aTc | Asym SLOPE gate |
| 30 | 0.01 | 399 | 985 | 471 | 1613 | 4.0 | 0.5 | 0.6 | OR-AND-aTc | Asym SLOPE gate |
| 50 | 0.01 | 399 | 1339 | 471 | 3802 | 9.5 | 0.5 | 0.7 | OR-AND-aTc | Asym AND gate |
| 10 | 0.02 | 399 | 433 | 597 | 817 | 2.0 | 0.4 | 0.7 | OR-AND-IPTG | Asym AND gate |
| 20 | 0.02 | 399 | 656 | 597 | 1088 | 2.7 | 0.1 | 0.6 | OR-AND-aTc | SLOPE gate |
| 30 | 0.02 | 399 | 985 | 597 | 1820 | 4.6 | 0.3 | 0.6 | OR-AND-aTc | Asym SLOPE gate |
| 50 | 0.02 | 399 | 1339 | 597 | 5946 | 14.9 | 0.3 | 0.7 | OR-AND-aTc | Asym AND gate |
| 10 | 0.5 | 399 | 433 | 445 | 833 | 2.1 | 0.0 | 0.9 | OR-AND-IPTG | AND gate |
| 20 | 0.5 | 399 | 656 | 445 | 1897 | 4.8 | 0.2 | 0.8 | OR-AND-aTc | Asym AND gate |
| 30 | 0.5 | 399 | 985 | 445 | 3359 | 8.4 | 0.4 | 0.8 | OR-AND-aTc | Asym AND gate |
| 50 | 0.5 | 399 | 1339 | 445 | 8615 | 21.6 | 0.4 | 0.8 | OR-AND-aTc | Asym AND gate |
| 10 | 1 | 399 | 433 | 373 | 628 | 1.7 | 0.2 | 0.8 | (a)I(I)-AND-aTc | AND gate |
| 20 | 1 | 399 | 656 | 373 | 2034 | 5.5 | 0.3 | 0.8 | (a)I(I)-AND-aTc | Asym AND gate |
| 30 | 1 | 399 | 985 | 373 | 3413 | 9.2 | 0.4 | 0.8 | (a)I(I)-AND-aTc | Asym AND gate |
| 50 | 1 | 399 | 1339 | 373 | 7167 | 19.2 | 0.4 | 0.8 | (a)I(I)-AND-aTc | Asym AND gate |

**Supplementary Table S11:** GFP expression data with the corresponding *a*, *l*, *r*, logic parameter space and logic behavior for construct pAE_LG04. Basal always corresponds to aTc=0 ng/mL and IPTG=0 mM.

| **Inducer concentrations** | | **Condition** | | | | **Logic parameters** | | | | |
|---|---|---|---|---|---|---|---|---|---|---|
| **aTc (ng/mL)** | **IPTG (mM)** | **Basal** | **aTc** | **IPTG** | **a+I** | ***r*** | ***a*** | ***l*** | **Logic Space** | **Behavior** |
| 10 | 0.01 | 313 | 387 | 313 | 372 | 1.2 | 0.8 | 0.6 | (I)I(a)-(a)N(I)-aTc | aTc gate |
| 20 | 0.01 | 313 | 952 | 313 | 965 | 3.1 | 1.0 | 0.5 | (a)I(I)-AND-aTc | aTc gate |
| 30 | 0.01 | 313 | 2118 | 313 | 1692 | 6.8 | 0.9 | 0.6 | (I)I(a)-(a)N(I)-aTc | aTc gate |
| 50 | 0.01 | 313 | 5802 | 313 | 4719 | 18.5 | 0.9 | 0.5 | (I)I(a)-(a)N(I)-aTc | aTc gate |
| 10 | 0.02 | 313 | 387 | 314 | 381 | 1.2 | 0.9 | 0.5 | OR-(a)N(I)-aTc | aTc gate |
| 20 | 0.02 | 313 | 952 | 314 | 720 | 3.0 | 0.7 | 0.6 | OR-(a)N(I)-aTc | Asym SLOPE gate* |
| 30 | 0.02 | 313 | 2118 | 314 | 2259 | 7.2 | 1.0 | 0.5 | OR-AND-aTc | aTc gate |
| 50 | 0.02 | 313 | 5802 | 314 | 4082 | 18.5 | 0.9 | 0.6 | OR-(a)N(I)-aTc | aTc gate |
| 10 | 0.5 | 313 | 387 | 319 | 389 | 1.2 | 0.9 | 0.5 | OR-AND-aTc | aTc gate |
| 20 | 0.5 | 313 | 952 | 319 | 933 | 3.0 | 1.0 | 0.5 | OR-(a)N(I)-aTc | aTc gate |
| 30 | 0.5 | 313 | 2118 | 319 | 1765 | 6.8 | 0.9 | 0.5 | OR-(a)N(I)-aTc | aTc gate |
| 50 | 0.5 | 313 | 5802 | 319 | 7184 | 22.9 | 0.9 | 0.5 | OR-AND-aTc | aTc gate |
| 10 | 1 | 313 | 387 | 328 | 384 | 1.2 | 0.7 | 0.4 | OR-(a)N(I)-aTc | Asym SLOPE gate* |
| 20 | 1 | 313 | 952 | 328 | 1074 | 3.4 | 0.9 | 0.5 | OR-AND-aTc | aTc gate |
| 30 | 1 | 313 | 2118 | 328 | 1867 | 6.8 | 0.9 | 0.5 | OR-(a)N(I)-aTc | aTc gate |
| 50 | 1 | 313 | 5802 | 328 | 5829 | 18.6 | 1.0 | 0.5 | OR-AND-aTc | aTc gate |

* Refers to the "asymmetric SLOPE (*a*=0.5, *l*=0.5) gate" in the OR-(a)N(I)-aTc space.

**Supplementary Table S12:** GFP expression data with the corresponding *a*, *l*, *r*, logic parameter space and logic behavior for construct pAE_LG05. Basal always corresponds to aTc=0 ng/mL and IPTG=0 mM.

| **Inducer concentrations** | | **Condition** | | | | **Logic parameters** | | | | |
|---|---|---|---|---|---|---|---|---|---|---|
| **aTc (ng/mL)** | **IPTG (mM)** | **Basal** | **aTc** | **IPTG** | **a+I** | ***r*** | ***a*** | ***l*** | **Logic Space** | **Behavior** |
| 10 | 0.01 | 317 | 342 | 324 | 353 | 1.1 | 0.5 | 0.5 | OR-AND-aTc | Asym SLOPE gate |
| 20 | 0.01 | 317 | 408 | 324 | 425 | 1.3 | 0.8 | 0.5 | OR-AND-aTc | aTc gate |
| 30 | 0.01 | 317 | 481 | 324 | 507 | 1.6 | 0.8 | 0.5 | OR-AND-aTc | aTc gate |
| 50 | 0.01 | 317 | 745 | 324 | 957 | 3.0 | 0.8 | 0.6 | OR-AND-aTc | aTc gate |
| 10 | 0.02 | 317 | 342 | 327 | 356 | 1.1 | 0.4 | 0.5 | OR-AND-aTc | Asym SLOPE gate |
| 20 | 0.02 | 317 | 408 | 327 | 437 | 1.4 | 0.7 | 0.6 | OR-AND-aTc | Asym SLOPE gate |
| 30 | 0.02 | 317 | 481 | 327 | 535 | 1.7 | 0.7 | 0.6 | OR-AND-aTc | Asym SLOPE gate |
| 50 | 0.02 | 317 | 745 | 327 | 892 | 2.8 | 0.8 | 0.6 | OR-AND-aTc | aTc gate |
| 10 | 0.5 | 317 | 342 | 337 | 380 | 1.2 | 0.1 | 0.6 | OR-AND-aTc | SLOPE gate |
| 20 | 0.5 | 317 | 408 | 337 | 493 | 1.6 | 0.4 | 0.6 | OR-AND-aTc | Asym AND gate |
| 30 | 0.5 | 317 | 481 | 337 | 651 | 2.1 | 0.5 | 0.7 | OR-AND-aTc | Asym AND gate |
| 50 | 0.5 | 317 | 745 | 337 | 1426 | 4.5 | 0.5 | 0.7 | OR-AND-aTc | Asym AND gate |
| 10 | 1 | 317 | 342 | 332 | 395 | 1.2 | 0.1 | 0.7 | OR-AND-aTc | SLOPE gate |
| 20 | 1 | 317 | 408 | 332 | 510 | 1.6 | 0.4 | 0.7 | OR-AND-aTc | Asym AND gate |
| 30 | 1 | 317 | 481 | 332 | 686 | 2.2 | 0.5 | 0.7 | OR-AND-aTc | Asym AND gate |
| 50 | 1 | 317 | 745 | 332 | 1437 | 4.5 | 0.5 | 0.7 | OR-AND-aTc | Asym AND gate |

**Supplementary Table S13:** GFP expression data with the corresponding *a*, *l*, *r*, logic parameter space and logic behavior for construct pAE_LG06. Basal always corresponds to aTc=0 ng/mL and IPTG=0 mM.

| **Inducer concentrations** | | **Condition** | | | | **Logic parameters** | | | | |
|---|---|---|---|---|---|---|---|---|---|---|
| **aTc (ng/mL)** | **IPTG (mM)** | **Basal** | **aTc** | **IPTG** | **a+I** | ***r*** | ***a*** | ***l*** | **Logic Space** | **Behavior** |
| 10 | 0.01 | 314 | 438 | 317 | 461 | 1.5 | 0.8 | 0.6 | OR-AND-aTc | aTc gate |
| 20 | 0.01 | 314 | 731 | 317 | 1082 | 3.4 | 0.7 | 0.7 | OR-AND-aTc | Asym AND gate |
| 30 | 0.01 | 314 | 1789 | 317 | 2610 | 8.3 | 0.8 | 0.6 | OR-AND-aTc | aTc gate |
| 50 | 0.01 | 314 | 3309 | 317 | 5085 | 16.2 | 0.8 | 0.6 | OR-AND-aTc | aTc gate |
| 10 | 0.02 | 314 | 438 | 330 | 482 | 1.5 | 0.7 | 0.6 | OR-AND-aTc | Asym SLOPE gate |
| 20 | 0.02 | 314 | 731 | 330 | 1431 | 4.6 | 0.5 | 0.7 | OR-AND-aTc | Asym AND gate |
| 30 | 0.02 | 314 | 1789 | 330 | 3010 | 9.6 | 0.7 | 0.6 | OR-AND-aTc | Asym SLOPE gate |
| 50 | 0.02 | 314 | 3309 | 330 | 7266 | 23.1 | 0.7 | 0.6 | OR-AND-aTc | Asym SLOPE gate |
| 10 | 0.5 | 314 | 438 | 324 | 565 | 1.8 | 0.5 | 0.7 | OR-AND-aTc | Asym AND gate |
| 20 | 0.5 | 314 | 731 | 324 | 1870 | 6.0 | 0.5 | 0.8 | OR-AND-aTc | Asym AND gate |
| 30 | 0.5 | 314 | 1789 | 324 | 4321 | 13.8 | 0.7 | 0.7 | OR-AND-aTc | Asym AND gate |
| 50 | 0.5 | 314 | 3309 | 324 | 11963 | 38.1 | 0.6 | 0.7 | OR-AND-aTc | Asym AND gate |
| 10 | 1 | 314 | 438 | 327 | 596 | 1.9 | 0.5 | 0.7 | OR-AND-aTc | Asym AND gate |
| 20 | 1 | 314 | 731 | 327 | 1725 | 5.5 | 0.5 | 0.7 | OR-AND-aTc | Asym AND gate |
| 30 | 1 | 314 | 1789 | 327 | 4788 | 15.2 | 0.6 | 0.7 | OR-AND-aTc | Asym AND gate |
| 50 | 1 | 314 | 3309 | 327 | 10201 | 32.5 | 0.7 | 0.7 | OR-AND-aTc | Asym AND gate |

**Supplementary Table S14:** GFP expression data with the corresponding *a*, *l*, *r*, logic parameter space and logic behavior for construct pAE_LG07. Basal always corresponds to aTc=0 ng/mL and IPTG=0 mM.

| **Inducer concentrations** | | **Condition** | | | | **Logic parameters** | | | | |
|---|---|---|---|---|---|---|---|---|---|---|
| **aTc (ng/mL)** | **IPTG (mM)** | **Basal** | **aTc** | **IPTG** | **a+I** | ***r*** | ***a*** | ***l*** | **Logic Space** | **Behavior** |
| 10 | 0.01 | 313 | 382 | 318 | 380 | 1.2 | 0.9 | 0.5 | OR-(a)N(I)-aTc | aTc gate |
| 20 | 0.01 | 313 | 615 | 318 | 535 | 2.0 | 0.8 | 0.6 | OR-(a)N(I)-aTc | aTc gate |
| 30 | 0.01 | 313 | 1551 | 318 | 1626 | 5.2 | 1.0 | 0.5 | OR-AND-aTc | aTc gate |
| 50 | 0.01 | 313 | 3421 | 318 | 3968 | 12.7 | 0.9 | 0.5 | OR-AND-aTc | aTc gate |
| 10 | 0.02 | 313 | 382 | 322 | 392 | 1.3 | 0.8 | 0.5 | OR-AND-aTc | aTc gate |
| 20 | 0.02 | 313 | 615 | 322 | 646 | 2.1 | 0.9 | 0.5 | OR-AND-aTc | aTc gate |
| 30 | 0.02 | 313 | 1551 | 322 | 1692 | 5.4 | 0.9 | 0.5 | OR-AND-aTc | aTc gate |
| 50 | 0.02 | 313 | 3421 | 322 | 4177 | 13.4 | 0.9 | 0.5 | OR-AND-aTc | aTc gate |
| 10 | 0.5 | 313 | 382 | 338 | 407 | 1.3 | 0.5 | 0.5 | OR-AND-aTc | Asym SLOPE gate |
| 20 | 0.5 | 313 | 615 | 338 | 650 | 2.1 | 0.8 | 0.5 | OR-AND-aTc | aTc gate |
| 30 | 0.5 | 313 | 1551 | 338 | 1702 | 5.4 | 0.9 | 0.5 | OR-AND-aTc | aTc gate |
| 50 | 0.5 | 313 | 3421 | 338 | 4091 | 13.1 | 0.9 | 0.5 | OR-AND-aTc | aTc gate |
| 10 | 1 | 313 | 382 | 341 | 374 | 1.2 | 0.5 | 0.3 | OR-(a)N(I)-aTc | Asym aTc gate |
| 20 | 1 | 313 | 615 | 341 | 593 | 2.0 | 0.8 | 0.5 | OR-(a)N(I)-aTc | aTc gate |
| 30 | 1 | 313 | 1551 | 341 | 1663 | 5.3 | 0.9 | 0.5 | OR-AND-aTc | aTc gate |
| 50 | 1 | 313 | 3421 | 341 | 4966 | 15.9 | 0.8 | 0.6 | OR-AND-aTc | aTc gate |

**Supplementary Table S15:** GFP expression data with the corresponding *a*, *l*, *r*, logic parameter space and logic behavior for construct pAE_LG13. Basal always corresponds to aTc=0 ng/mL and IPTG=0 mM.

| **Inducer concentrations** | | **Condition** | | | | **Logic parameters** | | | | |
|---|---|---|---|---|---|---|---|---|---|---|
| **aTc (ng/mL)** | **IPTG (mM)** | **Basal** | **aTc** | **IPTG** | **a+I** | ***r*** | ***a*** | ***l*** | **Logic Space** | **Behavior** |
| 10 | 0.01 | 329 | 540 | 312 | 816 | 2.6 | 0.5 | 0.7 | (a)I(I)-AND-aTc | Asym AND gate |
| 20 | 0.01 | 329 | 2005 | 312 | 2602 | 8.3 | 0.9 | 0.5 | (a)I(I)-AND-aTc | aTc gate |
| 30 | 0.01 | 329 | 3535 | 312 | 4010 | 12.9 | 0.9 | 0.5 | (a)I(I)-AND-aTc | aTc gate |
| 50 | 0.01 | 329 | 5872 | 312 | 7274 | 23.3 | 0.9 | 0.5 | (a)I(I)-AND-aTc | aTc gate |
| 10 | 0.02 | 329 | 540 | 322 | 837 | 2.6 | 0.5 | 0.7 | (a)I(I)-AND-aTc | Asym AND gate |
| 20 | 0.02 | 329 | 2005 | 322 | 2562 | 8.0 | 0.9 | 0.6 | (a)I(I)-AND-aTc | aTc gate |
| 30 | 0.02 | 329 | 3535 | 322 | 4259 | 13.2 | 0.9 | 0.5 | (a)I(I)-AND-aTc | aTc gate |
| 50 | 0.02 | 329 | 5872 | 322 | 8466 | 26.3 | 0.9 | 0.6 | (a)I(I)-AND-aTc | aTc gate |
| 10 | 0.5 | 329 | 540 | 331 | 655 | 2.0 | 0.7 | 0.6 | OR-AND-aTc | Asym AND gate |
| 20 | 0.5 | 329 | 2005 | 331 | 3379 | 10.3 | 0.8 | 0.6 | OR-AND-aTc | aTc gate |
| 30 | 0.5 | 329 | 3535 | 331 | 4961 | 15.1 | 0.9 | 0.6 | OR-AND-aTc | aTc gate |
| 50 | 0.5 | 329 | 5872 | 331 | 9889 | 30.1 | 0.8 | 0.6 | OR-AND-aTc | aTc gate |
| 10 | 1 | 329 | 540 | 352 | 423 | 1.6 | 0.4 | 0.7 | OR-(a)N(I)-aTc | Asym (a)N(I) gate |
| 20 | 1 | 329 | 2005 | 352 | 3389 | 10.3 | 0.7 | 0.6 | OR-AND-aTc | Asym SLOPE gate |
| 30 | 1 | 329 | 3535 | 352 | 6025 | 18.3 | 0.8 | 0.6 | OR-AND-aTc | aTc gate |
| 50 | 1 | 329 | 5872 | 352 | 10706 | 32.5 | 0.8 | 0.6 | OR-AND-aTc | aTc gate |

**Supplementary Table S16:** GFP expression data with the corresponding *a*, *l*, *r*, logic parameter space and logic behavior for construct pAE_LG15. Basal always corresponds to aTc=0 ng/mL and IPTG=0 mM.

| **Inducer concentrations** | | **Condition** | | | | **Logic parameters** | | | | |
|---|---|---|---|---|---|---|---|---|---|---|
| **aTc (ng/mL)** | **IPTG (mM)** | **Basal** | **aTc** | **IPTG** | **a+I** | ***r*** | ***a*** | ***l*** | **Logic Space** | **Behavior** |
| 10 | 0.01 | 307 | 334 | 4086 | 4032 | 13.3 | 1.0 | 0.5 | OR-(I)N(a)-IPTG | IPTG gate |
| 20 | 0.01 | 307 | 454 | 4086 | 4092 | 13.3 | 0.8 | 0.4 | OR-AND-IPTG | IPTG gate |
| 30 | 0.01 | 307 | 613 | 4086 | 3957 | 13.3 | 0.7 | 0.4 | OR-(I)N(a)-IPTG | Asym IPTG gate |
| 50 | 0.01 | 307 | 883 | 4086 | 6895 | 22.4 | 0.5 | 0.4 | OR-AND-IPTG | Asym SLOPE gate |
| 10 | 0.02 | 307 | 334 | 5994 | 6112 | 19.9 | 1.0 | 0.5 | OR-AND-IPTG | IPTG gate |
| 20 | 0.02 | 307 | 454 | 5994 | 6294 | 20.5 | 0.9 | 0.4 | OR-AND-IPTG | IPTG gate |
| 30 | 0.02 | 307 | 613 | 5994 | 6637 | 21.6 | 0.7 | 0.4 | OR-AND-IPTG | Asym SLOPE gate |
| 50 | 0.02 | 307 | 883 | 5994 | 10821 | 35.2 | 0.5 | 0.4 | OR-AND-IPTG | Asym SLOPE gate |
| 10 | 0.5 | 307 | 334 | 11816 | 11947 | 38.9 | 1.0 | 0.5 | OR-AND-IPTG | IPTG gate |
| 20 | 0.5 | 307 | 454 | 11816 | 12276 | 40.0 | 0.9 | 0.5 | OR-AND-IPTG | IPTG gate |
| 30 | 0.5 | 307 | 613 | 11816 | 14410 | 46.9 | 0.8 | 0.4 | OR-AND-IPTG | IPTG gate |
| 50 | 0.5 | 307 | 883 | 11816 | 22437 | 73.0 | 0.6 | 0.5 | OR-AND-IPTG | Asym SLOPE gate |
| 10 | 1 | 307 | 334 | 12415 | 12895 | 42.0 | 1.0 | 0.5 | OR-AND-IPTG | IPTG gate |
| 20 | 1 | 307 | 454 | 12415 | 12970 | 42.2 | 0.9 | 0.5 | OR-AND-IPTG | IPTG gate |
| 30 | 1 | 307 | 613 | 12415 | 15991 | 52.0 | 0.8 | 0.4 | OR-AND-IPTG | IPTG gate |
| 50 | 1 | 307 | 883 | 12415 | 24218 | 78.8 | 0.6 | 0.5 | OR-AND-IPTG | Asym SLOPE gate |

**Supplementary Table S17:** GFP expression data with the corresponding *a*, *l*, *r*, logic parameter space and logic behavior for construct pAE_LG21. Basal always corresponds to aTc=0 ng/mL and IPTG=0 mM.

| **Inducer concentrations** | | **Condition** | | | | **Logic parameters** | | | | |
|---|---|---|---|---|---|---|---|---|---|---|
| **aTc (ng/mL)** | **IPTG (mM)** | **Basal** | **aTc** | **IPTG** | **a+I** | ***r*** | ***a*** | ***l*** | **Logic Space** | **Behavior** |
| 10 | 0.01 | 306 | 335 | 2561 | 2211 | 8.4 | 0.9 | 0.5 | OR-(I)N(a)-IPTG | IPTG gate |
| 20 | 0.01 | 306 | 416 | 2561 | 2853 | 9.3 | 0.8 | 0.5 | OR-AND-IPTG | IPTG gate |
| 30 | 0.01 | 306 | 680 | 2561 | 3624 | 11.8 | 0.5 | 0.4 | OR-AND-IPTG | Asym SLOPE gate |
| 50 | 0.01 | 306 | 1262 | 2561 | 5336 | 17.4 | 0.2 | 0.4 | OR-AND-IPTG | SLOPE gate |
| 10 | 0.02 | 306 | 335 | 4011 | 3604 | 13.1 | 0.9 | 0.5 | OR-(I)N(a)-IPTG | IPTG gate |
| 20 | 0.02 | 306 | 416 | 4011 | 4548 | 14.8 | 0.8 | 0.5 | OR-AND-IPTG | IPTG gate |
| 30 | 0.02 | 306 | 680 | 4011 | 5569 | 18.2 | 0.6 | 0.4 | OR-AND-IPTG | Asym SLOPE gate |
| 50 | 0.02 | 306 | 1262 | 4011 | 11375 | 37.1 | 0.3 | 0.4 | OR-AND-IPTG | Asym SLOPE gate |
| 10 | 0.5 | 306 | 335 | 8007 | 8987 | 29.3 | 0.9 | 0.5 | OR-AND-IPTG | IPTG gate |
| 20 | 0.5 | 306 | 416 | 8007 | 9070 | 29.6 | 0.9 | 0.5 | OR-AND-IPTG | IPTG gate |
| 30 | 0.5 | 306 | 680 | 8007 | 12126 | 39.6 | 0.7 | 0.4 | OR-AND-IPTG | Asym SLOPE gate |
| 50 | 0.5 | 306 | 1262 | 8007 | 22972 | 75.0 | 0.4 | 0.5 | OR-AND-IPTG | Asym SLOPE gate |
| 10 | 1 | 306 | 335 | 9651 | 8689 | 31.5 | 0.9 | 0.5 | OR-(I)N(a)-IPTG | IPTG gate |
| 20 | 1 | 306 | 416 | 9651 | 9856 | 32.2 | 0.9 | 0.5 | OR-AND-IPTG | IPTG gate |
| 30 | 1 | 306 | 680 | 9651 | 12274 | 40.1 | 0.7 | 0.4 | OR-AND-IPTG | Asym SLOPE gate |
| 50 | 1 | 306 | 1262 | 9651 | 26493 | 86.5 | 0.5 | 0.5 | OR-AND-IPTG | Asym SLOPE gate |

**Supplementary Table S18:** GFP expression data with the corresponding *a*, *l*, *r*, logic parameter space and logic behavior for construct pAE_LG23. Basal always corresponds to aTc=0 ng/mL and IPTG=0 mM.

| **Inducer concentrations** | | **Condition** | | | | **Logic parameters** | | | | |
|---|---|---|---|---|---|---|---|---|---|---|
| **aTc (ng/mL)** | **IPTG (mM)** | **Basal** | **aTc** | **IPTG** | **a+I** | ***r*** | ***a*** | ***l*** | **Logic Space** | **Behavior** |
| 10 | 0.01 | 335 | 429 | 9786 | 8620 | 29.2 | 0.9 | 0.5 | OR-(I)N(a)-IPTG | IPTG gate |
| 20 | 0.01 | 335 | 739 | 9786 | 8980 | 29.2 | 0.7 | 0.4 | OR-(I)N(a)-IPTG | Asym SLOPE gate* |
| 30 | 0.01 | 335 | 1440 | 9786 | 9995 | 29.8 | 0.6 | 0.3 | OR-AND-IPTG | Asym IPTG gate |
| 50 | 0.01 | 335 | 3233 | 9786 | 9284 | 29.2 | 0.3 | 0.2 | OR-(I)N(a)-IPTG | Asym IPTG gate |
| 10 | 0.02 | 335 | 429 | 12128 | 10343 | 36.2 | 0.9 | 0.5 | OR-(I)N(a)-IPTG | IPTG gate |
| 20 | 0.02 | 335 | 739 | 12128 | 13214 | 39.4 | 0.8 | 0.4 | OR-AND-IPTG | IPTG gate |
| 30 | 0.02 | 335 | 1440 | 12128 | 12045 | 36.2 | 0.6 | 0.3 | OR-(I)N(a)-IPTG | Asym IPTG gate |
| 50 | 0.02 | 335 | 3233 | 12128 | 14092 | 42.1 | 0.4 | 0.2 | OR-AND-IPTG | Asym IPTG gate |
| 10 | 0.5 | 335 | 429 | 15626 | 15733 | 47.0 | 0.9 | 0.5 | OR-AND-IPTG | IPTG gate |
| 20 | 0.5 | 335 | 739 | 15626 | 15998 | 47.7 | 0.8 | 0.4 | OR-AND-IPTG | IPTG gate |
| 30 | 0.5 | 335 | 1440 | 15626 | 23256 | 69.4 | 0.6 | 0.4 | OR-AND-IPTG | Asym IPTG gate |
| 50 | 0.5 | 335 | 3233 | 15626 | 49741 | 148.5 | 0.3 | 0.4 | OR-AND-IPTG | Asym SLOPE gate |
| 10 | 1 | 335 | 429 | 15679 | 15580 | 46.8 | 0.9 | 0.5 | OR-(I)N(a)-IPTG | IPTG gate |
| 20 | 1 | 335 | 739 | 15679 | 16805 | 50.2 | 0.8 | 0.4 | OR-AND-IPTG | IPTG gate |
| 30 | 1 | 335 | 1440 | 15679 | 22992 | 68.6 | 0.6 | 0.4 | OR-AND-IPTG | Asym IPTG gate |
| 50 | 1 | 335 | 3233 | 15679 | 33779 | 100.8 | 0.3 | 0.3 | OR-AND-IPTG | Asym IPTG gate |

* Refers to the "asymmetric SLOPE (*a*=0.5, *l*=0.5) gate" in the OR-(a)N(I)-aTc space.

**Supplementary Table S19:** GFP expression data with the corresponding *a*, *l*, *r*, logic parameter space and logic behavior for construct pAE_LG25. Basal always corresponds to aTc=0 ng/mL and IPTG=0 mM.

| **Inducer concentrations** | | **Condition** | | | | **Logic parameters** | | | | |
|---|---|---|---|---|---|---|---|---|---|---|
| **aTc (ng/mL)** | **IPTG (mM)** | **Basal** | **aTc** | **IPTG** | **a+I** | ***r*** | ***a*** | ***l*** | **Logic Space** | **Behavior** |
| 10 | 0.01 | 310 | 350 | 10053 | 10131 | 32.7 | 1.0 | 0.5 | OR-AND-IPTG | IPTG gate |
| 20 | 0.01 | 310 | 733 | 10053 | 9896 | 32.5 | 0.7 | 0.4 | OR-(I)N(a)-IPTG | Asym SLOPE gate* |
| 30 | 0.01 | 310 | 1490 | 10053 | 10883 | 35.1 | 0.5 | 0.3 | OR-AND-IPTG | Asym IPTG gate |
| 50 | 0.01 | 310 | 3099 | 10053 | 11853 | 38.3 | 0.3 | 0.2 | OR-AND-IPTG | Asym IPTG gate |
| 10 | 0.02 | 310 | 350 | 12744 | 12666 | 41.1 | 1.0 | 0.5 | OR-(I)N(a)-IPTG | IPTG gate |
| 20 | 0.02 | 310 | 733 | 12744 | 12988 | 41.9 | 0.8 | 0.4 | OR-AND-IPTG | IPTG gate |
| 30 | 0.02 | 310 | 1490 | 12744 | 13915 | 44.9 | 0.6 | 0.3 | OR-AND-IPTG | Asym IPTG gate |
| 50 | 0.02 | 310 | 3099 | 12744 | 19675 | 63.5 | 0.3 | 0.3 | OR-AND-IPTG | Asym IPTG gate |
| 10 | 0.5 | 310 | 350 | 15299 | 15497 | 50.0 | 1.0 | 0.5 | OR-AND-IPTG | IPTG gate |
| 20 | 0.5 | 310 | 733 | 15299 | 14585 | 49.4 | 0.8 | 0.4 | OR-(I)N(a)-IPTG | IPTG gate |
| 30 | 0.5 | 310 | 1490 | 15299 | 16158 | 52.2 | 0.6 | 0.3 | OR-AND-IPTG | Asym IPTG gate |
| 50 | 0.5 | 310 | 3099 | 15299 | 37193 | 120.1 | 0.3 | 0.4 | OR-AND-IPTG | Asym IPTG gate |
| 10 | 1 | 310 | 350 | 14976 | 15175 | 49.0 | 1.0 | 0.5 | OR-AND-IPTG | IPTG gate |
| 20 | 1 | 310 | 733 | 14976 | 15130 | 48.9 | 0.8 | 0.4 | OR-AND-IPTG | IPTG gate |
| 30 | 1 | 310 | 1490 | 14976 | 12350 | 48.4 | 0.5 | 0.3 | OR-(I)N(a)-IPTG | Asym IPTG gate |
| 50 | 1 | 310 | 3099 | 14976 | 39472 | 127.4 | 0.3 | 0.4 | OR-AND-IPTG | Asym IPTG gate |

* Refers to the “asymmetric SLOPE (*a*=0.5, *l*=0.5) gate” in the OR-(a)N(I)-aTc space.

**Supplementary Table S20:** GFP expression data with the corresponding *a*, *l*, *r*, logic parameter space and logic behavior for construct pAE_LG26. Basal always corresponds to aTc=0 ng/mL and IPTG=0 mM.

| **Inducer concentrations** | | **Condition** | | | | **Logic parameters** | | | | |
|---|---|---|---|---|---|---|---|---|---|---|
| **aTc (ng/mL)** | **IPTG (mM)** | **Basal** | **aTc** | **IPTG** | **a+I** | ***r*** | ***a*** | ***l*** | **Logic Space** | **Behavior** |
| 10 | 0.01 | 442 | 589 | 6579 | 7092 | 16.0 | 0.9 | 0.5 | OR-AND-IPTG | IPTG gate |
| 20 | 0.01 | 442 | 1032 | 6579 | 8314 | 18.8 | 0.6 | 0.4 | OR-AND-IPTG | Asym SLOPE gate |
| 30 | 0.01 | 442 | 2317 | 6579 | 9786 | 22.1 | 0.3 | 0.3 | OR-AND-IPTG | Asym IPTG gate |
| 50 | 0.01 | 442 | 4199 | 6579 | 11760 | 26.6 | 0.1 | 0.2 | OR-AND-IPTG | OR gate |
| 10 | 0.02 | 442 | 589 | 9535 | 10211 | 23.1 | 0.9 | 0.5 | OR-AND-IPTG | IPTG gate |
| 20 | 0.02 | 442 | 1032 | 9535 | 11136 | 25.2 | 0.7 | 0.4 | OR-AND-IPTG | Asym SLOPE gate |
| 30 | 0.02 | 442 | 2317 | 9535 | 13293 | 30.1 | 0.4 | 0.3 | OR-AND-IPTG | Asym IPTG gate |
| 50 | 0.02 | 442 | 4199 | 9535 | 17241 | 39.0 | 0.2 | 0.3 | OR-AND-IPTG | Asym IPTG gate |
| 10 | 0.5 | 442 | 589 | 13124 | 11903 | 29.7 | 0.9 | 0.5 | OR-(I)N(a)-IPTG | IPTG gate |
| 20 | 0.5 | 442 | 1032 | 13124 | 14534 | 32.9 | 0.7 | 0.4 | OR-AND-IPTG | Asym SLOPE gate |
| 30 | 0.5 | 442 | 2317 | 13124 | 16379 | 37.0 | 0.5 | 0.3 | OR-AND-IPTG | Asym IPTG gate |
| 50 | 0.5 | 442 | 4199 | 13124 | 20640 | 46.7 | 0.3 | 0.3 | OR-AND-IPTG | Asym IPTG gate |
| 10 | 1 | 442 | 589 | 11329 | 12450 | 28.2 | 0.9 | 0.5 | OR-AND-IPTG | IPTG gate |
| 20 | 1 | 442 | 1032 | 11329 | 13886 | 31.4 | 0.7 | 0.4 | OR-AND-IPTG | Asym SLOPE gate |
| 30 | 1 | 442 | 2317 | 11329 | 16564 | 37.5 | 0.4 | 0.3 | OR-AND-IPTG | Asym IPTG gate |
| 50 | 1 | 442 | 4199 | 11329 | 23767 | 53.7 | 0.2 | 0.3 | OR-AND-IPTG | Asym IPTG gate |

**Supplementary Table S21:** GFP expression data with the corresponding *a*, *l*, *r*, logic parameter space and logic behavior for construct pAE_LG27. Basal always corresponds to aTc=0 ng/mL and IPTG=0 mM.

| **Inducer concentrations** | | **Condition** | | | | **Logic parameters** | | | | |
|---|---|---|---|---|---|---|---|---|---|---|
| **aTc (ng/mL)** | **IPTG (mM)** | **Basal** | **aTc** | **IPTG** | **a+I** | ***r*** | ***a*** | ***l*** | **Logic Space** | **Behavior** |
| 10 | 0.01 | 12343 | 12107 | 14293 | 14182 | 1.2 | 0.8 | 0.5 | (a)I(I)-(I)N(a)-IPTG | IPTG gate |
| 20 | 0.01 | 12343 | 12649 | 14293 | 14205 | 1.2 | 0.8 | 0.4 | OR-(I)N(a)-IPTG | IPTG gate |
| 30 | 0.01 | 12343 | 13949 | 14293 | 16594 | 1.3 | 0.1 | 0.5 | OR-AND-IPTG | SLOPE gate |
| 50 | 0.01 | 12343 | 16624 | 14293 | 19187 | 1.6 | 0.3 | 0.5 | OR-AND-aTc | Asym SLOPE gate |
| 10 | 0.02 | 12343 | 12107 | 14723 | 15221 | 1.3 | 0.8 | 0.5 | (a)I(I)-AND-IPTG | IPTG gate |
| 20 | 0.02 | 12343 | 12649 | 14723 | 15413 | 1.2 | 0.7 | 0.5 | OR-AND-IPTG | Asym SLOPE gate |
| 30 | 0.02 | 12343 | 13949 | 14723 | 16585 | 1.3 | 0.2 | 0.5 | OR-AND-IPTG | SLOPE gate |
| 50 | 0.02 | 12343 | 16624 | 14723 | 19623 | 1.6 | 0.3 | 0.5 | OR-AND-aTc | Asym SLOPE gate |
| 10 | 0.5 | 12343 | 12107 | 15369 | 15571 | 1.3 | 0.9 | 0.5 | (a)I(I)-AND-IPTG | IPTG gate |
| 20 | 0.5 | 12343 | 12649 | 15369 | 16212 | 1.3 | 0.7 | 0.6 | OR-AND-IPTG | Asym SLOPE gate |
| 30 | 0.5 | 12343 | 13949 | 15369 | 17490 | 1.4 | 0.3 | 0.5 | OR-AND-IPTG | Asym SLOPE gate |
| 50 | 0.5 | 12343 | 16624 | 15369 | 20170 | 1.6 | 0.2 | 0.5 | OR-AND-aTc | SLOPE gate |
| 10 | 1 | 12343 | 12107 | 15632 | 15438 | 1.3 | 0.9 | 0.5 | (a)I(I)-(I)N(a)-IPTG | IPTG gate |
| 20 | 1 | 12343 | 12649 | 15632 | 15804 | 1.3 | 0.9 | 0.5 | OR-AND-IPTG | IPTG gate |
| 30 | 1 | 12343 | 13949 | 15632 | 17289 | 1.4 | 0.3 | 0.5 | OR-AND-IPTG | Asym SLOPE gate |
| 50 | 1 | 12343 | 16624 | 15632 | 20313 | 1.6 | 0.1 | 0.5 | OR-AND-aTc | SLOPE gate |

### Mathematical Modeling Parameters and AIC

**Supplementary Table S22:** Nomenclature table for AND and OR gate mathematical models. Literature-derived parameters held fixed for transfer function modelling and sources are denoted. Values are not given for fitted constants here but are given in Supplementary Figures S11-S18, S25-S30.

| Parameter | Value | Units | Source | Description |
|---|---|---|---|---|
| [RNAP](Munro *et al*, 2016) | 3000 | nM | Literature(5) | Concentration of RNAP |
| [LacI](Munro *et al*, 2016) | 100 | nM | Literature(5) | Concentration of LacI |
| [TetR](Munro *et al*, 2016) | 100 | nM | Literature(5) | Concentration of TetR |
| $K_{a,IPTG:LacI}$(Chens *et al*, 1994) | 6.7e-04 | $nM^{-1}$ | Literature(6) | Association constant of IPTG to LacI |
| $K_{a,TetR:TetO}$(Kamionka *et al*, 2004) | 5.6 | $nM^{-1}$ | Literature(7) | Association constant of TetR to TetO |
| $K_{a,aTc:TetR}$ | - | $nM^{-1}$ | Fitted | Association constant of aTc to TetR |
| $K_{a,LacI:LacO}$ | - | $nM^{-1}$ | Fitted | Association constant of LacI to LacO |
| $K_{a,RNAP:pLac}$ | - | $nM^{-1}$ | Fitted | Association constant of RNAP to pLac |
| $K_{a,RNAP:pTet}$ | - | $nM^{-1}$ | Fitted | Association constant of RNAP to pTet |
| $a_{pTet}$ | - | n/a | Fitted | Promoter weight for pLac in OR gate modeling |
| $a_{pTet}$ | - | n/a | Fitted | Promoter weight for pTet in OR gate modeling |
| b | - | n/a | Fitted | Cooperativity weight |
| c | - | n/a | Fitted | Cooperativity threshold |
| k | - | n/a | Fitted | Translation constant |
| a | - | n/a | Fitted | AND gate readthrough constant |

**Supplementary Table S23:** Fitted promoter weights for OR gate model equations 2 and 3. The reported values for each construct correspond to the values from the construct's best fitting model equation, determined by the AIC and reported in Supplementary Table S7.

| **Construct** | $\mathbf{a_{pTet}}$ | $\mathbf{a_{pLac}}$ |
|---|---|---|
| pAE_LG15 | 0.04 | 0.55 |
| pAE_LG21 | 0.03 | 0.51 |
| pAE_LG23 | 0.12 | 0.5 |
| pAE_LG25 | 0.08 | 0.42 |
| pAE_LG26 | 1.83 | 0.41 |
| pAE_LG27 | 0.1 | 0.38 |

**Supplementary Table S24:** AIC values quantifying the goodness of fit for each model equation fit to a 5x5 grid of different aTc and IPTG concentrations for each AND construct. 'Hill' in the model equation name indicates a fitting of the transfer function Hill coefficients; model equations without 'Hill' have Hill equations set to 2.

| | Construct name pAE_ | | | | | | | |
|---|---|---|---|---|---|---|---|---|
| **Model equation** | **LG03** | **LG02** | **LG06** | **LG05** | **LG13** | **LG07** | **LG04** | **LG01** |
| **Eqn S4** | -54.37 | -79.19 | -69.44 | -94.86 | -73.33 | -125.13 | -86.82 | -102.52 |
| **Eqn S4- Hill** | -77.31 | -75.52 | -95.61 | -142.71 | -84.64 | -125.28 | -110.84 | -96.70 |
| **Eqn S5** | -46.41 | -65.06 | -69.44 | -116.46 | -71.43 | -124.02 | -86.23 | -101.40 |
| **Eqn S5 Hill** | -77.31 | -76.08 | -95.61 | -139.98 | -82.89 | -123.57 | -92.74 | -96.70 |
| **Eqn S6- Hill** | -74.10 | -89.28 | -126.98 | -180.27 | -96.53 | -129.91 | -99.59 | -104.90 |
| **Eqn S7- Hill** | -80.31 | -110.29 | -119.07 | -175.87 | -91.90 | -126.90 | -99.64 | -97.97 |

**Supplementary Table S25:** AIC values quantifying the goodness of fit for each model equation fit to a 5x5 grid of different aTc and IPTG concentrations for each OR construct. 'Hill' in the model equation name indicates a fitting of the transfer function Hill coefficients; model equations without 'Hill' have Hill parameters *m* and *n* set to 2.

| | Construct name pAE_ | | | | | |
|---|---|---|---|---|---|---|
| **Model equation** | **LG26** | **LG15** | **LG23** | **LG21** | **LG27** | **LG25** |
| **Eqn S8** | -32.72 | 67.19 | 44.19 | 67.57 | 66.28 | 43.95 |
| **Eqn S8- Hill** | -10.13 | 13.94 | -7.01 | 12.09 | -43.92 | 2.98 |
| **Eqn S9** | -84.30 | -67.00 | -49.36 | -62.65 | -111.02 | -55.97 |
| **Eqn S9- Hill** | -75.20 | -64.20 | - | -60.32 | -112.75 | -50.58 |
| **Eqn S10- Hill** | -67.67 | -68.13 | - | -69.33 | -110.99 | -51.70 |

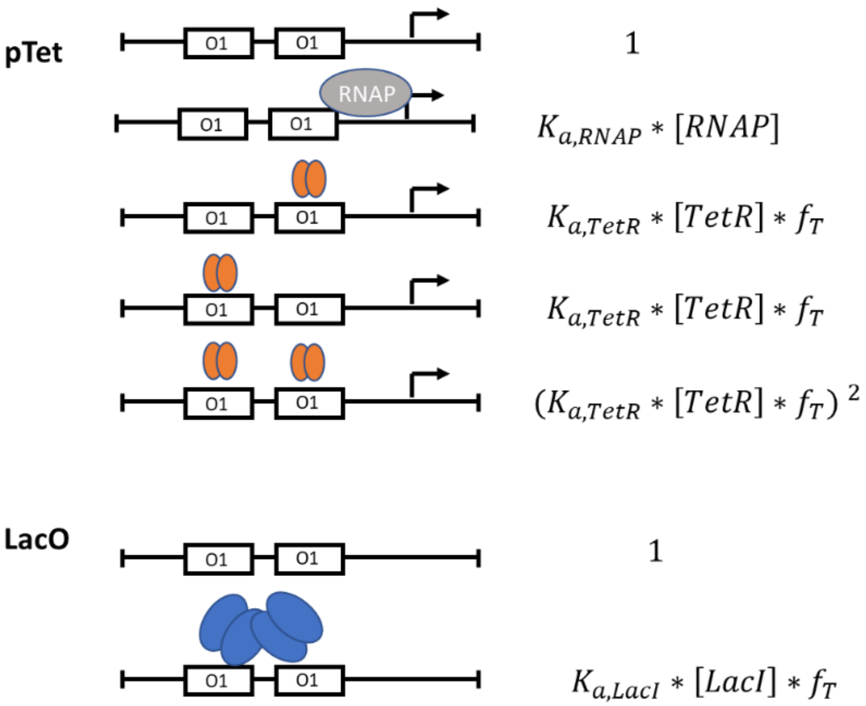

**Supplementary Figure S1:**

Graphical representation of pTet and LacO occupancy partitioning, adapted from Tamsir *et al* (Tamsir *et al*, 2011). The mathematical expressions representing each possible promoter state, shown on the right, are assembled into transfer functions, as shown in Equation S3.

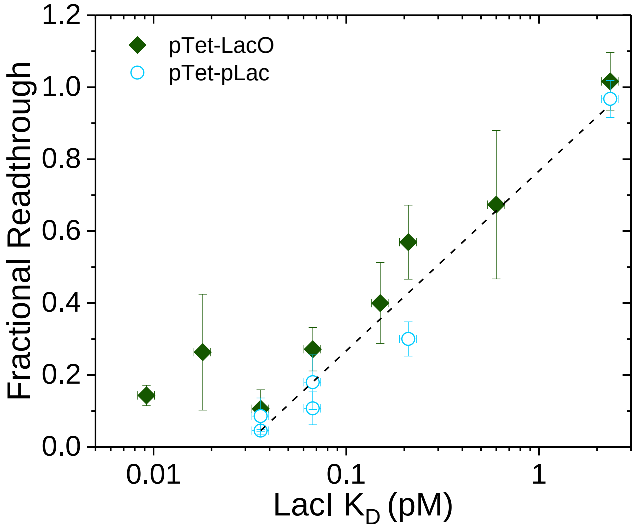

**Supplementary Figure S2:**

The fractional readthrough for constructs designed as OR gates (blue circles) is similar to that of constructs designed as AND gates (green rhombi). In fact, when fitted together for LacI $K_D$ values greater than ~0.03 pM, a good correlation is obtained (Adj. $R^2$=0.97). This suggests that the main mode of TI in our set of tested tandem promoters is also roadblock, as it was in the case when a LacO was placed downstream of pTet.

**AND Gate Logic Behaviors**

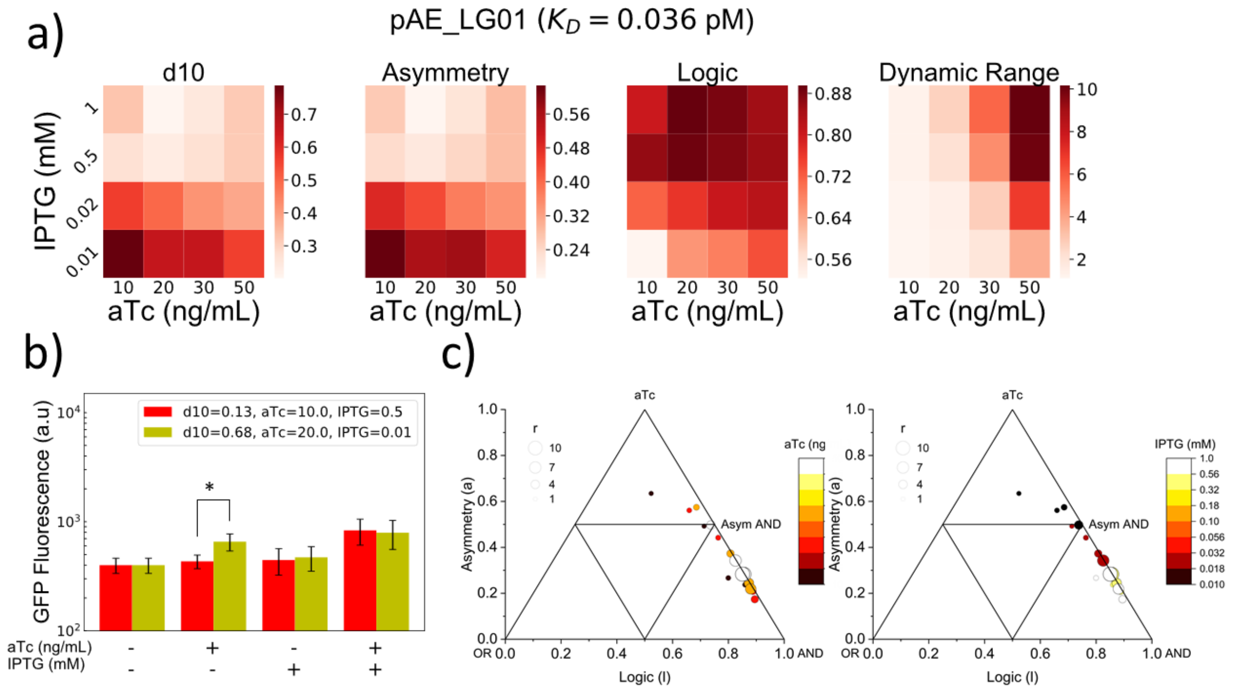

**Supplementary Figure S3:**

a) Heat maps of logic parameters of pAE_LG01. b) Bar plot of minimum and maximum d10 conditions for pAE_LG01. c) Triangle plots of pAE_LG01 at varying aTc and IPTG conditions, showing trends in aTc (left) and IPTG (right).

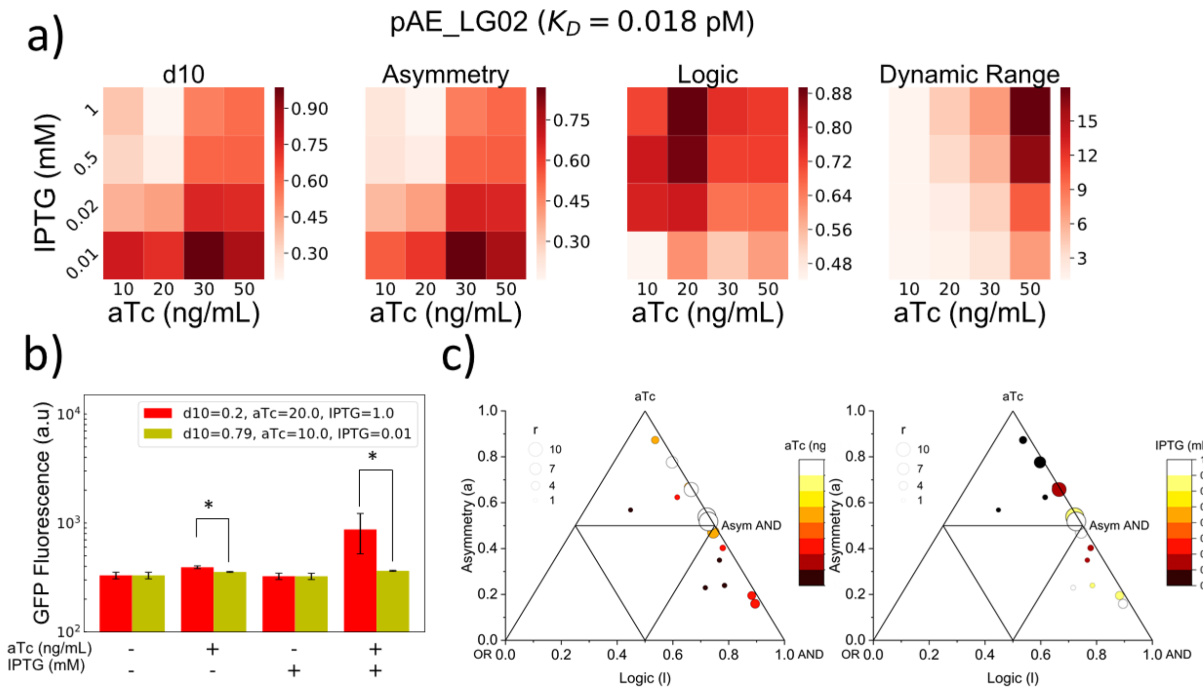

**Supplementary Figure S4:**

a) Heat maps of logic parameters of pAE_LG02. b) Bar plot of minimum and maximum d10 conditions for pAE_LG02. c) Triangle plots of pAE_LG02 at varying aTc and IPTG conditions, showing trends in aTc (left) and IPTG (right).

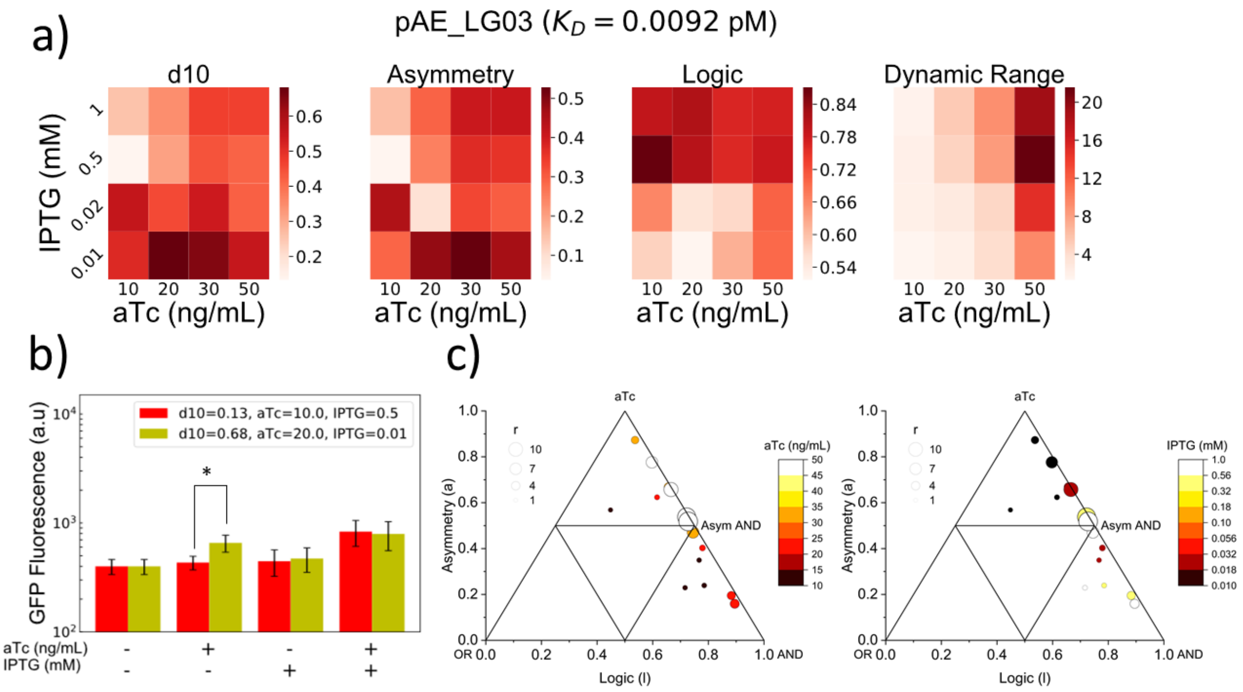

**Supplementary Figure S5:**

a) Heat maps of logic parameters of pAE_LG03. b) Bar plot of minimum and maximum d10 conditions for pAE_LG03. c) Triangle plots of pAE_LG03 at varying aTc and IPTG conditions, showing trends in aTc (left) and IPTG (right).

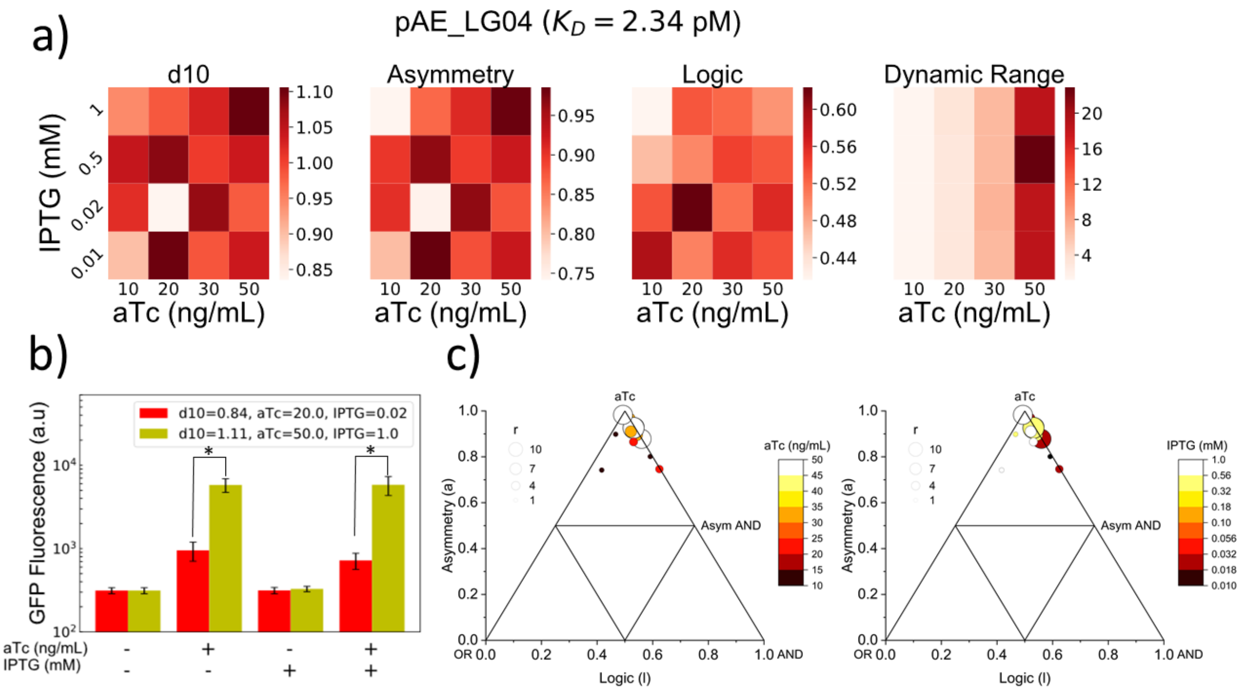

**Supplementary Figure S6:**

a) Heat maps of logic parameters of pAE_LG04. b) Bar plot of minimum and maximum d10 conditions for pAE_LG04. c) Triangle plots of pAE_LG04 at varying aTc and IPTG conditions, showing trends in aTc (left) and IPTG (right).

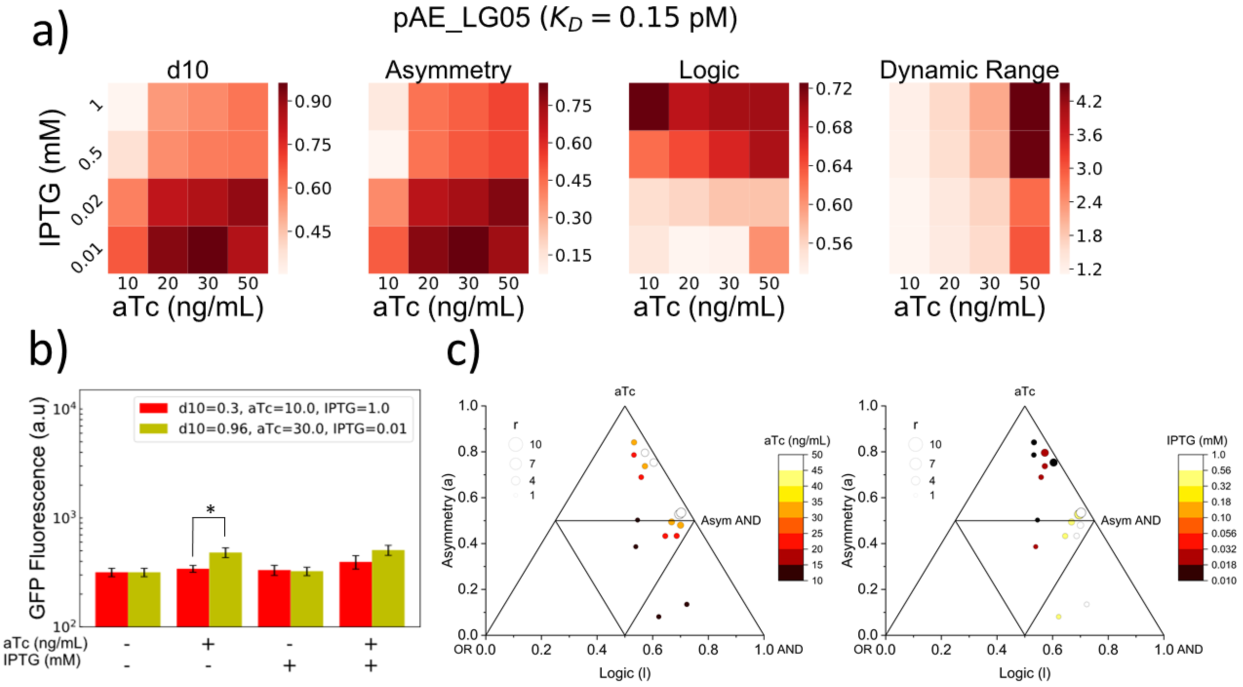

**Supplementary Figure S7:**

a) Heat maps of logic parameters of pAE_LG05. b) Bar plot of minimum and maximum d10 conditions for pAE_LG05. c) Triangle plots of pAE_LG05 at varying aTc and IPTG conditions, showing trends in aTc (left) and IPTG (right).

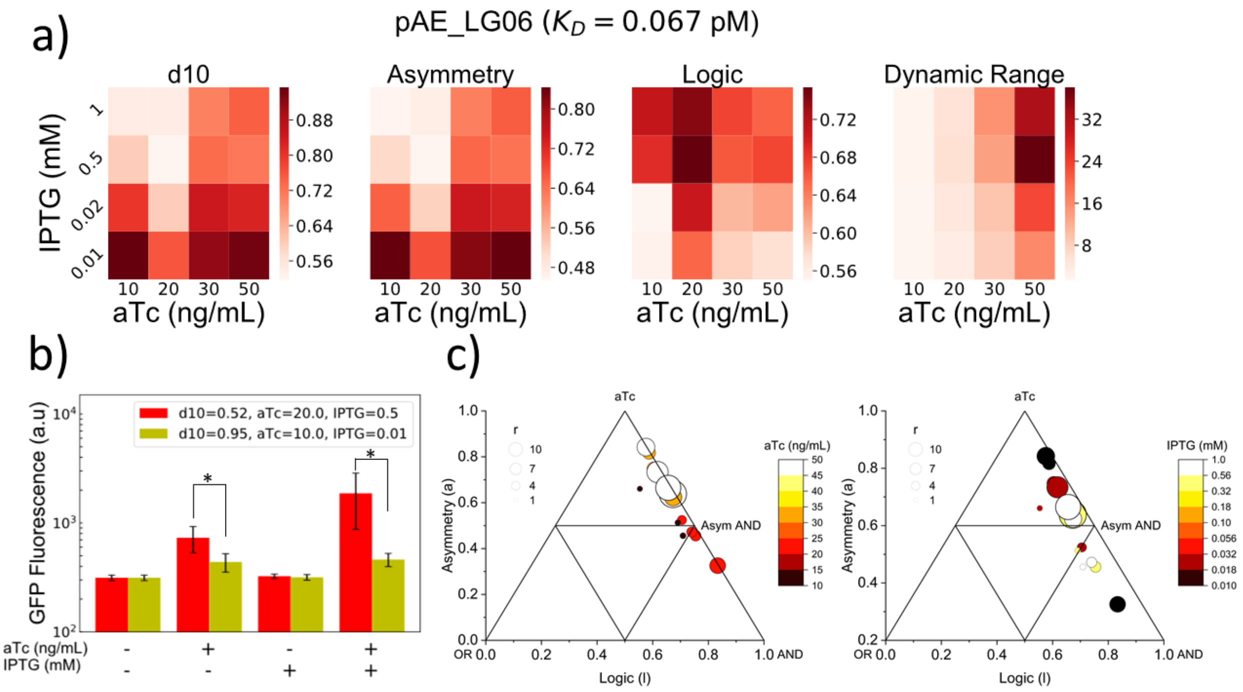

**Supplementary Figure S8:**

a) Heat maps of logic parameters of pAE_LG06. b) Bar plot of minimum and maximum d10 conditions for pAE_LG06. c) Triangle plots of pAE_LG06 at varying aTc and IPTG conditions, showing trends in aTc (left) and IPTG (right).

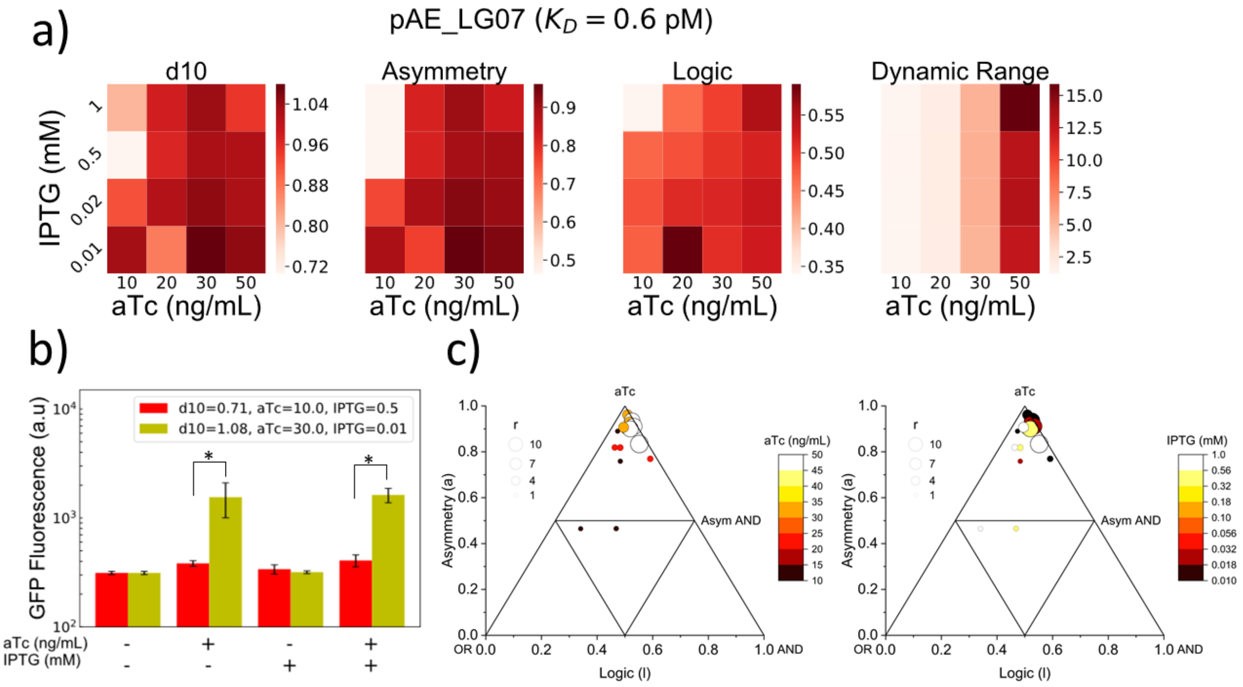

**Supplementary Figure S9:**

a) Heat maps of logic parameters of pAE_LG07. b) Bar plot of minimum and maximum d10 conditions for pAE_LG07. c) Triangle plots of pAE_LG07 at varying aTc and IPTG conditions, showing trends in aTc (left) and IPTG (right).

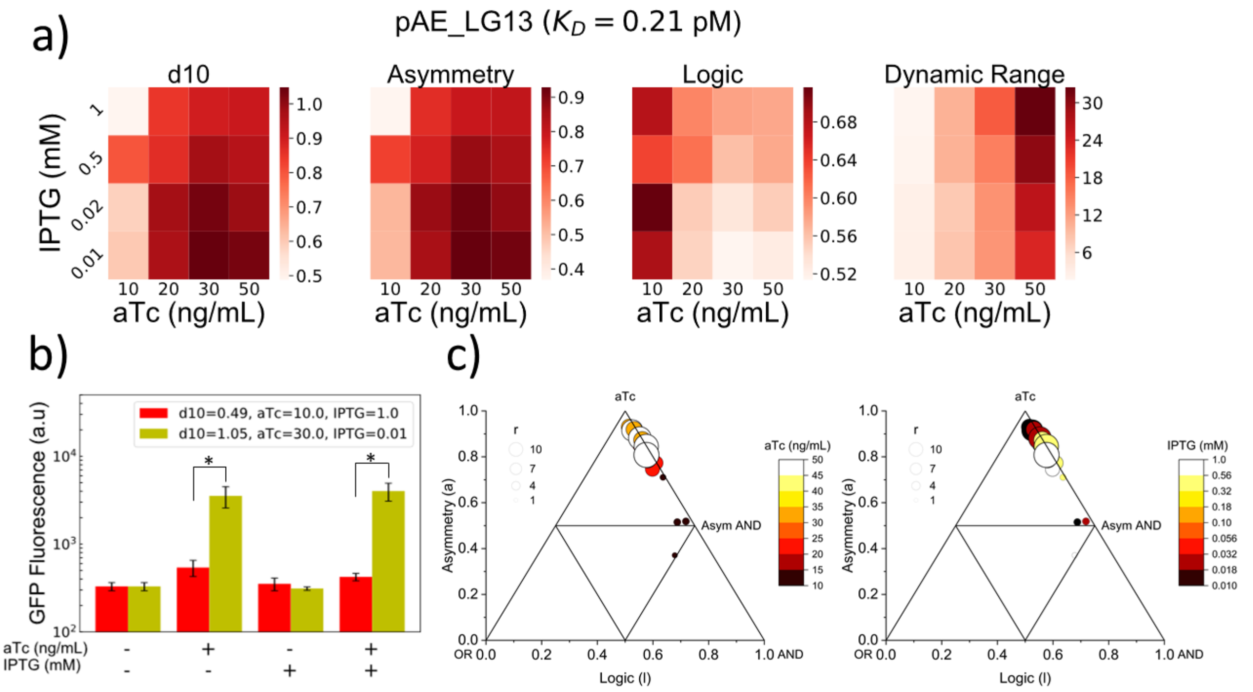

**Supplementary Figure S10:**

a) Heat maps of logic parameters of pAE_LG13. b) Bar plot of minimum and maximum d10 conditions for pAE_LG13. c) Triangle plots of pAE_LG13 at varying aTc and IPTG conditions, showing trends in aTc (left) and IPTG (right).

### AND Gate Model Fits

**pAE_LG01**

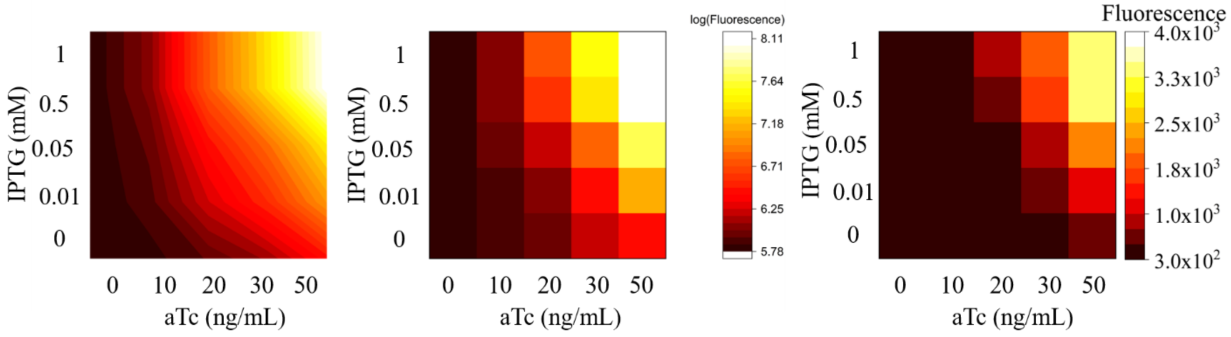

| | value | +/- |
|---|---|---|
| K_aTcbind | 1.00 | (at bound) |
| K_LacIbind | 0.22 | 0.15 |
| K_RNAPbind | 0.00 | 0.00 |
| k_translation | 12.00 | (at bound) |
| m_TetRHill | 1.79 | 0.10 |
| n_LacIHill | 1.34 | 0.14 |
| a_Readthrough | - | - |

| | |
|---|---|
| AIC | -104.90 |
| BIC | -100.03 |
| RMSE | 0.11 |
| $R^2$ | 0.98 |

| IPTG | Flow Cytometry Std. Deviation (n>=4) | | | | |
|---|---|---|---|---|---|
| 1 | 22 | 55.9 | 351 | 806 | 731 |
| 0.5 | 21.5 | 69.2 | 220 | 444 | 816 |
| 0.02 | 21.6 | 37.2 | 106 | 233 | 691 |
| 0.01 | 21.6 | 4.58 | 55.9 | 97.4 | 254 |
| 0 | 23.1 | 3.74 | 11.4 | 98.2 | 177 |
| aTc | 0 | 10 | 20 | 30 | 50 |

**Supplementary Figure S11:**

Heat maps of (from left to right) fitted log-transformed, log-transformed, and raw GFP expression data are reported, along with fitted parameter values, goodness of fit statistics, and standard deviations from biological replicates for AND gate pAE_LG01.

### pAE_LG02

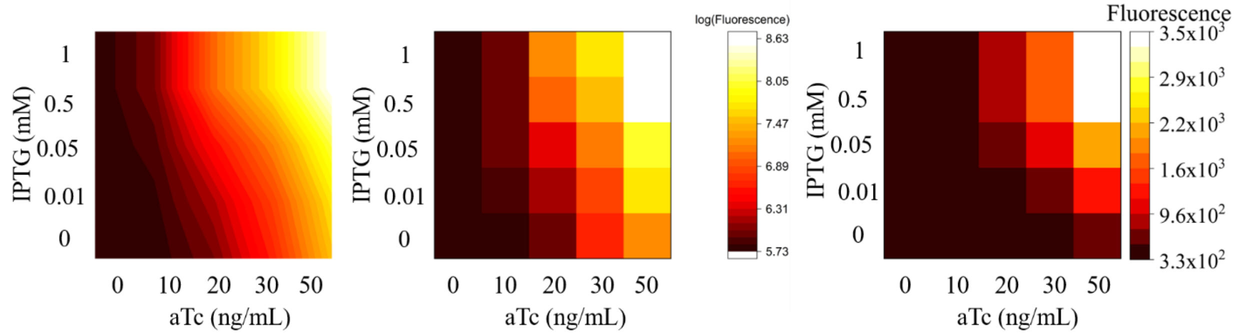

| | value | +/- |
|---|---|---|
| K_aTcbind | 0.14 | 0.29 |
| K_LacIbind | 3.42 | 7.48 |
| K_RNAPbind | 0.00 | 0.04 |
| k_translation | 3.58 | 3.47 |
| m_TetRHill | 4.00 | 3.23 |
| n_LacIHill | 2.24 | 0.89 |
| a_Readthrough | 0.33 | 0.07 |

| | |
|---|---|
| AIC | -110.29 |
| BIC | -101.76 |
| RMSE | 0.10 |
| $R^2$ | 0.99 |

| IPTG | **Flow Cytometry Std. Deviation (n>=4)** | | | | |
|---|---|---|---|---|---|
| 1 | 20 | 51 | 714 | 769 | 1979 |
| 0.5 | 21 | 64 | 689 | 623 | 1572 |
| 0.02 | 22 | 40 | 163 | 270 | 634 |
| 0.01 | 31 | 21 | 73 | 127 | 375 |
| 0 | 21 | 39 | 39 | 271 | 296 |
| aTc | 0 | 10 | 20 | 30 | 50 |

**Supplementary Figure S12:**

Heat maps of (from left to right) fitted log-transformed, log-transformed, and raw GFP expression data are reported, along with fitted parameter values, goodness of fit statistics, and standard deviations from biological replicates for AND gate pAE_LG02.

### pAE_LG03

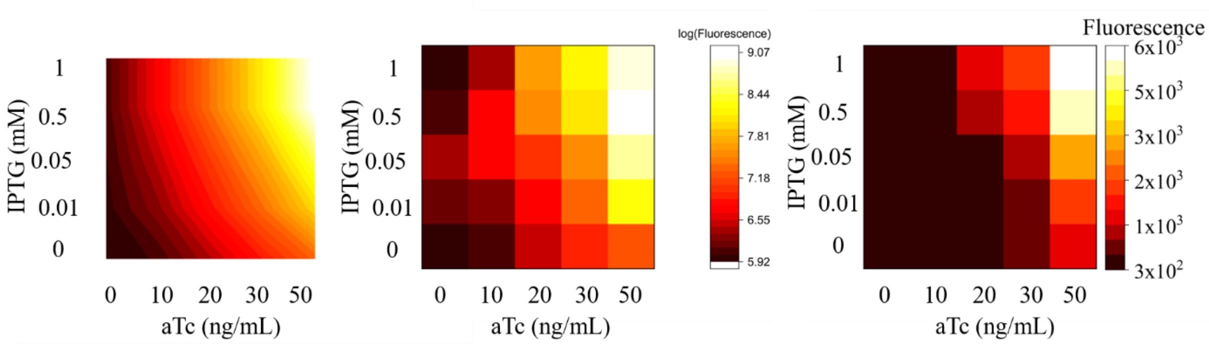

| | value | +/- |
|---|---|---|
| K_aTcbind | 0.06 | 0.09 |
| K_LacIbind | 0.33 | 0.63 |
| K_RNAPbind | 1.56 | 6.00 |
| k_translation | 12.00 | 34.38 |
| m_TetRHill | 0.54 | 0.49 |
| n_LacIHill | 1.63 | 0.84 |
| a_Readthrough | 0.87 | 0.37 |

| | |
|---|---|
| AIC | -80.31 |
| BIC | -71.78 |
| RMSE | 0.18 |
| $R^2$ | 0.97 |

| IPTG | Flow Cytometry Std. Deviation (n>=4) | | | | |
|---|---|---|---|---|---|
| 1 | 41.3 | 77.3 | 786 | 1142 | 1487 |
| 0.5 | 122 | 223 | 844 | 1004 | 4676 |
| 0.02 | 183 | 39.5 | 388 | 231 | 1937 |
| 0.01 | 120 | 110 | 235 | 595 | 1651 |
| 0 | 63.6 | 62.1 | 117 | 144 | 57.2 |
| aTc | 0 | 10 | 20 | 30 | 50 |

**Supplementary Figure S13:**

Heat maps of (from left to right) fitted log-transformed, log-transformed, and raw GFP expression data are reported, along with fitted parameter values, goodness of fit statistics, and standard deviations from biological replicates for AND gate pAE_LG03.

### pAE_LG04

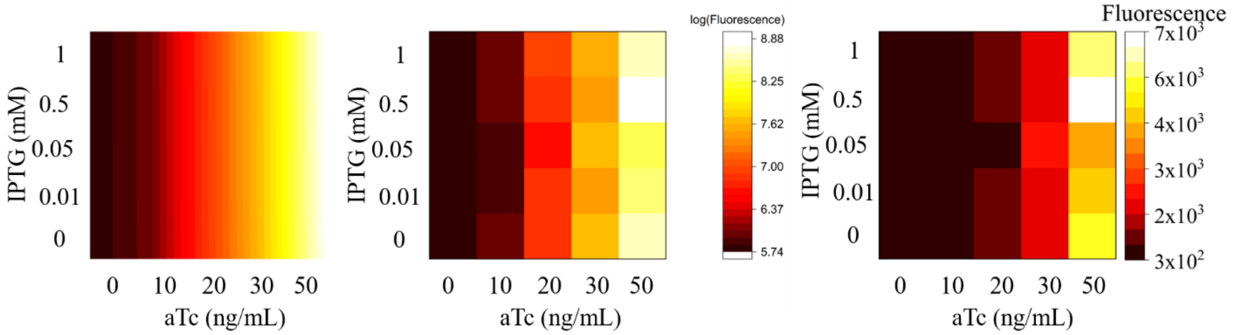

| | value | +/- |
|---|---|---|
| K_aTcbind | 0.82 | 0.52 |
| K_LacIbind | 0.00 | At bound |
| K_RNAPbind | 0.00 | At bound |
| k_translation | 5.51 | -4.61 |
| m_TetRHill | 1.51 | 0.30 |
| n_LacIHill | 0.26 | 8.26 |
| a_Readthrough | - | - |

| | |
|---|---|
| AIC | -110.84 |
| BIC | -105.96 |
| RMSE | 0.10 |
| $R^2$ | 0.99 |

| IPTG | Flow Cytometry Std. Deviation (n>=4) | | | | |
|---|---|---|---|---|---|
| 1 | 24 | 28 | 536 | 547 | 1499 |
| 0.5 | 27 | 47 | 332 | 431 | 3659 |
| 0.02 | 27 | 44 | 158 | 1189 | 736 |
| 0.01 | 26 | 29 | 439 | 254 | 775 |
| 0 | 26 | 33 | 246 | 462 | 1073 |
| aTc | 0 | 10 | 20 | 30 | 50 |

**Supplementary Figure S14:**

Heat maps of (from left to right) fitted log-transformed, log-transformed, and raw GFP expression data are reported, along with fitted parameter values, goodness of fit statistics, and standard deviations from biological replicates for AND gate pAE_LG04.

### pAE_LG05

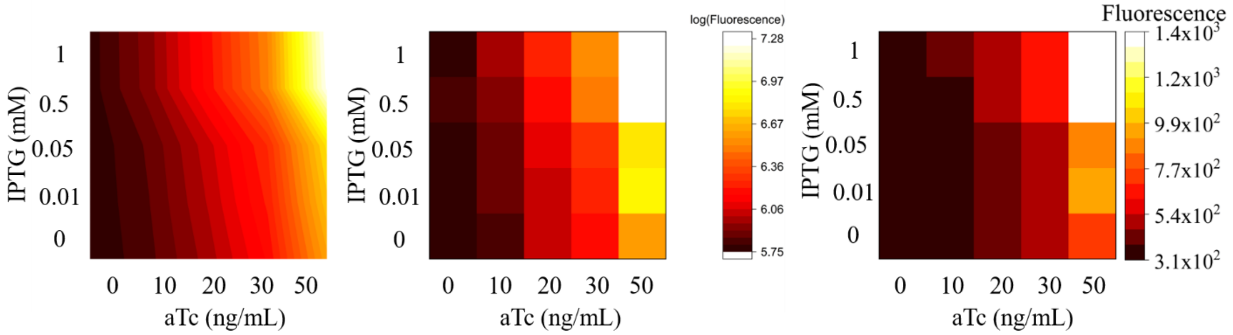

| | value | +/- |
|---|---|---|
| K_aTcbind | 0.05 | 0.03 |
| K_LacIbind | 0.43 | 0.32 |
| K_RNAPbind | 0.87 | 1.12 |
| k_translation | 2.83 | 2.62 |
| m_TetRHill | 0.98 | 0.17 |
| n_LacIHill | 1.06 | 0.26 |
| a_Readthrough | - | - |

| | |
|---|---|
| AIC | -180.27 |
| BIC | -172.95 |
| RMSE | 0.02 |
| $R^2$ | 1.00 |

| IPTG | Flow Cytometry Std. Deviation (n>=4) | | | | |
|---|---|---|---|---|---|
| 1 | 35 | 55 | 94 | 141 | 499 |
| 0.5 | 42 | 43 | 88 | 105 | 501 |
| 0.02 | 32 | 32 | 62 | 44 | 153 |
| 0.01 | 29 | 28 | 57 | 53 | 260 |
| 0 | 28 | 25 | 49 | 49 | 174 |
| aTc | 0 | 10 | 20 | 30 | 50 |

**Supplementary Figure S15:**

Heat maps of (from left to right) fitted log-transformed, log-transformed, and raw GFP expression data are reported, along with fitted parameter values, goodness of fit statistics, and standard deviations from biological replicates for AND gate pAE_LG05.

### pAE_LG06

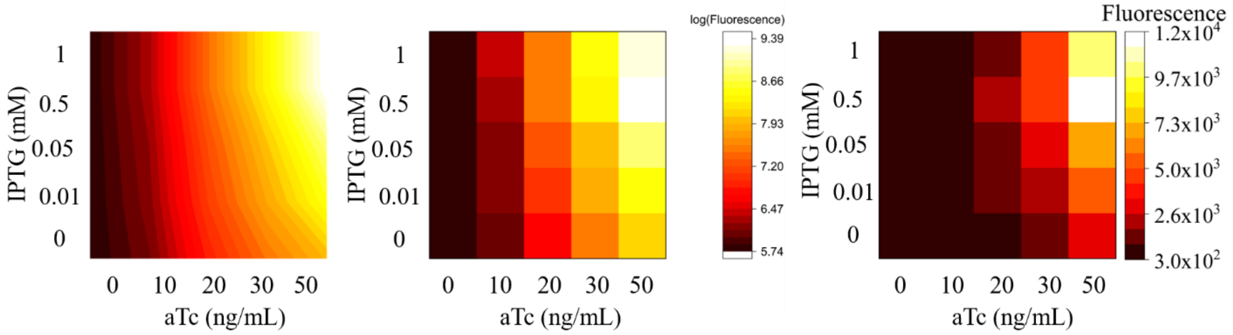

| | value | +/- |
|---|---|---|
| K_aTcbind | 0.12 | 0.11 |
| K_LacIbind | 0.20 | 0.16 |
| K_RNAPbind | 0.88 | 1.46 |
| k_translation | 2.39 | 0.18 |
| m_TetRHill | 1.02 | 0.10 |
| n_LacIHill | 1.39 | 0.29 |
| a_Readthrough | - | - |

| | |
|---|---|
| AIC | -126.98 |
| BIC | -119.67 |
| RMSE | 0.07 |
| $R^2$ | 1.00 |

| IPTG | Flow Cytometry Std. Deviation (n>=4) | | | | |
|---|---|---|---|---|---|
| 1 | 19 | 253 | 872 | 2340 | 3389 |
| 0.5 | 15 | 157 | 996 | 2193 | 5833 |
| 0.02 | 25 | 61 | 646 | 1339 | 3720 |
| 0.01 | 19 | 64 | 476 | 1014 | 2290 |
| 0 | 18 | 84 | 197 | 538 | 437 |
| aTc | 0 | 10 | 20 | 30 | 50 |

**Supplementary Figure S16:**

Heat maps of (from left to right) fitted log-transformed, log-transformed, and raw GFP expression data are reported, along with fitted parameter values, goodness of fit statistics, and standard deviations from biological replicates for AND gate pAE_LG06.

### pAE_LG07

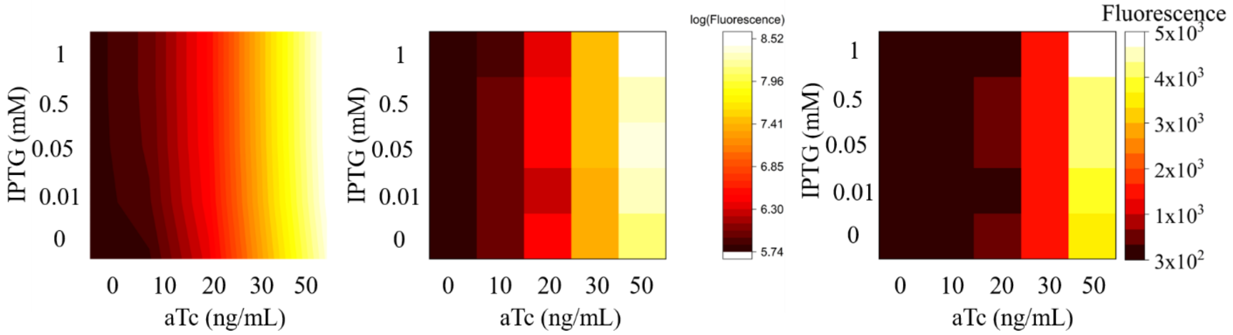

| | value | +/- |
|---|---|---|
| K_aTcbind | 0.04 | 0.09 |
| K_LacIbind | 0.44 | 0.63 |
| K_RNAPbind | 1.42 | 6.00 |
| k_translation | 2.64 | 34.38 |
| m_TetRHill | 0.01 | 0.49 |
| n_LacIHill | 2.34 | 0.84 |
| a_Readthrough | - | - |

| | |
|---|---|
| AIC | -129.91 |
| BIC | -123.81 |
| RMSE | 0.07 |
| $R^2$ | 1.00 |

| IPTG | Flow Cytometry Std. Deviation (n>=4) | | | | |
|---|---|---|---|---|---|
| 1 | 35 | 14 | 225 | 360 | 1195 |
| 0.5 | 33 | 51 | 310 | 644 | 797 |
| 0.02 | 11 | 35 | 179 | 480 | 175 |
| 0.01 | 9 | 20 | 89 | 245 | 133 |
| 0 | 10 | 22 | 161 | 548 | 175 |
| aTc | 0 | 10 | 20 | 30 | 50 |

**Supplementary Figure S17:**

Heat maps of (from left to right) fitted log-transformed, log-transformed, and raw GFP expression data are reported, along with fitted parameter values, goodness of fit statistics, and standard deviations from biological replicates for AND gate pAE_LG07.

### pAE_LG13

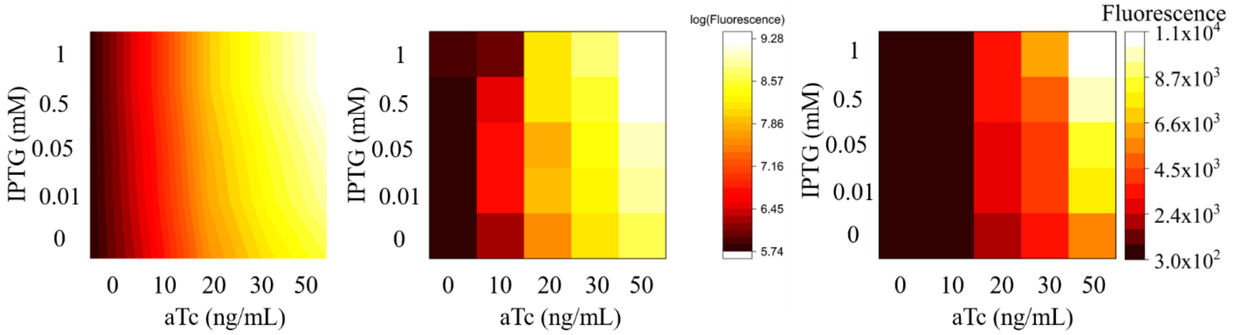

| | value | +/- |
|---|---|---|
| K_aTcbind | 0.17 | 0.09 |
| K_LacIbind | 0.12 | 0.63 |
| K_RNAPbind | 1.07 | 6.00 |
| k_translation | 3.00 | 34.38 |
| m_TetRHill | 0.54 | 0.49 |
| n_LacIHill | 1.23 | 0.84 |
| a_Readthrough | - | - |

| | |
|---|---|
| AIC | -96.53 |
| BIC | -89.21 |
| RMSE | 0.13 |
| $R^2$ | 0.99 |

| IPTG | Flow Cytometry Std. Deviation (n>=4) | | | | |
|---|---|---|---|---|---|
| 1 | 58 | 40 | 1249 | 755 | 3372 |
| 0.5 | 31 | 325 | 1139 | 1694 | 2276 |
| 0.02 | 20 | 459 | 549 | 1074 | 1850 |
| 0.01 | 14 | 438 | 765 | 925 | 1666 |
| 0 | 35 | 114 | 330 | 959 | 1046 |
| aTc | 0 | 10 | 20 | 30 | 50 |

**Supplementary Figure S18:**

Heat maps of (from left to right) fitted log-transformed, log-transformed, and raw GFP expression data are reported, along with fitted parameter values, goodness of fit statistics, and standard deviations from biological replicates for AND gate pAE_LG13.

**OR Gate Logic Behaviors**

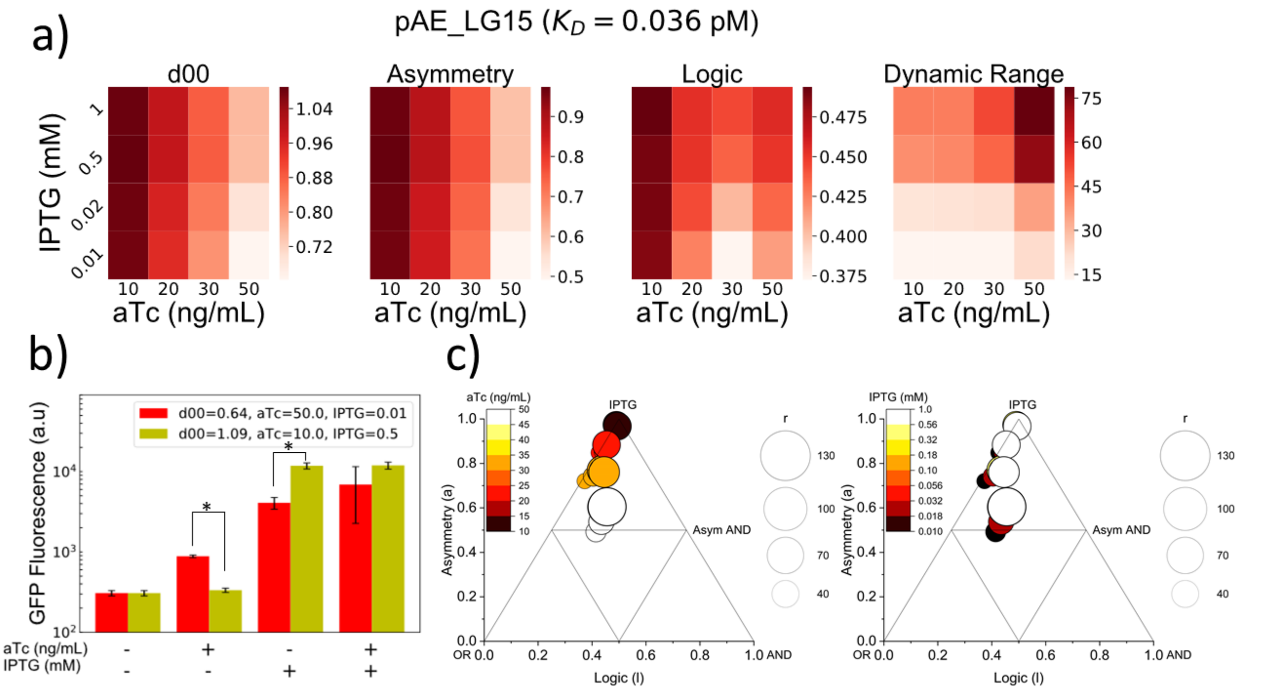

**Supplementary Figure S19:**

a) Heat maps of logic parameters of pAE_LG15. b) Bar plot of minimum and maximum d10 conditions for pAE_LG15. c) Triangle plots of pAE_LG15 at varying aTc and IPTG conditions, showing trends in aTc (left) and IPTG (right).

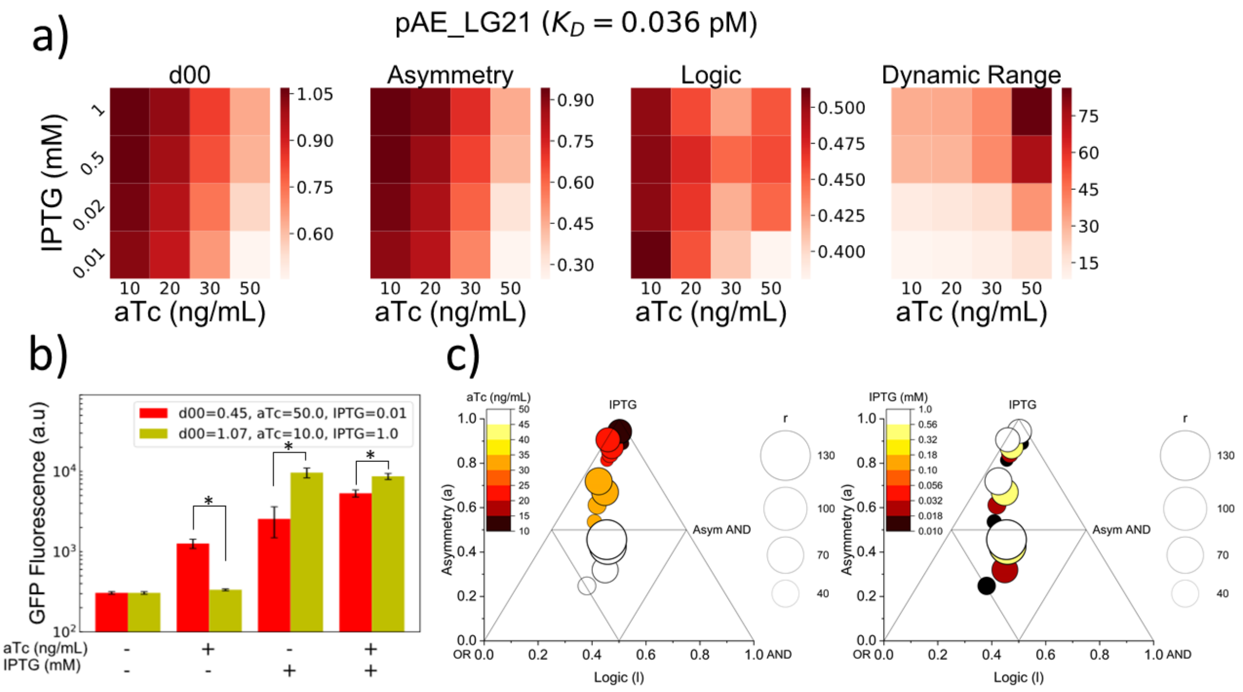

**Supplementary Figure S20:**

a) Heat maps of logic parameters of pAE_LG21. b) Bar plot of minimum and maximum d10 conditions for pAE_LG21. c) Triangle plots of pAE_LG21 at varying aTc and IPTG conditions, showing trends in aTc (left) and IPTG (right).

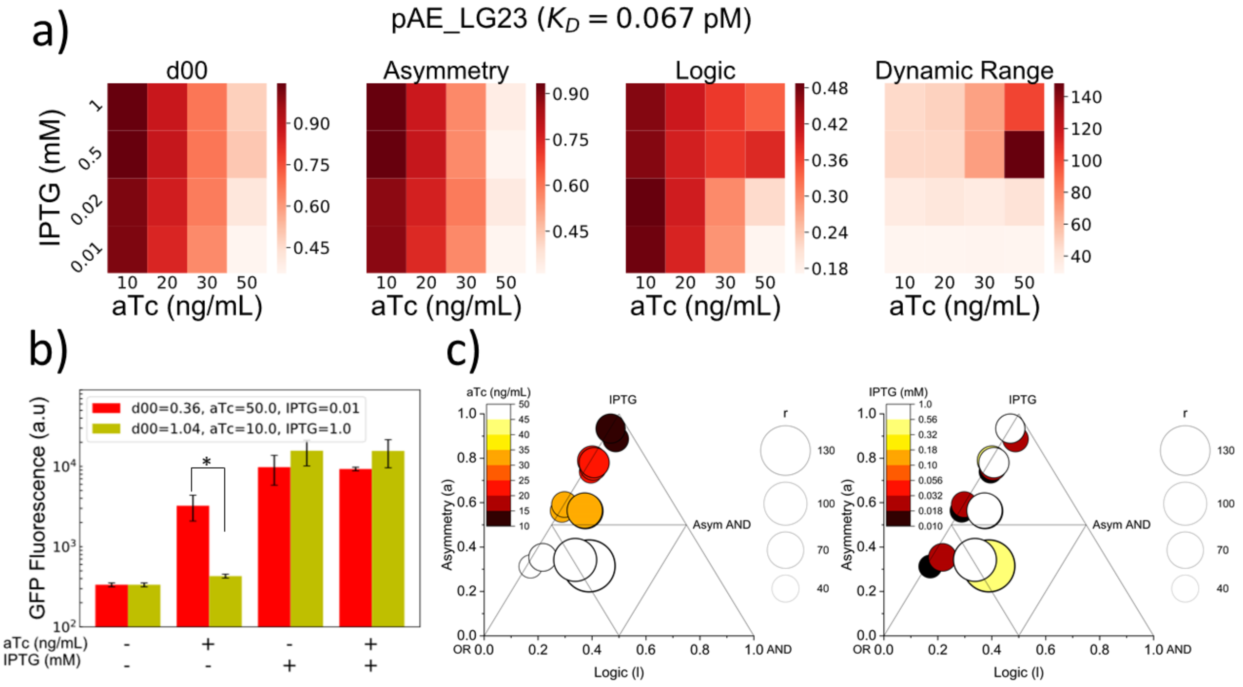

**Supplementary Figure S21:**

a) Heat maps of logic parameters of pAE_LG23. b) Bar plot of minimum and maximum d10 conditions for pAE_LG23. c) Triangle plots of pAE_LG23 at varying aTc and IPTG conditions, showing trends in aTc (left) and IPTG (right).

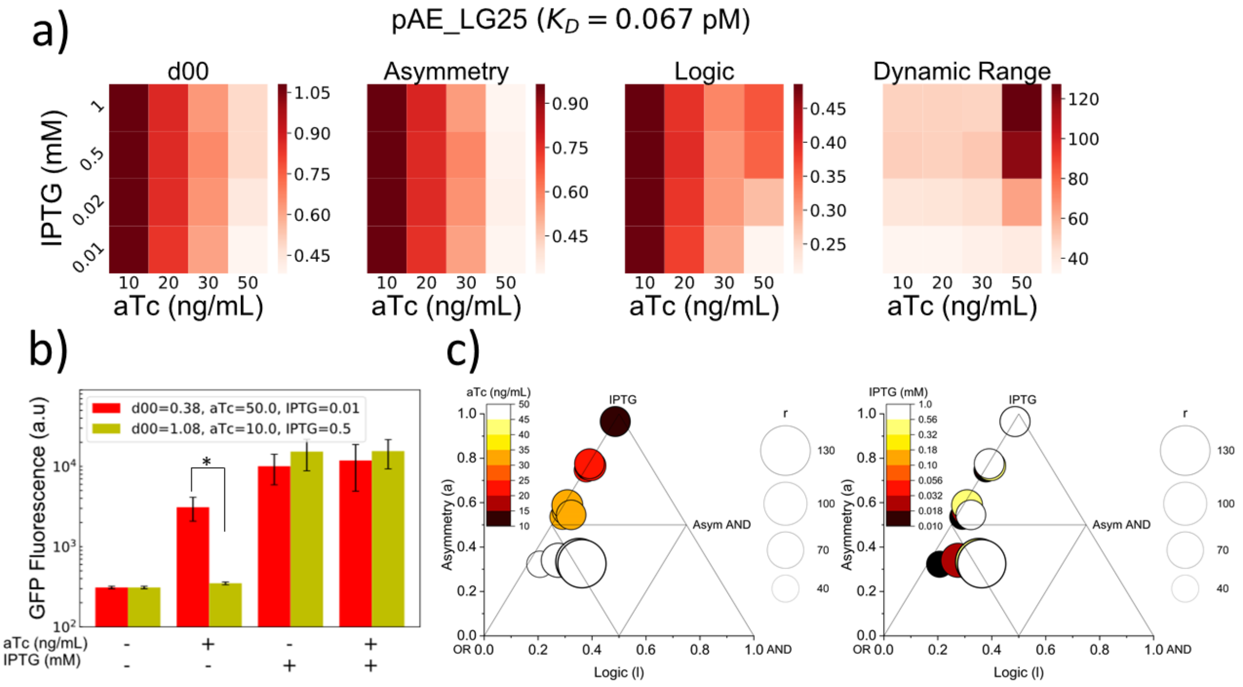

**Supplementary Figure S22:**

a) Heat maps of logic parameters of pAE_LG25. b) Bar plot of minimum and maximum d10 conditions for pAE_LG25. c) Triangle plots of pAE_LG25 at varying aTc and IPTG conditions, showing trends in aTc (left) and IPTG (right).

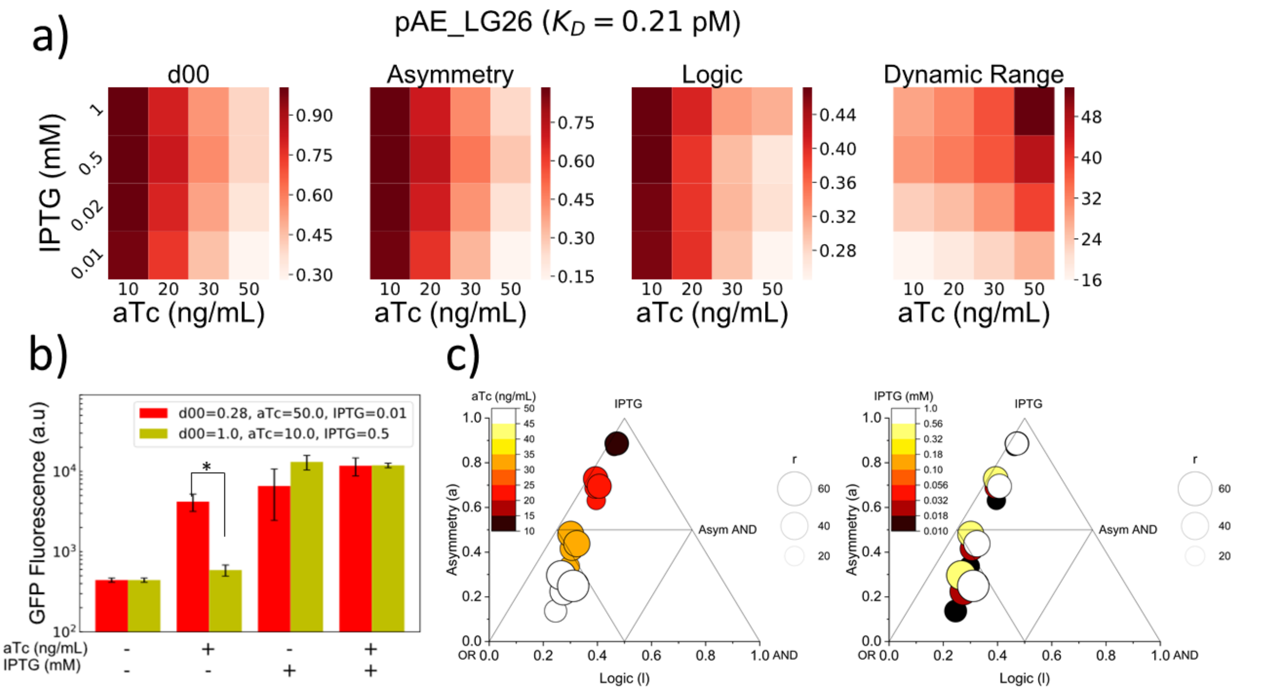

**Supplementary Figure S23:**

a) Heat maps of logic parameters of pAE_LG26. b) Bar plot of minimum and maximum d10 conditions for pAE_LG26. c) Triangle plots of pAE_LG26 at varying aTc and IPTG conditions, showing trends in aTc (left) and IPTG (right).

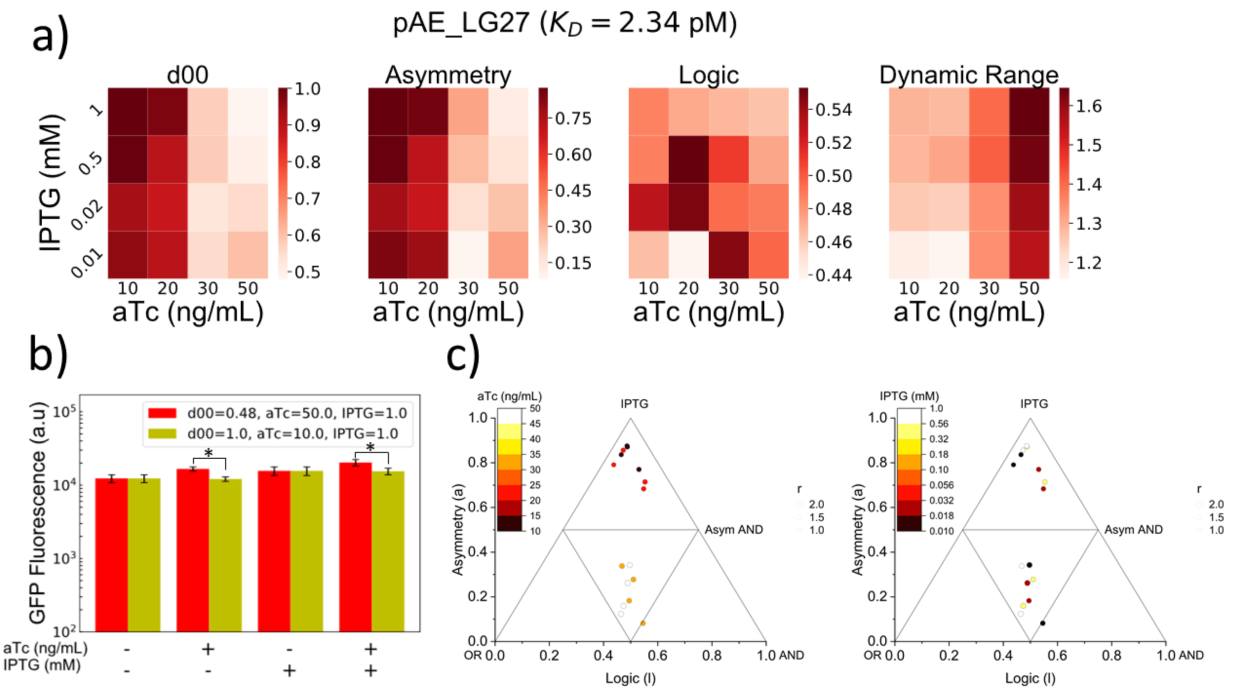

**Supplementary Figure S24:**

a) Heat maps of logic parameters of pAE_LG27. b) Bar plot of minimum and maximum d10 conditions for pAE_LG27. c) Triangle plots of pAE_LG27 at varying aTc and IPTG conditions, showing trends in aTc (left) and IPTG (right).

### OR Gate Model Fits

#### pAE_LG15

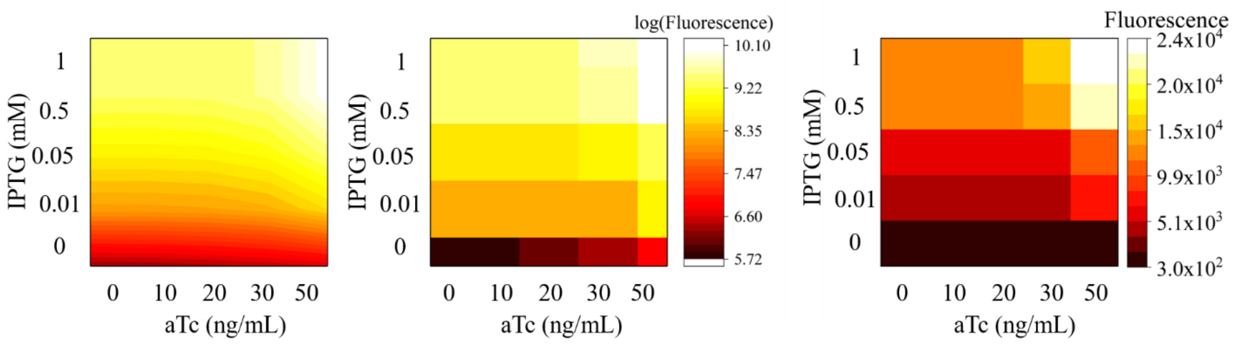

| | value | +/- |
|---|---|---|
| RNAPbind2 | 6.15E-04 | 1.20E-03 |
| a1 | 0.03 | 0.11 |
| a2 | 0.55 | 0.38 |
| m | 1.95 | 8.69 |
| n | 1.68 | 0.61 |
| b | 1.90E-04 | 0.00 |
| crit | 75.83 | 78.07 |

| | |
|---|---|
| AIC | -68.13 |
| BIC | -59.60 |
| $R^2$ | 0.98 |
| RMSE | 0.23 |

| IPTG | Flow Cytometry Std. Deviation (n>=4) | | | | |
|---|---|---|---|---|---|
| 1 | 998 | 1045 | 884 | 1507 | 1366 |
| 0.5 | 1011 | 1188 | 1101 | 1392 | 2353 |
| 0.02 | 611 | 775 | 722 | 541 | 1003 |
| 0.01 | 684 | 528 | 117 | 454 | 4627 |
| 0 | 23 | 21 | 102 | 80 | 33 |
| aTc | 0 | 10 | 20 | 30 | 50 |

**Supplementary Figure S25:**

Heat maps of (from left to right) fitted log-transformed, log-transformed, and raw GFP expression data are reported, along with fitted parameter values, goodness of fit statistics, and standard deviations from biological replicates for OR gate pAE_LG15.

### pAE_LG21

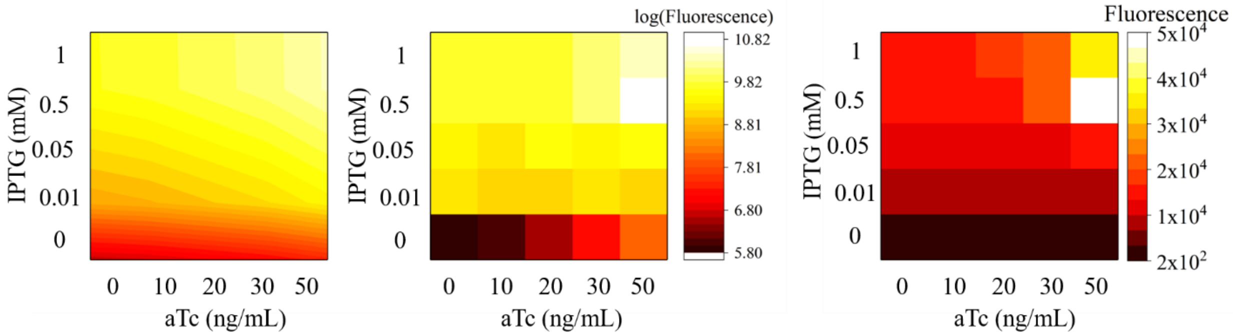

| | value | +/- |
|---|---|---|
| RNAPbind2 | 5.39E-04 | 1.10E-03 |
| a1 | 0.03 | 0.06 |
| a2 | 0.51 | 0.41 |
| m | 2.53 | 6.25 |
| n | 1.59 | 0.59 |
| b | 2.54E-04 | 0.00 |
| crit | 50.35 | 81.18 |

| | |
|---|---|
| AIC | -69.33 |
| BIC | -60.80 |
| $R^2$ | 0.98 |
| RMSE | 0.22 |

| IPTG | Flow Cytometry Std. Deviation (n>=4) | | | | |
|---|---|---|---|---|---|
| 1 | 1355 | 744 | 683 | 780 | 7573 |
| 0.5 | 1087 | 1907 | 656 | 1213 | 6382 |
| 0.02 | 1072 | 741 | 755 | 432 | 2534 |
| 0.01 | 1071 | 258 | 764 | 1368 | 537 |
| 0 | 12 | 9 | 2 | 115 | 166 |
| aTc | 0 | 10 | 20 | 30 | 50 |

**Supplementary Figure S26:**

Heat maps of (from left to right) fitted log-transformed, log-transformed, and raw GFP expression data are reported, along with fitted parameter values, goodness of fit statistics, and standard deviations from biological replicates for OR gate pAE_LG21.

### pAE_LG23

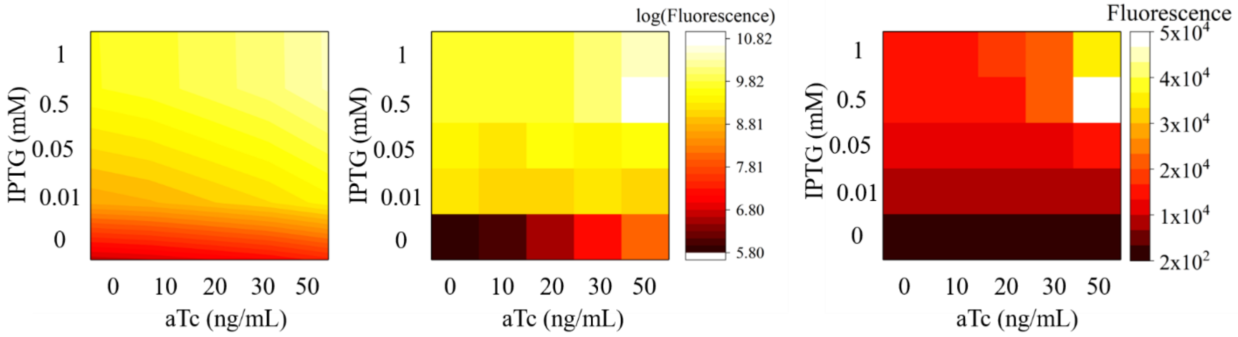

| | value | +/- |
|---|---|---|
| RNAPbind2 | 1.01E-03 | 5.95E-04 |
| a1 | 0.12 | 0.06 |
| a2 | 0.50 | 0.09 |
| m | | |
| n | | |
| b | | |
| crit | | |

| | |
|---|---|
| AIC | -49.36 |
| BIC | -45.70 |
| $R^2$ | 0.93 |
| RMSE | 0.35 |

| IPTG | Flow Cytometry Std. Deviation (n>=4) | | | | |
|---|---|---|---|---|---|
| 1 | 5552 | 5950 | 5400 | 11490 | 24286 |
| 0.5 | 5968 | 7136 | 9747 | 12149 | 40875 |
| 0.02 | 4718 | 4964 | 5632 | 3943 | 2599 |
| 0.01 | 3939 | 3037 | 5278 | 5933 | 481 |
| 0 | 19 | 24 | 189 | 385 | 1148 |
| aTc | 0 | 10 | 20 | 30 | 50 |

**Supplementary Figure S27:**

Heat maps of (from left to right) fitted log-transformed, log-transformed, and raw GFP expression data are reported, along with fitted parameter values, goodness of fit statistics, and standard deviations from biological replicates for OR gate pAE_LG23.

### pAE_LG25

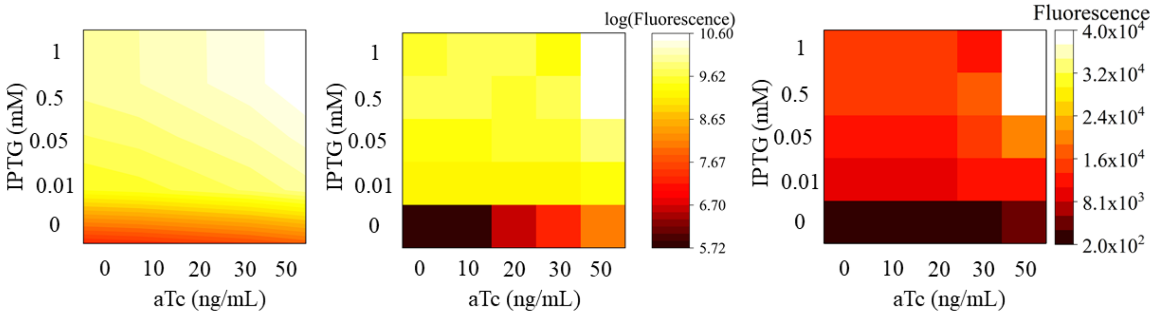

| | value | +/- |
|---|---|---|
| K_aTcbind | 0.12 | 0.11 |
| K_LacIbind | 0.20 | 0.16 |
| K_RNAPbind | 0.88 | 1.46 |
| k | 2.39 | 0.18 |
| m | 1.02 | 0.10 |
| n | 1.39 | 0.29 |
| RNAPbind2 | 2.55E-03 | 1.15E-03 |

| | |
|---|---|
| AIC | -55.97 |
| BIC | -52.31 |
| $R^2$ | 0.95 |
| RMSE | 0.31 |

| IPTG | **Flow Cytometry Std. Deviation (n>=4)** | | | | |
|---|---|---|---|---|---|
| 1 | 7408 | 6157 | 3883 | 258 | 20269 |
| 0.5 | 6505 | 6169 | 4218 | 6115 | 18166 |
| 0.02 | 5936 | 5504 | 4984 | 4458 | 8327 |
| 0.01 | 4155 | 3838 | 3552 | 3765 | 6932 |
| 0 | 11 | 14 | 176 | 439 | 1018 |
| aTc | 0 | 10 | 20 | 30 | 50 |

**Supplementary Figure S28:**

Heat maps of (from left to right) fitted log-transformed, log-transformed, and raw GFP expression data are reported, along with fitted parameter values, goodness of fit statistics, and standard deviations from biological replicates for OR gate pAE_LG25.

### pAE_LG26

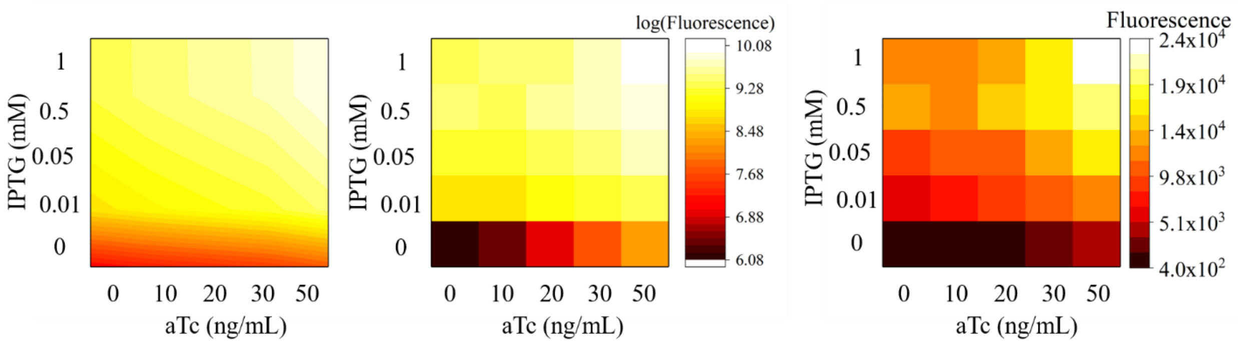

| | value | +/- |
|---|---|---|
| RNAPbind2 | 1.58E-03 | 4.57E-04 |
| a1 | 1.83 | 0.59 |
| a2 | 0.41 | 0.03 |

| | |
|---|---|
| AIC | -84.30 |
| BIC | -80.64 |
| RMSE | 0.18 |
| $R^2$ | 0.98 |

| IPTG | Flow Cytometry Std. Deviation (n>=4) | | | | |
|---|---|---|---|---|---|
| 1 | 707 | 1001 | 696 | 941 | 3978 |
| 0.5 | 2697 | 768 | 3126 | 1082 | 1288 |
| 0.02 | 1774 | 2070 | 2139 | 2837 | 3134 |
| 0.01 | 4120 | 2780 | 2773 | 3262 | 2982 |
| 0 | 28 | 93 | 188 | 542 | 1022 |
| aTc | 0 | 10 | 20 | 30 | 50 |

**Supplementary Figure S29:**

Heat maps of (from left to right) fitted log-transformed, log-transformed, and raw GFP expression data are reported, along with fitted parameter values, goodness of fit statistics, and standard deviations from biological replicates for OR gate pAE_LG26.

### pAE_LG27

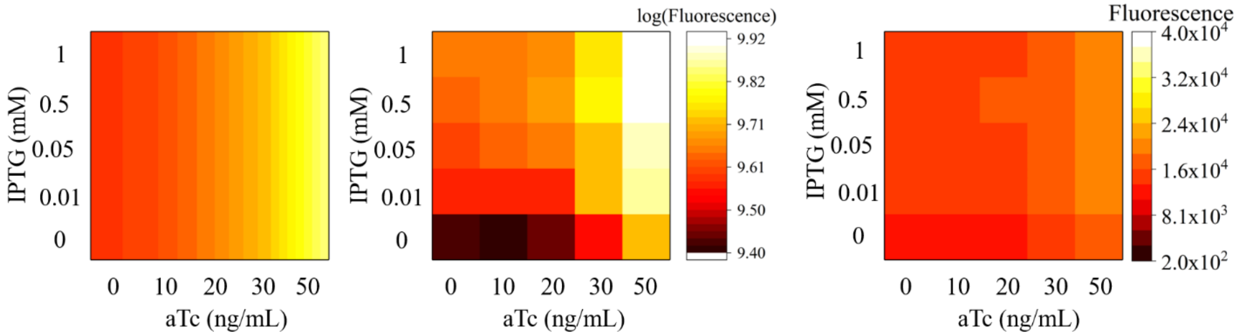

| | value | +/- |
|---|---|---|
| RNAPbind2 | 4.94 | 3.73E+06 |
| a1 | 0.04 | 0.14 |
| a2 | 0.37 | 18.36 |
| m | 1.34 | 2.65 |
| n | 2 | 1.10E+08 |

| | |
|---|---|
| AIC | -112.75 |
| BIC | -106.65 |
| $R^2$ | 0.61 |
| RMSE | 0.10 |

| IPTG | Flow Cytometry Std. Deviation (n>=4) | | | | |
|---|---|---|---|---|---|
| 1 | 2114 | 1584 | 1191 | 1228 | 2022 |
| 0.5 | 1535 | 1541 | 1872 | 1785 | 2288 |
| 0.02 | 1394 | 1448 | 1617 | 1409 | 1628 |
| 0.01 | 1355 | 1131 | 1463 | 1771 | 1565 |
| 0 | 1502 | 823 | 1006 | 1202 | 1148 |
| aTc | 0 | 10 | 20 | 30 | 50 |

**Supplementary Figure S30:**

Heat maps of (from left to right) fitted log-transformed, log-transformed, and raw GFP expression data are reported, along with fitted parameter values, goodness of fit statistics, and standard deviations from biological replicates for OR gate pAE_LG27.

### Bibliography

Betz JL, Sasmor HM, Buck F, Insley MY & Caruthers MH (1986) Base substitution mutants of the lac operator: in vivo and in vitro affinities for lac repressor. *Gene* **50:** 123–132 Available at: http://linkinghub.elsevier.com/retrieve/pii/0378111986903173

Bozdogan H (1987) Model selection and Akaike's Information Criterion (AIC): The general theory and its analytical extensions. *Psychometrika* **52:** 345–370

Chens J, Albertil S & Matthewsm KS (1994) Wild-type Operator Binding and Altered Cooperativity for Inducer Binding of. **269:** 12482–12487

Cox RS, Surette MG & Elowitz MB (2007) Programming gene expression with combinatorial promoters. *Mol. Syst. Biol.* **3:** 145 Available at: http://www.nature.com/doifinder/10.1038/msb4100187

Kamionka A, Bogdanska-urbaniak J, Scholz O, Hillen W, Mikrobiologie È & Genetik B (2004) Two mutations in the tetracycline repressor change the inducer anhydrotetracycline to a corepressor. **32:** 842–847

Munro PD, Ackers GK & Shearwin KE (2016) Aspects of protein–DNA interactions: a review of quantitative thermodynamic theory for modelling synthetic circuits utilising LacI and CI repressors, IPTG and the reporter gene lacZ. *Biophys. Rev.* **8:** 331–345 Available at: http://dx.doi.org/10.1007/s12551-016-0231-9

Shea MA & Ackers GK (1985) The OR control system of bacteriophage lambda. A physical-chemical model for gene regulation. *J. Mol. Biol.* **181:** 211–230

Tamsir A, Tabor JJ & Voigt CA (2011) Robust multicellular computing using genetically encoded NOR gates and chemical 'wires'. *Nature* **469:** 212–215 Available at: http://www.pubmedcentral.nih.gov/articlerender.fcgi?artid=3904220&tool=pmcentrez&rendertype=abstract [Accessed July 14, 2014]
